# Conserved “late” effector genes from *Leptosphaeria maculans* inducing gene-for-gene quantitative resistance in *Brassica napus* semi-winter genotypes

**DOI:** 10.1101/2025.09.22.677120

**Authors:** Camille Rabeau, Armand Wagner, Nicolas Lapalu, Audren Jiquel, Sébastien Faure, Isabelle Fudal

## Abstract

*Leptosphaeria maculans* is a phytopathogenic fungus responsible for stem canker on *Brassica napus*. Its infectious cycle goes through an early phase of leaf infection and a late phase of colonization and infection of the stem. The disease is mainly controlled by plant genetic resistances targeting a limited set of early fungal effector genes overexpressed during leaf infection and located in dynamic repeat-rich genomic regions. Thus, these resistances can be rapidly overcome by the pathogen. To find new sources of resistance, we focused on late effector genes, expressed during stem infection and located in gene-rich regions. A previous study revealed a quantitative resistance in the stem partly relying on a gene-for-gene interaction with a late effector gene. In this study, we deciphered whether all late effector genes shared the same genomic and evolutionary characteristics and if they could be more stable than early effector genes, rendering the resistances they trigger more durable. In addition, as previous studies highlighted new criteria for selecting late effectors and suggested *B. napus* semi-winter genotypes as an interesting genetic pool for uncovering resistance sources, we selected six new late effector gene candidates and screened an enlarged panel of semi-winter genotypes. We revealed that early and late effector genes diverged for most of their genomic characteristics supplying for the hypothesis of late effector genes being more conserved. Moreover, we revealed new resistance sources to late effector genes, almost all belonging to the semi-winter genetic pool, validating their importance to uncover new resistance sources.

## INTRODUCTION

During plant infection, fungal pathogens secrete effectors, mainly small secreted proteins, that modulate plant immunity and facilitate infection. Some of them are recognized by the plant immune system and termed avirulence effectors (Rocafort *et al*., 2020). Part of the plant pathogenic fungi display dual-compartmentalized genome structures divided into gene-dense / repeat poor regions containing essential and highly conserved housekeeping genes, and gene-sparse / repeat-rich regions harboring fast-evolving genes, including effector-encoding genes (Dong *et al*., 2015; Sánchez-Vallet *et al*., 2018). These repeat-rich regions correspond to sub-telomeric regions, accessory chromosomes, or are dispersed along the chromosomes and are usually impacted by various mechanisms of rapid evolution: recombination, partial or complete gene deletion, mutation accumulation, transposon insertion (Lo Presti *et al.,* 2015). In ascomycetes, repeat-rich regions can also be submitted to RIP (Repeat-Induced Point mutation) mechanism, a premeiotic process controlling repeated element invasion by introducing cytosine to thymine mutations generating stop codons (Cambareri *et al*., 1989). As RIP can extend beyond the duplicated sequences up to 4 kb (Irelan *et al*., 1994), it also plays an active role in rapid gene evolution.

Genetic control, using plants displaying qualitative and/or quantitative resistance, is an efficient strategy to control fungal diseases (Jones and Dangl, 2006). Quantitative resistance is a partial resistance taking place at adult stage and under polygenic control usually involving several Quantitative Trait Loci (QTLs) with partial effect on the phenotype. These resistances are frequently dependent on the environment and do not prevent the colonization of the plant but limit the development of the disease and reduce symptom severity (Delourme *et al*., 2006; Niks *et al*., 2015; St.Clair, 2010). Conversely, qualitative resistance is usually conferred by race-specific resistance (*R*) genes recognizing defined effector genes through a gene-for-gene interaction (Flor, 1971). This recognition generally induces a localized cell death, called hypersensitive response (HR), preventing further colonization by avirulent isolates. However, in case of extensive use of a specific *R* gene, the resistance can be rapidly overcome by the pathogen that becomes virulent (Rouxel *et al*., 2003; Sprague *et al*., 2006). Several mechanisms allow pathogens to overcome *R* genes including a partial or complete deletion of the avirulence gene, a point mutation allowing to escape recognition while maintaining effector function, a down-regulation of the avirulence gene, or the acquisition of new effectors that suppress recognition. Many of these mechanisms are favored by the location of avirulence genes in repeat-rich regions of the fungal genomes, leading to a rapid bypass of specific resistant sources deployed in the fields. We hypothesize that *R* genes interacting with effector genes located in conserved regions of fungal genomes would be arduous to overcome.

*Leptosphaeria maculans* is an ascomycete responsible for one of the most damaging diseases on *Brassica napus* (rapeseed): phoma stem canker. This disease can cause yield losses reaching up to 50% and US$900 million per year (Fitt *et al*., 2008). Phoma stem canker epidemics are monocyclic and usually initiated by ascospores, produced after sexual reproduction on stem residues, landing on the aerial organs of *B. napus*. After spore germination, hyphae penetrate leaves and cotyledons through stomata or wounding. Once inside the plant, the fungus colonizes the apoplast during a short biotrophic stage (5 to 12 days) and then switches to a necrotrophic lifestyle, inducing leaf spots (Fitt *et al*., 2006; Rouxel and Balesdent, 2005). After leaf infection, a lengthy endophytic systemic colonization of leaf and stem tissues takes place and can last several months in Europe (West *et al*., 2001). At the end of cultural season, the fungus suddenly switches to necrotrophy causing crown canker responsible for lodging of the plant and associated yield losses. *L. maculans* exhibits a compartmentalized genome structure, with alternating GC-, gene-rich, and AT-, repeat-rich regions (Rouxel *et al*. 2011). Furthermore, Soyer *et al*. (2021) revealed that this dual compartmentalization was also visible through chromatin methylation enrichments during axenic growth, gene-rich regions being enriched in the trimethylation of the lysine 27 of histone H3 (H3K27me3) associated with facultative heterochromatin and in the di-methylation of the lysine 4 of histone H3 (H3K4me2) specific of euchromatin. In contrast, repeat-rich regions were enriched in the trimethylation of the lysine 9 of the histone H3 (H3K9me3) associated with constitutive heterochromatin.

During the complex interaction between *L. maculans* and rapeseed, an arsenal of effectors is secreted, facilitating infection of the host. In a first transcriptomic analysis comparing cotyledon and stem infection of *B. napus* by *L. maculans*, Gervais *et al*. (2017) differentiated two types of effector genes according to their expression kinetics: ‘early’ effector genes overexpressed during the cotyledon stage of infection, and ‘late’ effector genes or *LmSTEE* (*Leptosphaeria maculans* Stem Expressed Effectors, Jiquel *et al*., 2021) overexpressed during stem colonization. Complementing these results, a larger scale transcriptomics analysis covering all stages of interaction between *L. maculans* and its host identified clusters of effector gene expression (Gay *et al*., 2021). The first cluster grouped genes expressed during conidia germination and hyphae penetration into cotyledons and the third cluster genes expressed during necrotrophic stages of infection on cotyledons and petioles. Cluster 5 was specific to asymptomatic colonization of the stem and cluster 6 to the necrotrophic stage of stem infection. In contrast, clusters 2 and 4 contained genes overexpressed both at the early and late stages of infection, with genes of cluster 2 expressed during all the biotrophic/asymptomatic stages in cotyledons, petioles and stem and cluster 4 containing genes highly expressed during the shifts from biotrophy to necrotrophy in cotyledons and stem basis. Thus, the combination of both studies refined the classification of early effector genes to genes grouped in clusters 1 to 3 and late effector genes in clusters 4 to 6.

Most of the rapeseed resistant cultivars currently deployed in the field harbor specific *R* genes targeting early effector genes (Vasquez-Teuber *et al*., 2024) located in AT-rich regions. *Rlm1*, a resistance gene targeting the early effector *AvrLm1*, was overcome in less than three cultural cycles (Rouxel *et al*., 2003). The main event explaining the switch to virulence was a deletion of the genomic region containing *AvrLm1* (Gout *et al*., 2007). To identify more durable resistance sources to *L. maculans*, Jiquel *et al*. (2021) focused on late effector genes, located in gene-rich regions. They hypothesized that gene-for-gene interactions with late effector genes may occur during stem colonization and contribute to quantitative resistance in *B. napus.* To demonstrate this hypothesis, an innovative phenotyping method, consisting in overexpressing a selection of late effector genes at early stages of infection allowed the identification of new resistance sources. Indeed, four semi-winter cultivars, ‘Yudal’, RG021, RG047, and RG072 presented an HR when inoculated with transformants overexpressing *LmSTEE98*, a late effector gene from cluster 4, and one semi-winter cultivar, RG007, was similarly resistant to *LmSTEE6826,* a late effector gene from cluster 5 (Jiquel *et al*., 2021, 2022). CRISPR-Cas9 experiments inactivating *LmSTEE98* demonstrated that the interaction at the cotyledon stage followed a classical gene-for-gene scheme. In addition, stem inoculation with a wildtype isolate and transformants inactivated for *LmSTEE98* demonstrated that the presence of *RlmSTEE98* induced a reduction of symptom severity in the stem indicating that the *LmSTEE98* / *RlmSTEE98* interaction contributed to quantitative resistance phenotype (Jiquel *et al*., 2021).

These pioneer studies highlighted the interest of using late effector genes to search for new resistance sources. However, the refined classification brought by Gay *et al*. (2021) raised the question of whether all late effector genes shared the same characteristics and if they would be more stable than early effector genes, rendering the resistances they triggered more durable. In addition, Jiquel *et al*. (2022) highlighted new criteria for selecting late effectors and suggested that semi-winter rapeseed and rutabagas would be an interesting genetic pool to identify new resistance sources.

In this study, we tested these hypotheses by comparing the characteristics of a large set of early and late effector genes (genomic location, proximity of transposable elements (TE), enrichment in chromatin methylation marks, conservation in a worldwide collection of *L. maculans* isolates). We then selected six new late effector candidates using refined criteria of selection, and enriched the previous *B. napus* panel from Jiquel *et al*. (2022) with 100 additional semi-winter rapeseed and rutabaga genotypes. This study revealed that early and late effector genes diverged for most of their genomic characteristics strengthening the hypothesis of late effector genes being more conserved. Finally, we identified two new resistance sources to two late effector genes and 14 new genotypes resistant to *LmSTEE98*, all belonging to the semi-winter panel.

## RESULTS

### Early and late effector genes harbored highly distinct genomic surroundings

Among the 1207 genes up-regulated *in planta* described by Gay *et al*. (2021), we selected 151 genes predicted to encode effectors to possess a signal peptide or to have an extracellular location and one or no predicted transmembrane domain (Table S1). This selection was divided into 75 early effector genes (8 from cluster 1, 59 from cluster 2, and 8 from cluster 3) and 76 late effector genes (41 from cluster 4, 26 from cluster 5, and 9 from cluster 6).

The genomic location of effector genes was assessed using isochore annotation on the JN3 genome (Dutreux *et al*., 2018). We categorized three genomic locations: GC-equilibrated, AT-rich isochores, and borders corresponding to a 10kb overlapping region in between. We observed that late effector genes were distributed between GC-equilibrated isochores and borders. In contrast, early effector genes were equally present in the three types of genomic regions (Chi² test, p-value = 8.294e-10; Figure 1a).

**Figure 1:**
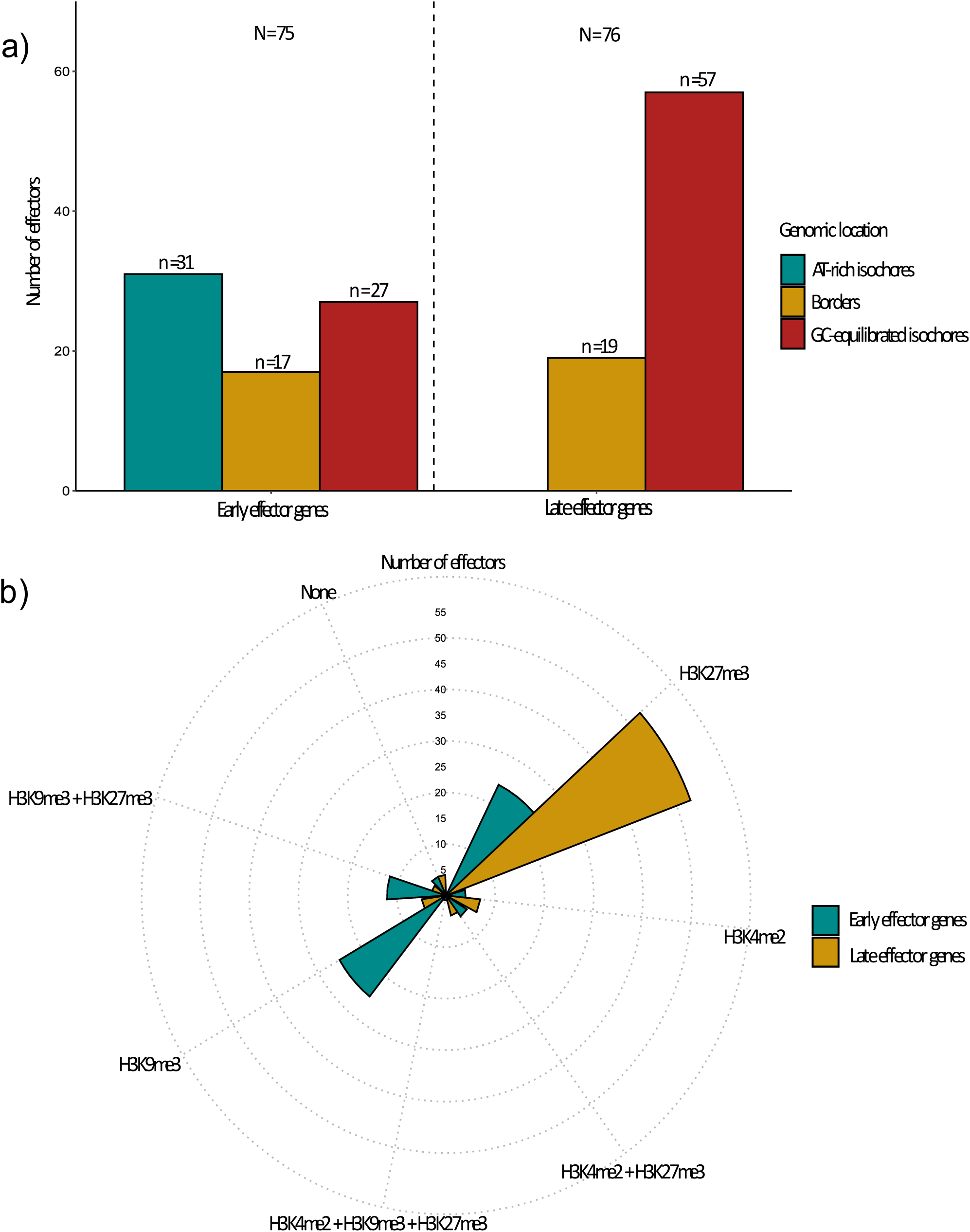
Genomic location (a) and chromatin methylation enrichment during axenic growth (b) of early and late effector genes from *Leptosphaeria maculans.* Seventy-five early and seventy-six late effector genes respectively overexpressed during rapeseed leaf infection and stem infection were characterized. Genomic location of effector genes was assessed using the annotations on JN3 genome (Dutreux *et al.,* 2018). Enrichment in chromatin methylation marks was assessed using ChiP-seq data from Soyer *et al*. (2021).

Soyer *et al*. (2021) previously characterized the chromatin landscape of *L. maculans* genome during axenic growth. Using these data, we found that early and late effector genes were essentially enriched in heterochromatin methylation marks. Indeed, 53 late effector genes were enriched in H3K27me3, 5 in H3K29me3 and 7 in H3K4me2. Seven late effector genes were enriched with both H3K27me3 and H3K4me2 or H3K27me3 and H3K9me3, and four presented no enrichment. Early effector genes also showed enrichments in heterochromatin methylation marks but presented a more even distribution between H3K27me3 and H3K9me3 (Chi² test; p-value = 1.964e-05; Figure 1b).

Using the TE annotation of Grandaubert *et al*. (2014), we assessed their presence, number, distance, and type up to 4kb around effector genes (Figure 2). There was no difference between early and late effector genes for the presence of TE in the interval (Chi² test, p-value = 0.09261; Figure 2a). However, both sets differed in the number of TE in their surroundings (Kruskal-Wallis test, p-value = 7.689e-06; Figure 2b) and the distance from the first TE (Kruskal-Wallis test, p-value = 1.028e-07; Figure 2c). There was a mean of 4.9 TE close to early effector genes, with the first TE being, on average, at 209 bp, compared to 1.5 TE close to late effector genes, and the first TE being at 1123 bp. Finally, we found no significant difference in the type of TE surrounding the two sets of effector genes, even if we noticed a higher frequency of the DNA transposon “DTx” close to late effector genes (31% compared to 5%; Figure 2d).

**Figure 2:**
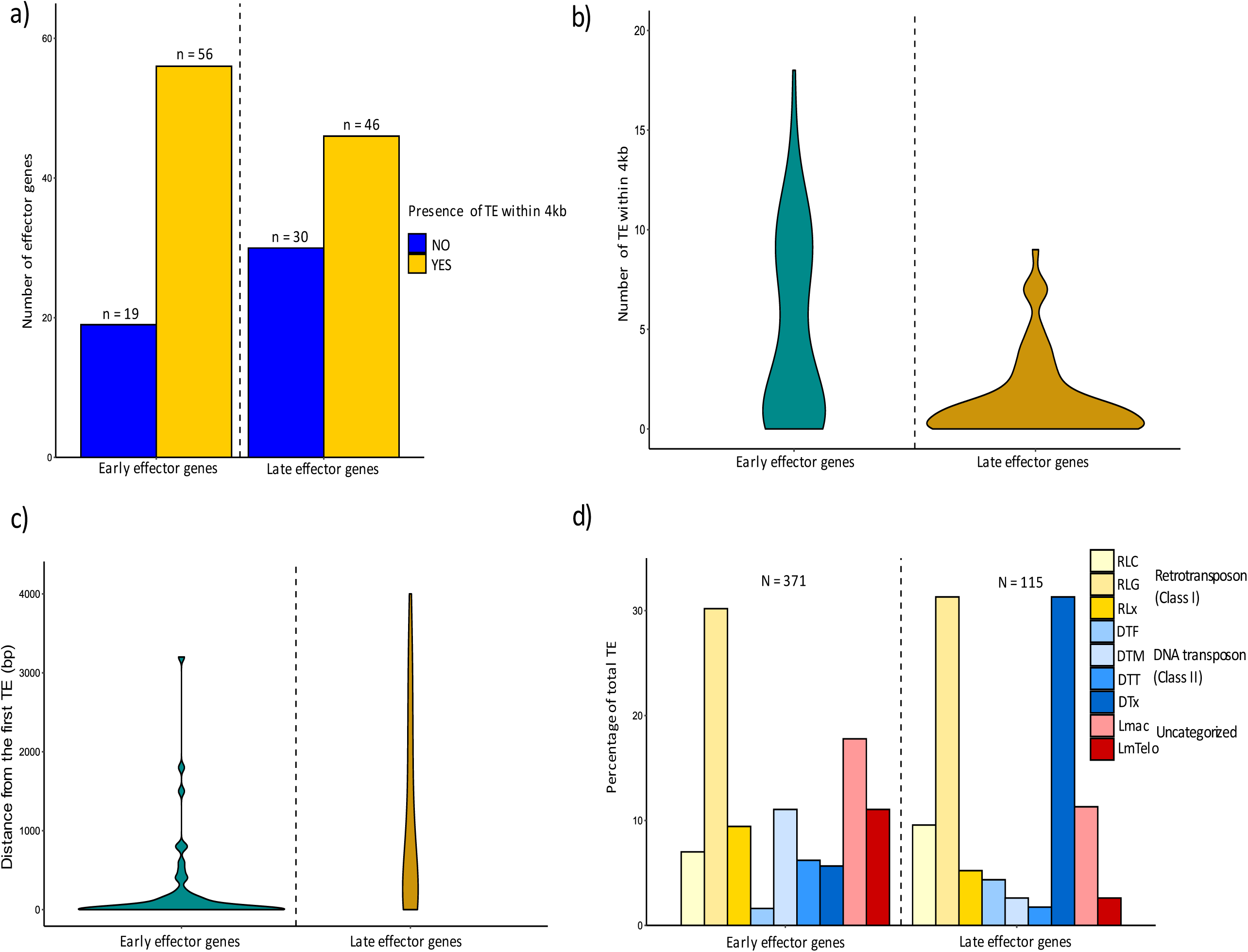
Presence (a), number (b), distance (c) and type (d) of transposable element (TE) in the genomic surroundings of effector genes of *Leptosphaeria maculans*. The presence, number, and distance of TEs within 4kb upstream and downstream of the effector genes was assessed using the annotations of Grandaubert *et al*. (2014). The type of TE (DNA transposon, retrotransposon or uncategorized) was calculated as a percent of the total number of TE in the surrounding genomic region.

### Late effector genes were more conserved than early effector genes in a worldwide collection of *L. maculans* isolates

We used the genomic data of a worldwide collection gathering 205 isolates of *L. maculans* (Van de Wouw *et al*., 2024) to determine whether the 151 effector genes were conserved. We were able to assemble 201 isolate genomes and determined protein sequences for the 151 effectors using JN3 as a reference (Table S2). Four effector genes were not sufficiently well assembled to be analyzed: two from cluster 2 (*Lmb_jn3_08794* and *Lmb_jn3_12242*), one from cluster 5 (*Lmb_jn3_07669* ) and one from cluster 6 (*Lmb_jn3_09959*).

Nine early and four late effectors were absent in more than 10% of the isolates. Three effector genes were absent from a large majority of isolates: *Lmb_jn3_03238* and *AvrLm1 (Lmb_jn3_13126*) (cluster 2) respectively present in 70 and 34 isolates and *Lmb_jn3_05550* (cluster 5), present in 11 isolates. We found that late effectors were significantly more present in the collection than early effectors, with a mean difference of 4 isolates (Kruskal-Wallis test, p-value = 0.0003379). The number of protein isoforms varied from 0 to 26 with a mean of 2.9 for early effectors and 1.2 for late effectors (Kruskal-Wallis, p-value = 0.004418; Figure 3a). The percentage of isolates carrying a protein isoform differing from JN3 ranged from 0 to 97% with a mean of 4% for early effectors and 0.32% for late effectors (Kruskal-Wallis, p-value = 0.0014; Figure 3b).

**Figure 3:**
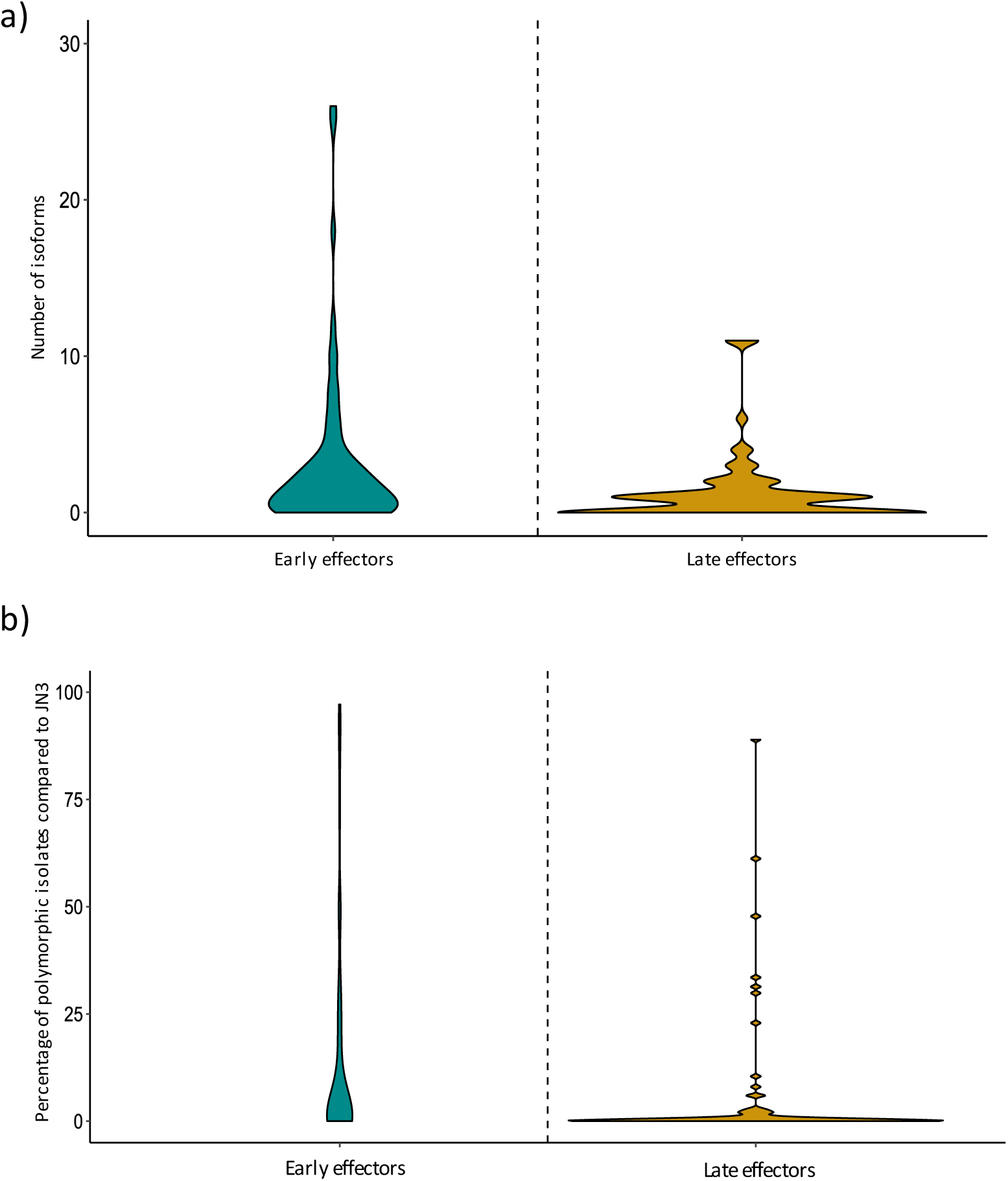
Conservation of early and late effectors in a worldwide collection of *Leptosphaeria maculans* isolates (Van de Wouw *et al.,* 2024). We compared the protein sequences of 201 *L. maculans* isolates for 73 early and 74 late effectors, respectively overexpressed during rapeseed leaf infection and stem infection, using JN3 as a reference. We assessed the number of isoforms identified for each effector (a) and the percentage of isolates carrying a polymorphic effector compared to JN3 proteome (b).

New selection criteria led to a set of six promising effector candidates to uncover new resistance sources Jiquel *et al*. (2022) had displayed recommendations for the selection of late effectors candidate genes toward identification of new plant resistance: (i) belonging to cluster 4 or 5, (ii) having no homolog in other fungal species and (iii) being located in a GC-equilibrated region or in a border.

Among the 67 late effectors genes belonging to clusters 4 and 5, we first selected 54 genes not previously studied (Gervais *et al*., 2017; Jiquel *et al*., 2021, 2022). Using Gay *et al*. (2021) gene expression analysis, we conserved effectors having an expression profile similar to *LmSTEEE98* or *LmSTEE6826* in controlled and field conditions (i.e. genes overexpressed during stem infection, preferentially during early stages of stem colonization, but with a low expression in cotyledons; Figure S1) leading to 32 candidates. Among these candidates, only 16 had no homolog in other fungal species. We selected candidates enriched in H3K27me3 chromatin methylation marks leading to 11 potential candidates. Finally, when a reliable 3- D structure prediction was available using AlphaFold (Mirdita *et al*., 2022), we favored a diversity of structural families in our six final candidates (Table 1).

**Table 1:**
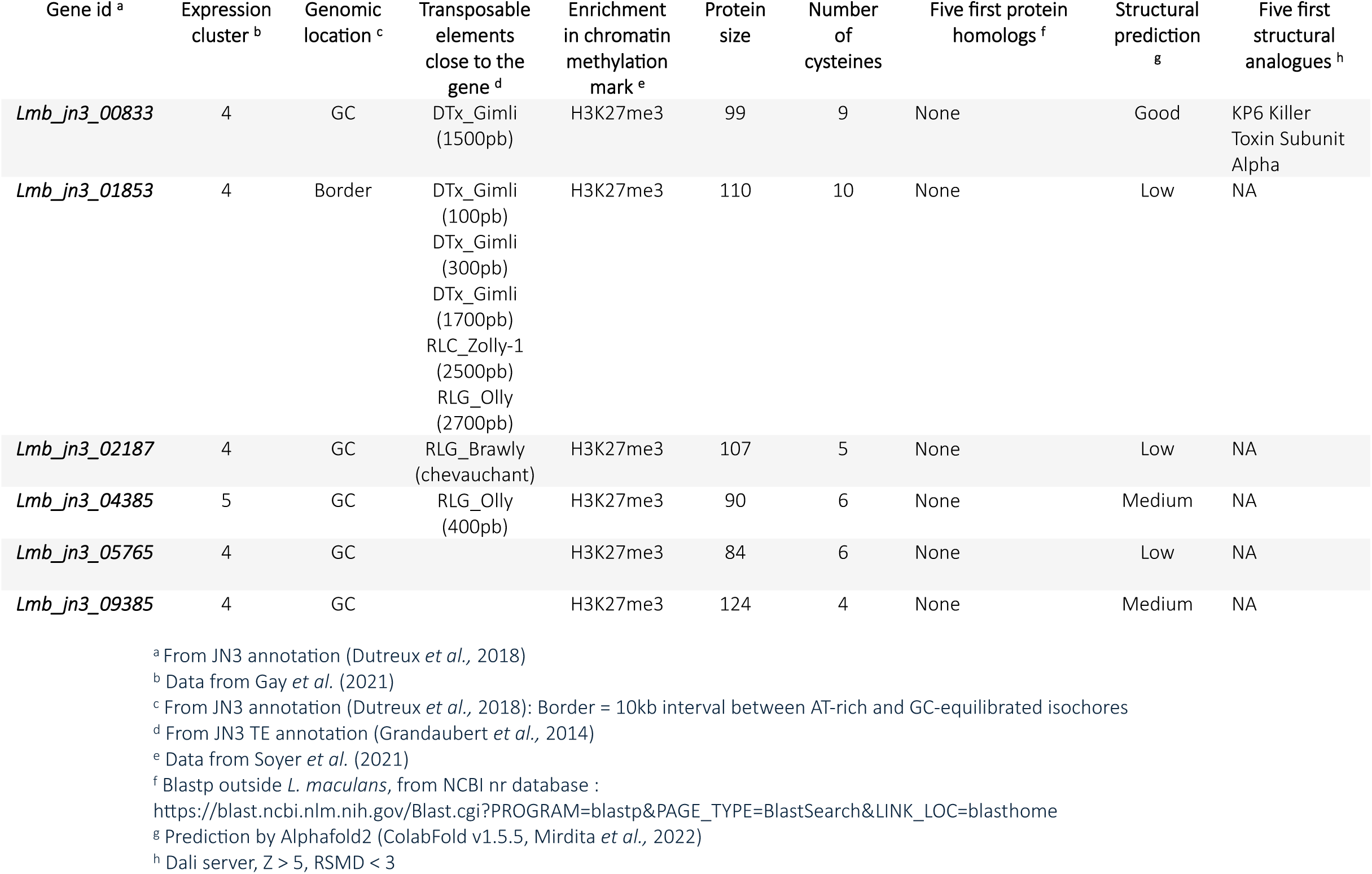
Description of the six *Leptosphaeria maculans* late effector genes selected to uncover new resistance sources in *Brassica napus*.

Five of these candidates were assigned to cluster 4 and one to cluster 5. They all encoded small secreted proteins ranging from 84 to 124 amino acids (aa). Two were enriched in cysteines, *Lmb_jn3_00833* with 9 cysteines for 99 aa (9%) and *Lmb_jn3_01853* with 10 cysteines for 110 aa (9%). Five candidate genes were located in GC-equilibrated regions while *Lmb_jn3_01853* was located in a border. Two candidates, *Lmb_jn3_00833* and *Lmb_jn3_09385*, harbored a DNA transposon ‘Gimli’ respectively at 200 bp and 1500 bp. Two other candidates harbored a retrotransposon at less than 400 bp. One candidate was not surrounded by TE and one harbored three DNA transposons ‘Gimli’ and two retrotransposons. Except for *Lmb_jn3_00833*, which was predicted to have structural analogy with KP6 Killer Toxin proteins, all candidates had a medium to low structural prediction.

These six candidate effector genes were renamed *LmSTEE833*, *LmSTEE1853*, *LmSTEE2187*, *LmSTEE4385*, *LmSTEE5765,* and *LmSTEE9385* and completed the ten effector genes studied in Jiquel *et al* studies: *LmSTEE1*, *LmSTEE1277*, *LmSTEE1852*, *LmSTEE35*, *LmSTEE5465*, *LmSTEE78*, *LmSTEE7919*, *LmSTEE10933*, *LmSTEE6826*, and *LmSTEE98*.

### Overexpression of new *LmSTEE* genes at the cotyledon stage of infection

We placed the six effector genes selected for this study, plus *LmSTEE98* (Jiquel *et al*., 2021) and *LmSTEE6826* (Jiquel *et al*., 2022) effector genes, under the control of the *AvrLm4-7* promoter. We introduced the eight constructs into the X83.51 isolate, virulent toward most known *Rlm* genes. We obtained from four, up to twelve transformants for each construct. Transformants growth was tested on V8 agar medium and pathogenicity by inoculation on a susceptible genotype, Darmor. All transformants displayed no growth or pathogenicity defects except for pA4-7::*LmSTEE98*.15 and pA4-7::*LmSTEE98*.16 which were not virulent on Darmor. All transformants were amplified and sequenced to confirm the correct insertion of the construct. We tested the level of *LmSTEE* gene expression at 7dpi in a maximum of four transformants per gene. Expression in the infected cotyledons varied between 1.10E^02^ and 1.10E^04^ compared to actin expression, and all transformants overexpressed the *LmSTEE* gene when compared with the native gene in X83.51 (from a factor 100 to a factor 1000; Figure 4). For each *LmSTEE*, the two transformants having the highest level of expression at 7 dpi and a correct sequencing result were selected for the screening of 207 genotypes of *B. napus* (Table S3).

**Figure 4:**
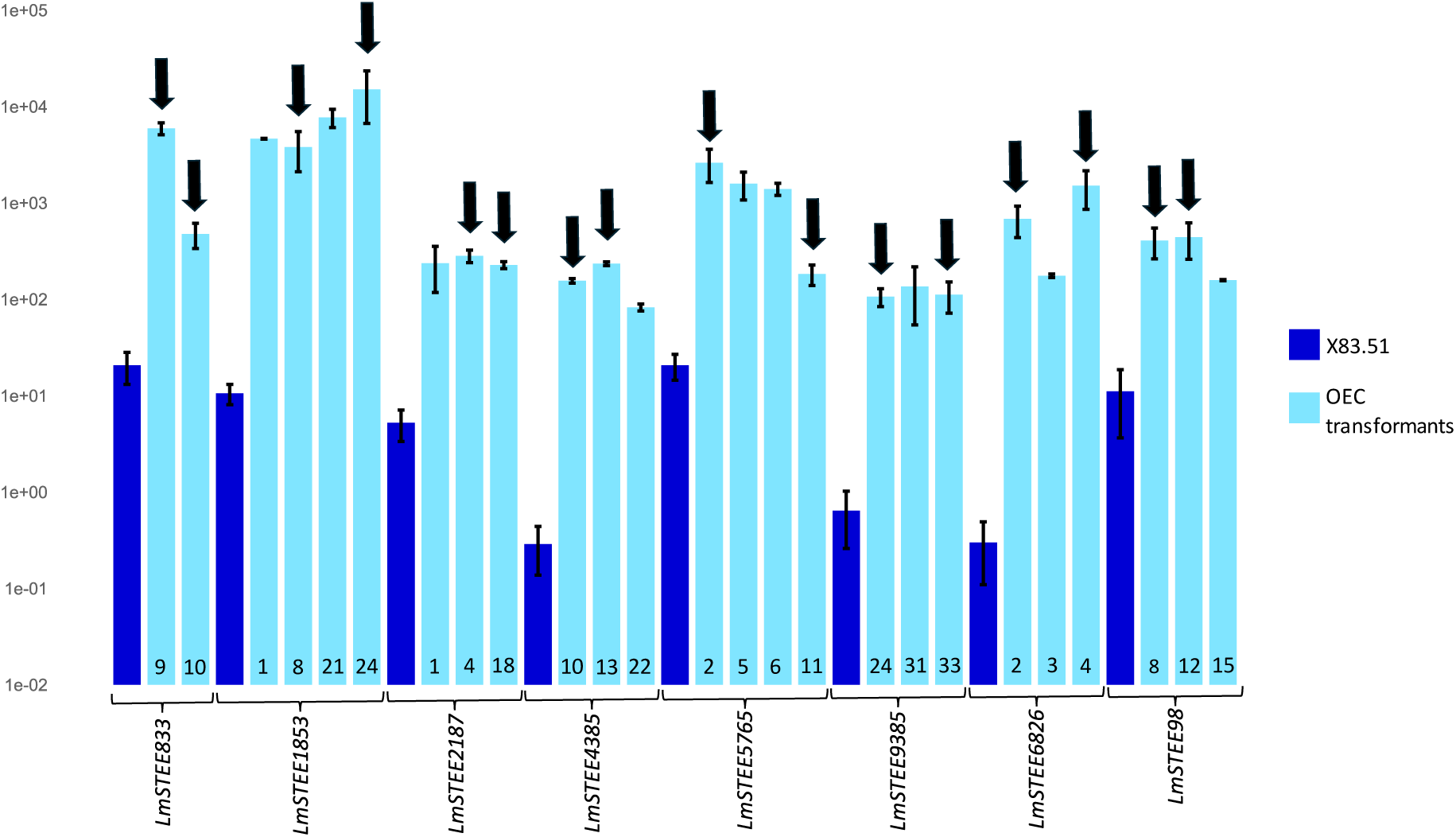
Expression of six new LmSTEE genes in ‘overexpressed in cotyledons’ (OEC) transformants of *Leptosphaeria maculans* at an early stage of cotyledon colonization. We obtained OEC transformants overexpressing the *LmSTEE* genes during rapeseed cotyledon infection by putting these genes under the control of *AvrLm4-7* promoter. The expression of *LmSTEE* genes was assessed by quantitative reverse transcription (qRT)-PCR in infected cotyledons of the cultivar Darmor, 7 days post-inoculation. Transformant identification numbers are mentioned at the bottom of expression bars. Black arrows indicate OEC transformants selected for the screening of plant genotypes. Mean expression is normalized against *actin*, with *EF1α* used as a control (Fudal *et al*., 2007). Error bars represent the standard error for two biological and two technical replicates.

### New resistance responses were highlighted using new *LmSTEE* genes and an enlarged *B. napus* panel

OverExpressed in Cotyledons (OEC) transformants expressing *LmSTEE* genes described by Jiquel *et al*. (2021, 2022) were inoculated on 100 genotypes belonging to the semi-winter genetic pool, including rapeseed and rutabaga (Table 2, Table S4). This screening revealed 14 genotypes, 13 rapeseed and one rutabaga, displaying an HR when inoculated with *LmSTEE98* OEC transformants in INV13.269 genetic background. All these 14 genotypes displayed the same resistance response when inoculated with *LmSTEE98* OEC transformants obtained during the present study in the X83.51 background (Figure 5a).

**Figure 5:**
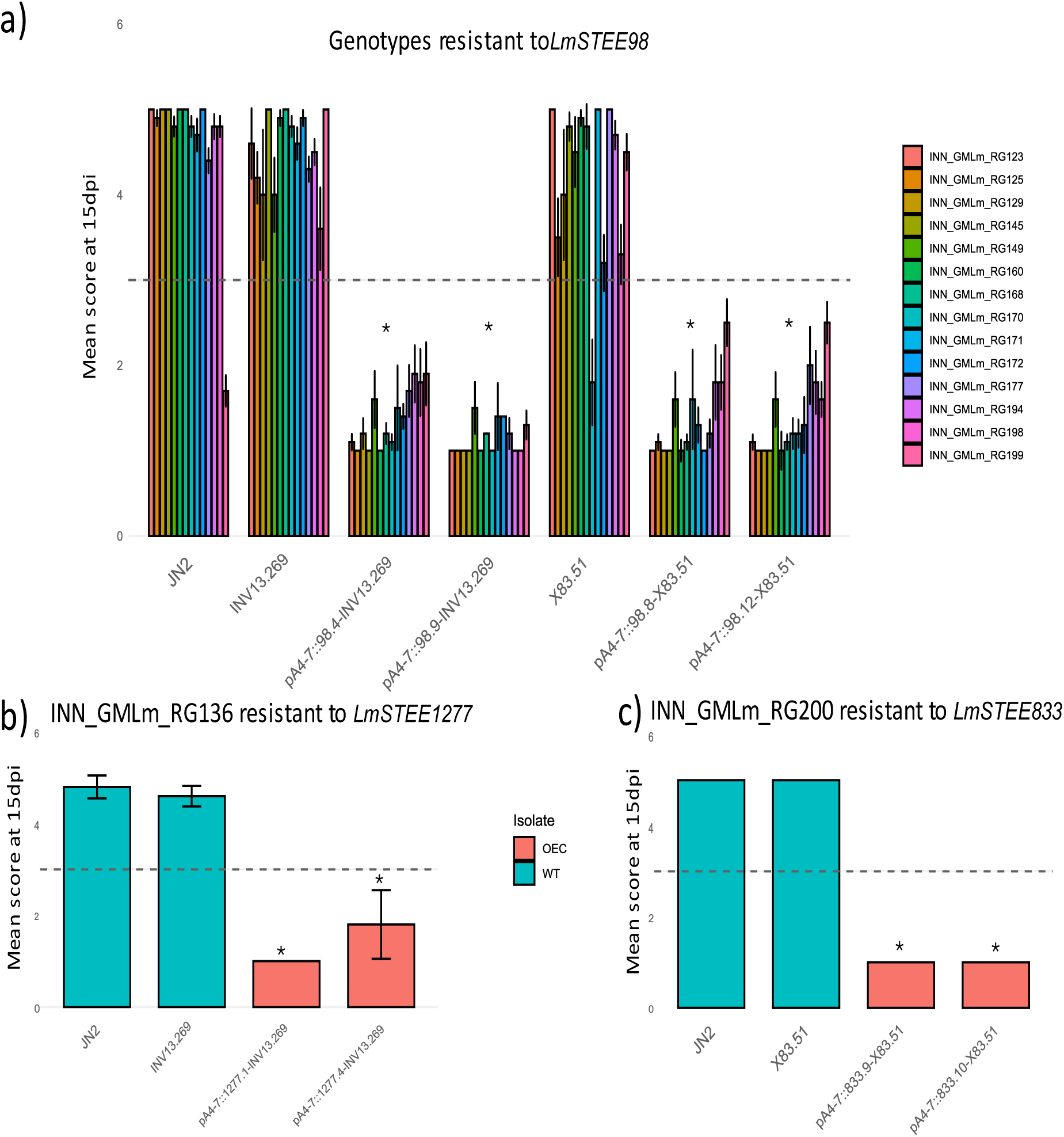
Resistance of *B. napus* to *LmSTEE* genes revealed by pathogenicity assays using transformants overexpressing these genes at the cotyledon stage of infection. (a) 14 semi-winter genotypes displayed hypersensitive responses (HR) when inoculated with *LmSTEE98* OverExpressed in Cotyledons (OEC) transformants (in INV13.269 and X83.51 backgrounds). (b) INN_GMLm_RG136 displayed an HR when inoculated with *LmSTEE1277* OEC transformants. (c) INN_GMLm_RG200 displayed an HR when inoculated with *LmSTEE833* OEC transformants. Scoring was performed 15 days post-inoculation. Scores from 1 to 3 correspond to a resistant phenotype and scores from 4 to 6 to a susceptible phenotype (IMASCORE scale, Balesdent *et al*. (2001)). Error bars represent the standard error for ≥ 6 biological replicates. The asterisks indicate a significant difference between the wild-type isolates and the OEC transformants (Kruskal–Wallis test: *p-value < 0.001)

**Table 2:** Incompatible interactions between *Brassica napus* genotypes and *LmSTEE* genes.

| Genotype | Diversity group | *LmSTEE*<sup>a</sup> | Type of resistance response<sup>b</sup> |
| --- | --- | --- | --- |
| INN_GMLm_RG123 | Semi-winter | *LmSTEE98* (INV13.269 and X83.51 background) | HR |
| INN_GMLm_RG125 | Semi-winter | *LmSTEE98* (INV13.269 and X83.51 background) | HR |
| INN_GMLm_RG129 | Semi-winter | *LmSTEE98* (INV13.269 and X83.51 background) | HR |
| INN_GMLm_RG145 | Semi-winter | *LmSTEE98* (INV13.269 and X83.51 background) | HR |
| INN_GMLm_RG149 | Semi-winter | *LmSTEE98* (INV13.269 and X83.51 background) | HR |
| INN_GMLm_RG160 | Semi-winter | *LmSTEE98* (INV13.269 and X83.51 background) | HR |
| INN_GMLm_RG168 | Semi-winter | *LmSTEE98* (INV13.269 and X83.51 background) | HR |
| INN_GMLm_RG170 | Semi-winter | *LmSTEE98* (INV13.269 and X83.51 background) | HR |
| INN_GMLm_RG171 | Semi-winter | *LmSTEE98* (INV13.269 and X83.51 background) | HR |
| INN_GMLm_RG172 | Semi-winter | *LmSTEE98* (INV13.269 and X83.51 background) | HR |
| INN_GMLm_RG177 | Semi-winter | *LmSTEE98* (INV13.269 and X83.51 background) | HR |
| INN_GMLm_RG194 | Semi-winter | *LmSTEE98* (INV13.269 and X83.51 background) | HR |
| INN_GMLm_RG198 | Semi-winter | *LmSTEE98* (INV13.269 and X83.51 background) | HR |
| INN_GMLm_RG199 | Semi-winter | *LmSTEE98* (INV13.269 and X83.51 background) | HR |
| INN_GMLm_RG108 | Semi-winter | *LmSTEE98* (INV13.269 background) | HR |
| INN_GMLm_RG110 | Semi-winter | *LmSTEE98* (X83.51 background) | HR |
| INN_GMLm_RG136 | Semi-winter | *LmSTEE1277* (INV13.269 background) | HR |
| INN_GMLm_RG078 | Spring | *LmSTEE5765* (X83.51 background) | HR |
| INN_GMLm_RG200 | Semi-winter | *LmSTEE833* (X83.51 background) | HR |
| INN_GMLm_RG047 | Semi-winter | *LmSTEE5765* (X83.51 background) | Intermediate resistance |
| INN_GMLm_RG024 | Semi-winter | *LmSTEE5765* (X83.51 background) | Heterogeneous resistance |
| INN_GMLm_RG056 | Semi-winter | *LmSTEE5765* (X83.51 background) | Heterogeneous resistance |
| INN_GMLm_RG089 | Semi-winter | *LmSTEE6826* (X83.51 background) | Heterogeneous resistance |
| INN_GMLm_RG136 | Semi-winter | *LmSTEE10933* (INV13.269 background) | Heterogeneous resistance |
| INN_GMLm_RG139 | Semi-winter | *LmSTEE35* (INV13.269 background) | Heterogeneous resistance |
| INN_GMLm_RG151 | Semi-winter | *LmSTEE35* (INV13.269 background) | Heterogeneous resistance |
| INN_GMLm_RG151 | Semi-winter | *LmSTEE5465* (INV13.269 background) | Heterogeneous resistance |
| INN_GMLm_RG151 | Semi-winter | *LmSTEE10933* (INV13.269 background) | Heterogeneous resistance |
| INN_GMLm_RG163 | Semi-winter | *LmSTEE1853* (X83.51 background) | Heterogeneous resistance |
| INN_GMLm_RG163 | Semi-winter | *LmSTEE9385* (X83.51 background) | Heterogeneous resistance |
| INN_GMLm_RG198 | Semi-winter | *LmSTEE5765* (X83.51 background) | Heterogeneous resistance |
| INN_GMLm_RG020 | Unknown | X83.51 | HR |
| INN_GMLm_RG072 | Semi-winter | X83.51 | HR |
| INN_GMLm_RG090 | Semi-winter | X83.51 | HR |
| INN_GMLm_RG091 | Semi-winter | X83.51 | Heterogeneous resistance |
| INN_GMLm_RG093 | Unknown | X83.51 | HR |
| INN_GMLm_RG147 | Semi-winter | X83.51 | Heterogeneous resistance |
| INN_GMLm_RG170 | Semi-winter | X83.51 | HR |
| INN_GMLm_RG172 | Semi-winter | X83.51 | HR |
| INN_GMLm_RG178 | Semi-winter | X83.51 | HR |
| INN_GMLm_RG179 | Semi-winter | X83.51 | HR |
| INN_GMLm_RG078 | Spring | X83.51 | Heterogeneous resistance |

207 genotypes were screened using transformants overexpressing six new *LmSTEE* genes and ten *LmSTEE* genes previously selected for the identification of new resistance sources (Jiquel et al., 2021, 2022).
<sup>a</sup> *LmSTEE* genes involved in the interaction with the genotype and background isolates in which the *LmSTEE* gene was introduced to obtained OverExpressed in Cotyledons (OEC) transformants (INV13.269 or X83.51)
<sup>b</sup> Resistance phenotype observed during the interaction. HR = hypersensitive response, intermediate resistance corresponds to a delayed resistance phenotype and heterogeneous resistance corresponds to an alternance of susceptible and resistant phenotype for a genotype x *LmSTEE* interaction.

In addition, an HR was induced by *LmSTEE1277* on the semi-winter rapeseed INN_GMLm_RG136. This genotype displayed a resistance response towards the two independent OEC transformants but not with the wildtype isolates INV13.269 (Kruskal-Wallis test, p-value = 0.001; Figure 5b).

The screening of the 207 genotypes with new *LmSTEE* genes also revealed promising interactions (Table 2, Table S4). The rutabaga INN_GMLm_RG200 was resistant to *LmSTEE833* (Kruskal-Wallis test, p-value = 1.542e-08; Figure 5c) and the spring genotype INN_GMLm_RG078 displayed an HR when inoculated with *LmSTEE5765* OEC transformants. However, depending on the biological replicate, this genotype switched between susceptible and resistance responses when inoculated with X83.51, making the resistance phenotype induced by *LmSTEE5765* difficult to interpret.

One intermediate resistance corresponding to a delayed resistance phenotype was triggered by *LmSTEE5765* on the semi-winter rapeseed INN_GMLm_RG047. Several heterogeneous resistance phenotypes were revealed by *LmSTEE* transformants on various semi-winter genotypes, resulting in an alternance of susceptible and resistant phenotypes. Finally, ten genotypes harbored resistance phenotypes when inoculated with X83.51.

Except for the spring genotype INN_GMLm_RG078, all genotypes displaying an HR to *LmSTEE* genes belonged to the semi-winter genetic pool, including rapeseed and rutabaga (Figure 6).

**Figure 6:**
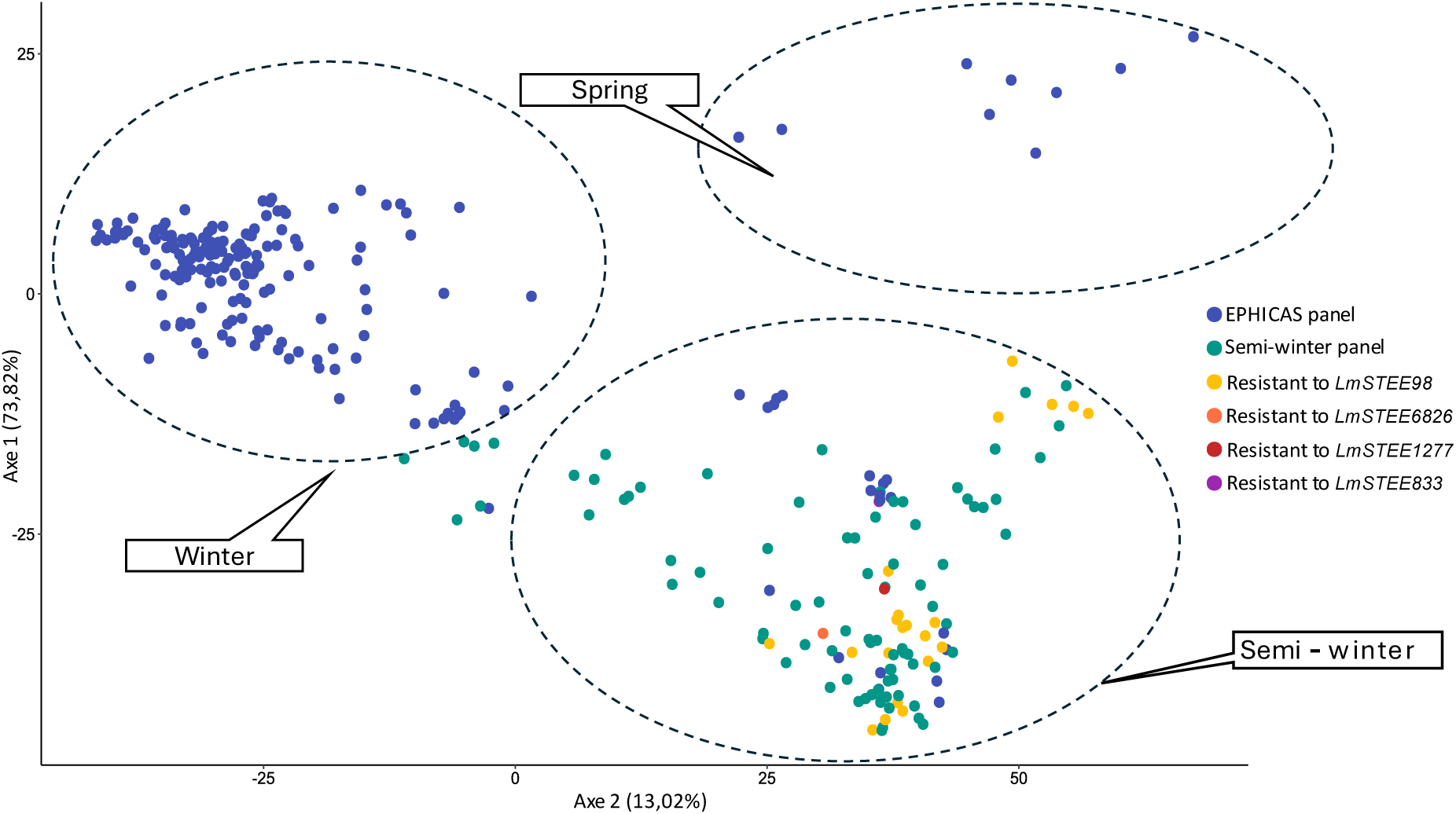
Genotypic diversity of *Brassica napus* genotypes used for the screening experiments. 207 genotypes were screened including 107 genotypes form the EPHICAS panel (Jiquel *et al.,* 2022) and 100 from the semi-winter panel (present study). A principal coordinate analysis was performed based on an Identity by State matrix resulting from a genome-wide genotyping with 6631 Single Nucleotide Polymorphisms. Black frames distinguish the three main types of *B. napus*: winter, spring, and semi-winter. Genotypes resistant to *LmSTEE* genes are indicated in yellow, orange, red and purple.

## DISCUSSION

Genetic resistance is the most efficient strategy to control stem canker disease in rapeseed crops. However, as gene-for-gene resistance to *L. maculans* conferred by *Rlm* genes can rapidly be overcome in the fields, there is an urge to find new resistance sources with better durability combined with a coordinated and managed R gene deployment in the fields (Vasquez-Teuber *et al*., 2024). In the present study, we revealed that late effector genes highly differed in their genomic and epigenomic context from early effector genes. We also showed that late effector genes were more conserved in a worldwide collection of *L. maculans* isolates, suggesting that the potential cognate resistances they could trigger would be more difficult to overcome. In addition, the screening assays confirmed the interest of late effector genes in uncovering new resistance sources, with the identification of three new potential gene-for-gene interactions. Except for one spring genotype, all resistance sources were found in the semi-winter genetic pool, emphasizing the importance of this latter to identify resistances to late effector genes.

We studied a set of 151 effector genes (75 early and 76 late) and revealed that their genomic characteristics were significantly distinct. Late effector genes were specifically located in GC-equilibrated isochores and borders with AT-rich isochores while early effector genes were well distributed between the three genomic locations, with all known *AvrLm* genes located in AT-rich isochores in agreement with previous studies. Rouxel *et al*. (2011) described AT-rich isochores as a genomic environment promoting rapid sequence diversification and underpinning evolutionary potential of the fungus to rapidly adapt to novel host-derived constraints. The absence of late effector genes from this dynamic genomic environment suggests that they are not affected by rapid evolutionary mechanisms and are conserved in *L. maculans* populations. TEs also have an important role in this rapid genomic evolution. Indeed, they can be inserted in gene sequences or promoters and are targeted by RIP mechanism. RIP was detected in *L. maculans* by Idnurm and Howlett (2003) and is known to have an active role in effector gene rapid evolution as it can extend to 4kbp beyond the targeted duplicated sequence (Irelan *et al*., 1994). To determine if late effector genes could be affected by the presence of TEs, we investigated their presence, number, distance, and type 4kb around the 151 effector genes. We demonstrated that both sets of effector genes are not different in terms of presence of TEs in their close genomic surrounding. However, early effector genes harbored, on average, three times more TEs than late effector genes, and their first TE was six times closer. The lower number of TEs in late effector genes environment also suggests that they could be less affected by RIP leaks, as RIP in *L. maculans* required large repeated regions (Van De Wouw *et al*., 2019). In addition, the higher distance between late effector genes and their first TE suggests that TEs are less susceptible to affect their promoter or coding sequences. These results support the hypothesis that late effector genes are better conserved in *L. maculans* populations, and that their associated resistances, if existing, may be more durable.

To test this hypothesis, we determined the presence and polymorphism of early and late effector genes in a worldwide collection. We demonstrated that late effector genes : (i) were significantly more present in the collection, (ii) presented half less isoforms, and (iii) had 12 times fewer polymorphic isolates compared to JN3 than early effector genes. Except for *AvrLm10A*, all known *AvrLm* gene were absent or polymorphic in at least 20% of isolates, with a maximum observed for *AvrLm1* (absent in 167 isolates and polymorphic in three isolates). In contrast, *LmSTEE* genes, described in previous and present studies, presented some polymorphic isolates ranging from one for *AvrLmSTEE6826* to four for *AvrLmSTEE98*, all isoforms differing for one amino acid substitution. These results confirmed the better conservation of late effector genes in *L. maculans* populations. However, these analyses are preliminary as Illumina Novaseq sequencing was used by Van de Wouw *et al*. (2024), making the assembly and annotation difficult. In the analysis for isoform number and polymorphic isolates ratio, we did not consider isolates lacking the start or the end of the protein, as it could be due to an error during sequencing or assembly. Long read sequencing should be used to validate these first results.

The results of both genomic characteristics and conservation analyses in *L. maculans* populations highlighted the interest in late effector genes to uncover new resistance sources in *B. napus*. Using Jiquel *et al*. (2022) recommendations, we selected six new late effector genes added to *AvrLmSTEE98* and *AvrLmSTEE6826,* previously characterized, and expressed them in X83.51 under the control of *AvrLm4-7* promoter. We confirmed the resistance phenotypes triggered by *AvrLmSTEE98* in the four genotypes Yudal, RG021, RG047 and RG072 and by *AvrLmSTEE6826* in RG007 (Jiquel *et al*., 2021, 2022).

We also revealed new resistance sources to *LmSTEE* genes through the combination of two strategies: the use of a new set of *LmSTEE* genes added to the ten previously selected, and the enrichment of *B. napus* panel in semi-winter rapeseed and rutabaga genotypes. Indeed, the screening of a semi-winter panel revealed 14 new genotypes resistant to LmSTEE98. While the four already known genotypes resistant to *LmSTEE98* were all from Asian origins (Korea and Japan), among the 14 genotypes uncovered in this study six were from Japan and two from Korea, but four genotypes were from Australia, one from Germany and one from the USA (Table S5). However, as Australian breeding programs were initiated with Japanese spring varieties and French winter varieties (Salisbury *et al*., 1995), we can hypothesize that most resistance sources to *LmSTEE98* are derived from Asian genotypes. The absence of Asian isolates in the IBCN collection (Van de Wouw *et al*., 2024) did not allow us to determine whether *AvrLmSTEE98* was conserved in Asian isolates or if the gene was subject to selection pressure in these populations.

Screening semi-winter and rutabaga genotypes also allowed the discovery of two new resistances induced by *LmSTEE* genes. The Japanese semi-winter rapeseed INN_GMLm_RG136 displayed HR to *LmSTEE1277,* and the Danish rutabaga INN_GMLm_RG200 was resistant to *LmSTEE833*. *LmSTEE1277* had already been studied by Jiquel *et al*. (2022), but no HR had been observed, confirming the interest in the semi-winter genetic pool. The absence of resistance sources to *LmSTEE* genes in the winter genetic pool compared to the semi-winter rise questions about their divergence. We can hypothesize a sampling bias, an independent evolution between both pools or a linkage disequilibrium between resistance to *LmSTEE* genes and another deleterious character which would have been eliminated by selection from the elite varieties composing winter genotypes.

One last resistance phenotype was more difficult to interpret. Indeed, INN_GMLm_RG078 presented a resistance phenotype to *LmSTEE5765* for all experiments; however, depending on the experiment, the genotype alternated between susceptibility and resistance to X83.51, making it difficult to conclude on the sole recognition *of LmSTEE5765* to trigger the HR. INN_GMLm_RG078 is an old (1963) German elite line with 13% of residual heterogeneity (Table S5) potentially acting on the resistance phenotype to X83.51. We can hypothesize that INN_GMLm_RG078 has a resistance gene recognizing X83.51 (eventually *Rlm5*, *Rlm10* or *Rlm14*) but not fixed at the locus, explaining the heterogeneity of the phenotype. Additionally, even if performed in controlled conditions, the pathogenicity assays may have been subjected to temperature or humidity variations that could have influenced the resistance phenotype, as previously found for other *AvrLm-Rlm* interactions (Neik *et al*., 2022; Yang *et al*., 2021). One intermediate resistance interaction was induced by *LmSTEE5765* in INN_GMLm_RG047, a Korean semi-winter rapeseed already known for its resistance to *LmSTEE98*. First described in Blondeau *et al*. (2015), intermediate resistance phenotypes correspond to delayed recognition of an AvrLm protein by its cognate Rlm protein. As already postulated by Jiquel *et al*.(2022), this type of interaction could derive from a time lag between the peak of expression of *LmSTEE5765* in the OEC transformants and the expression profile of the corresponding *Rlm* gene. Another hypothesis is a partial recognition of the AvrLm protein due to an allelic variation in its encoding gene (Blondeau *et al*., 2015)

The two strategies developed in this study for uncovering new resistant sources were efficient. In one hand, the enlargement of the screening panel with genotypes from the semi-winter genetic pool allowed the identification of 14 new resistance sources to *LmSTEE98* and one to *LmSTEE1277*, two late effector genes already studied by Jiquel *et al*. (2021, 2022). In the other hand, the selection of six new late effector genes with improved selection criteria permitted the uncovering of two other resistance sources, one to *LmSTEE833* and one to *LmSTEE5765*. Resistance to *LmSTEE833* was found in a rutabaga, underlying the importance of combining the two strategies to maximize the chances of discovering new resistances.

Our study provides new perspectives for the identification of new resistant genotypes and their use for durable disease management strategies. However, resistance to *LmSTEE* genes has not yet been tested in the field. As semi-winter genotypes are not adapted to European growing conditions, resistance to *LmSTEE* genes should be tested by introgression in elite winter cultivars. In addition, while we proved that late effector genes were more conserved in *L. maculans* worldwide population, there is currently no proof that this conservation would be maintained in case of extensive use of the corresponding resistance. Before deployment of resistances to late effectors, *L. maculans* population should be monitored for *LmSTEE* genes polymorphism in the field in order to avoid rapid evolution of the pathogen against these new resistances.

## EXPERIMENTAL PROCEDURES

### Fungal and plant materials

The JN2 isolate (v.23.1.2; Balesdent *et al*., 2002), closely related to JN3 (v.23.1.3; Balesdent *et al*., 2002), INV13.269 (Plissonneau *et al*., 2016) and X83.51 were used as controls in inoculation tests. X83.51 results from a cross between IBCN14 (Balesdent *et al*., 2005) and X22.14 itself being an offspring of INV13.269 crossed with WT50-3 (Neik *et al*., 2020). OEC transformants described in Jiquel *et al*., (2021, 2022) were used for inoculation tests. All fungal isolates used in this study are described in Table S3. Fungal isolates were grown and sporulated on V8 juice agar medium as described by Ansan-Melayah *et al*., (1995).

The *B. napus* panel is composed of 207 genotypes. The first 107 genotypes were described by Jiquel *et al*. (2022) and are referred to as the “EPHICAS panel”. This panel was set by genotyping *B. napus* genotypes with an internal array of 6331 SNPs covering the 19 chromosomes. A kinship matrix was created with the R package emma using the genotyping results. This Identity by State (IBS) matrix was used to compute a principal coordinate analysis (PCA) using the R packages FactoMineR and factoextra. Graphical representations were obtained using the R package ggplot2.

Using the same method, 100 genotypes were added to the “EPHICAS panel” for their belonging to the semi-winter diversity group which included rapeseed and rutabaga. This second panel is referred to as the “semi-winter panel.” Genotypes from EPHICAS and semi-winter panels are described in Table S5.

### Plant inoculation tests

Plant inoculation tests were performed independently in two locations with different protocols. The *LmSTEE* transformants described by Jiquel *et al*., (2021, 2022) were inoculated on the semi-winter panel at Innolea. Fifteen individuals per genotype were distributed in three repetitions of five plants following a longitudinal gradient in a growth chamber. The plants were inoculated with two control isolates, JN2 and INV13.269 and with one transformant per *LmSTEE* gene. One isolate was deposited on each half of the cotyledons. Inoculation was performed by puncturing 12-day-old seedlings and deposing 10 µL of inoculum (10^7^ pycnidiospores per mL) on each point. Plants were incubated 48 hours in the dark at 18°C at night and 22°C at day, and then in the same temperature conditions with a 12-h photoperiod. Symptoms were scored at 13 and 15 dpi using the IMASCORE scale, scores of 1-3 corresponding to resistance and 4–6 to susceptibility phenotypes (Balesdent *et al*., 2001).

The OEC transformants obtained in this study were inoculated on the 207 genotypes from EPHICAS and semi-winter panels. Six individuals per genotype were inoculated with two control strains, JN2 and X83.51, and one transformant per *LmSTEE* gene. Inoculations were done as described above but on cotyledons of 10-day-old seedlings, with an incubation of 48 hours at room temperature and growth chamber condition set at 19 °C (night)/24°C(day), 16-h photoperiod, 90% humidity. Symptoms were scored at 10 and 13 dpi using the IMASCORE rating scale.

For promising interactions, a validation test was made. Inoculation was performed on six to twelve plants per genotype but using two OEC transformants per *LmSTEE* gene. Symptoms were scored at 10, 13 and 15 dpi using IMASCORE rating scale.

### Vector construction and fungal transformation

The expression of *LmSTEE* genes was induced during cotyledons colonization using the promoter of *AvrLm4-7*, an early effector gene known to be up-regulated at 7 days post infection of cotyledons (Parlange *et al*., 2009).

The promoter of the *AvrLm4-7* gene was cloned into the pPZPNat1 vector as described in Jiquel *et al*. (2021). The six new *LmSTEE* genes were amplified from their START codon to their terminator region using primers described in Table S6. The amplicons were then digested by *Eco*RI and *Xho*I or *Sal*I and *Xho*I and ligated into pPZPNat1_AvrLm4-7 vector. Because of the presence of *Xho*I restriction site in *LmSTEE5765* terminator, this gene was cloned with *LmSTEE833* terminator. In addition, *LmSTEE4385* was inserted into pPZPNat1_AvrLm4-7 vector using Gibson assembly (GeneArt™ Gibson Assembly HiFi Cloning Kit, Thermo Fisher Scientific, USA) according to manufacturer’s recommendations.

The plasmids were introduced into *Agrobacterium tumefaciens* C58 pGV2260 by electroporation at 2.5 kV, 200 Ohm and 25 μF. *L. maculans* isolate X83.51 was transformed with the resulting *A. tumefaciens* following the transformation protocol described by Idnurm *et al*. (2017). X83.51 isolate is virulent against most known *Rlm* genes except *Rlm5*, *Rlm10* and *Rlm14* allowing the screening of a large panel of *B. napus* genotypes without interference with known *AvrLm-Rlm* interactions.

Selection of positive transformants was made using nourseothricin (50 μg.ml-1) and cefotaxime (50 μg.ml-1). Positive transformants were grown on V8 with antibiotics and sporulated for isolation of a single pycnidiospore.

X83.51 was also transformed with pPZPNat1_AvrLm4-7:LmSTEE98 and pPZPNat1_AvrLm4- 7:LmSTEE6826, previously constructed by Jiquel *et al*. (2021, 2022), following the same protocol.

Fungal transformants were checked for their growth *in vitro* using V8 agar medium and for their pathogenicity by inoculation on a susceptible cultivar, Darmor. Finally, for each selected transformant, the correct insertion of pPZPNat1_AvrLm4-7 in frame with the corresponding *LmSTEE* gene was verified by PCR and sequencing on pycnidiospores harvested in sterile water.

### RNA manipulation and quantitative reverse transcription (qRT)-PCR

*LmSTEE* gene expression at 7dpi was quantified as described by Jiquel *et al*. (2021). Total RNA was extracted from inoculated cotyledons 7dpi on two biological replicates. We generated cDNA using oligo-dT-primed reverse transcription with the PowerScript reverse transcriptase (Clontech, Palo Alto, CA, USA), according to the manufacturer’s protocol. qRT-PCR was performed on two technical replicates as described by Fudal *et al*. (2007) with the primers indicated in Table S6. *Actin* was used as a constitutively expressed reference gene, and levels of *EF1α* expression relative to actin expression were used as control.

### Bioinformatics and statistical analyses

*L. maculans* genomic analyses were performed using JN3 genome assembly and annotations (Dutreux *et al*., 2018, https://bioinfo.bioger.inrae.fr/myGenomeBrowser?browse=1&portalname=Leptosphaeria_maculans_JN3&owner=bioinfobioger@inrae.fr&key=7d2D4XyQ). Effectors were predicted using EffectorP (Sperschneider *et al*., 2016). The presence of a signal peptide was predicted by SignalP (version 4.1, Nielsen, 2017) and the extracellular location by TargetP (version 1.1, Almagro Armenteros *et al*., 2019). The number of transmembrane domains was assigned using TMHMM tool (version 2.0, Möller *et al*., 2001). Structural prediction of effector was performed using Alphafold2 (ColabFold v1.5.5) with standard parameters (Mirdita *et al*., 2022).

Effector genes were assigned to “early” or “late” categories using the expression clusters defined by Gay *et al*. (2021). Effectors belonging to clusters 1 to 3 were defined as early and effectors from clusters 4 to 6 as late. The genomic location of effector genes was determined using Occultercut (Testa *et al*., 2016) on JN3 genome (Dutreux *et al*., 2018). The location was assigned as “border” when the gene was within a 10kb distance of an overlapping between AT-rich and GC-equilibrated isochores. TE were annotated by Grandaubert *et al*. (2014) on JN3 genome. The enrichment in chromatin methylation marks on effector genes was determined using ChiP-seq data generated by Soyer *et al*. (2021) during axenic growth of JN3.

The 205 previously published *L. maculans* genomes (Van de Wouw *et al*., 2024) were analysed for sequence variation content relative to the reference strain JN3. Raw reads were cleaned with Trimmomatic v0.32 (Bolger *et al*., 2014) and aligned with bwa-mem v 0.7.7 (Li and Durbin, 2009). Variant calling was performed with Freebayes v 0.9 (Garrison and Marth, 2012) before filtering with the same workflow previously established on other fungal species (Amezrou *et al*., 2024; Zhong *et al*., 2017). Raw reads were also assembled with a pipeline using a combination of Velvet (Zerbino and Birney, 2008), SOAPdenovo and SOAP GapCloser (Luo *et al*., 2012) as follows : (1) reads were trimmed at the first N, (2) contigs were generated with several k-mer values using SOAPdenovo, (3) several Velvet assemblies were built using several k-mer values and as the input the trimmed reads and all SOAPdenovo contigs considered as "long reads ", (4) the assembly that maximizes the criterion (N50*size of the assembly) was selected, (5) SOAP GapCloser was run on the selected assembly, and (6) contigs completely enclosed in other longer contigs were removed. The gene annotation was transferred from the JN3 reference to other strains using Liftoff (Shumate and Salzberg, 2021), enabling the variant content and detection of protein isoform for each genetic locus to be analyzed. The results are available on the web portal https://bioinfo.bioger.inrae.fr/portal/variant-explorer/. The protein sequence alignments were performed using the msa package (v 1.34.0) with the ClustalW method.

Statistical analyses were performed using RStudio (R 4.3.2). Chi² test, Kruskal-Wallis test and Anova were done using the package Rstatix (v 0.7.2).

## ACKNOWLEDGEMENTS

The authors wish to thank Marie-Hélène Balesdent for the obtention of X83.51 isolate, all members of the “Effectors and Pathogenesis of *Leptosphaeria maculans*” (EPLM) team of the BIOGER research unit, the greenhouse technician staff, the administrative supporting staff, the laboratory glassware staff and the bio-informatic staff of BIOGER and Innolea for their help with experimentations, analysis and administrative support. We thank the Genotoul bioinformatics platform Toulouse Occitanie (Bioinfo Genotoul, https://doi.org/10.15454/1.5572369328961167E12) and the BARIC workgroup (https://www.cesgo.org/catibaric/) for providing help, computing and storage resource. CR was funded by ANRT and Innolea (Cifre project no. 2022/1063). EPLM team benefits from the support of Saclay Plant Sciences-SPS (ANR-17-EUR-0007).

## REFERENCES

Almagro Armenteros, J.J., Salvatore, M., Emanuelsson, O., Winther, O., Von Heijne, G., Elofsson, A., Nielsen, H., 2019. Detecting sequence signals in targeting peptides using deep learning. Life Sci. Alliance 2, e201900429. 10.26508/lsa.201900429

Amezrou, R., Ducasse, A., Compain, J., Lapalu, N., Pitarch, A., Dupont, L., Confais, J., Goyeau, H., Kema, G.H.J., Croll, D., Amselem, J., Sanchez-Vallet, A., Marcel, T.C., 2024. Quantitative pathogenicity and host adaptation in a fungal plant pathogen revealed by whole-genome sequencing. Nat. Commun. 15, 1933. 10.1038/s41467-024-46191-1

Ansan-Melayah, D., Balesdent, M.-H., Buée, M., Rouxel, T., 1995. Genetic Characterization of *AvrLm1*, the First Avirulence Gene of *Leptosphaeria maculans*. Phytopathology 85, 1525. 10.1094/Phyto-85-1525

Balesdent, M.H., Attard, A., Ansan-Melayah, D., Delourme, R., Renard, M., Rouxel, T., 2001. Genetic Control and Host Range of Avirulence Toward *Brassica napus* Cultivars Quinta and Jet Neuf in *Leptosphaeria maculans*. Phytopathology® 91, 70–76. 10.1094/PHYTO.2001.91.1.70

Balesdent, M.H., Attard, A., Kühn, M.L., Rouxel, T., 2002. New Avirulence Genes in the Phytopathogenic Fungus *Leptosphaeria maculans*. Phytopathology® 92, 1122–1133. 10.1094/PHYTO.2002.92.10.1122

Balesdent, M.H., Barbetti, M.J., Li, H., Sivasithamparam, K., Gout, L., Rouxel, T., 2005. Analysis of *Leptosphaeria maculans* Race Structure in a Worldwide Collection of Isolates. Phytopathology® 95, 1061–1071. 10.1094/PHYTO-95-1061

Blondeau, K., Blaise, F., Graille, M., Kale, S.D., Linglin, J., Ollivier, B., Labarde, A., Lazar, N., Daverdin, G., Balesdent, M.-H., Choi, D.H.Y., Tyler, B.M., Rouxel, T., Van Tilbeurgh, H., Fudal, I., 2015. Crystal structure of the effector AvrLm4-7 of *Leptosphaeria maculans* reveals insights into its translocation into plant cells and recognition by resistance proteins. Plant J. 83, 610–624. 10.1111/tpj.12913

Bolger, A.M., Lohse, M., Usadel, B., 2014. Trimmomatic: a flexible trimmer for Illumina sequence data. Bioinformatics 30, 2114–2120. 10.1093/bioinformatics/btu170

Cambareri, E.B., Jensen, B.C., Schabtach, E., Selker, E.U., 1989. Repeat-induced G-C to A-T Mutations in *Neurospora*. Science 244, 1571–1575. 10.1126/science.2544994

Delourme, R., Chèvre, A.M., Brun, H., Rouxel, T., Balesdent, M.H., Dias, J.S., Salisbury, P., Renard, M., Rimmer, S.R., 2006. Major Gene and Polygenic Resistance to *Leptosphaeria maculans* in Oilseed Rape (*Brassica napus*). Eur. J. Plant Pathol. 114, 41–52. 10.1007/s10658-005-2108-9

Dong, S., Raffaele, S., Kamoun, S., 2015. The two-speed genomes of filamentous pathogens: waltz with plants. Curr. Opin. Genet. Dev. 35, 57–65. 10.1016/j.gde.2015.09.001

Dutreux, F., Da Silva, C., d’Agata, L., Couloux, A., Gay, E.J., Istace, B., Lapalu, N., Lemainque, A., Linglin, J., Noel, B., Wincker, P., Cruaud, C., Rouxel, T., Balesdent, M.-H., Aury, J.-M., 2018. De novo assembly and annotation of three *Leptosphaeria* genomes using Oxford Nanopore MinION sequencing. Sci. Data 5, 180235. 10.1038/sdata.2018.235

Fitt, B.D.L., Brun, H., Barbetti, M.J., Rimmer, S.R., 2006. World-Wide Importance of Phoma Stem Canker (*Leptosphaeria maculans* and *L. biglobosa*) on Oilseed Rape (*Brassica napus*). Eur. J. Plant Pathol. 114, 3–15. 10.1007/s10658-005-2233-5

Fitt, B.D.L., Hu, B.C., Li, Z.Q., Liu, S.Y., Lange, R.M., Kharbanda, P.D., Butterworth, M.H., White, R.P., 2008. Strategies to prevent spread of *Leptosphaeria maculans* (phoma stem canker) onto oilseed rape crops in China; costs and benefits. Plant Pathol. 57, 652–664. 10.1111/j.1365-3059.2008.01841.x

Flor, H.H., 1971. Current Status of the Gene-For-Gene Concept. Annu. Rev. Phytopathol. 9, 275–296. 10.1146/annurev.py.09.090171.001423

Fudal, I., Ross, S., Gout, L., Blaise, F., Kuhn, M.L., Eckert, M.R., Cattolico, L., Bernard-Samain, S., Balesdent, M.H., Rouxel, T., 2007. Heterochromatin-Like Regions as Ecological Niches for Avirulence Genes in the *Leptosphaeria maculans* Genome: Map-Based Cloning of *AvrLm6*. Mol. Plant-Microbe Interactions® 20, 459–470. 10.1094/MPMI-20-4-0459

Garrison, E., Marth, G., 2012. Haplotype-based variant detection from short-read sequencing. 10.48550/arXiv.1207.3907

Gay, E.J., Soyer, J.L., Lapalu, N., Linglin, J., Fudal, I., Da Silva, C., Wincker, P., Aury, J.-M., Cruaud, C., Levrel, A., Lemoine, J., Delourme, R., Rouxel, T., Balesdent, M.-H., 2021. Large-scale transcriptomics to dissect 2 years of the life of a fungal phytopathogen interacting with its host plant. BMC Biol. 19, 55. 10.1186/s12915-021-00989-3

Gervais, J., Plissonneau, C., Linglin, J., Meyer, M., Labadie, K., Cruaud, C., Fudal, I., Rouxel, T., Balesdent, M.-H., 2017. Different waves of effector genes with contrasted genomic location are expressed by *Leptosphaeria maculans* during cotyledon and stem colonization of oilseed rape: Hunting down *L. maculans* late effectors. Mol. Plant Pathol. 18, 1113–1126. 10.1111/mpp.12464

Gout, L., Kuhn, M.L., Vincenot, L., Bernard-Samain, S., Cattolico, L., Barbetti, M., Moreno-Rico, O., Balesdent, M.-H., Rouxel, T., 2007. Genome structure impacts molecular evolution at the *AvrLm1* avirulence locus of the plant pathogen *Leptosphaeria maculans*. Environ. Microbiol. 9, 2978–2992. 10.1111/j.1462-2920.2007.01408.x

Grandaubert, J., Lowe, R.G., Soyer, J.L., Schoch, C.L., Van de Wouw, A.P., Fudal, I., Robbertse, B., Lapalu, N., Links, M.G., Ollivier, B., Linglin, J., Barbe, V., Mangenot, S., Cruaud, C., Borhan, H., Howlett, B.J., Balesdent, M.-H., Rouxel, T., 2014. Transposable element-assisted evolution and adaptation to host plant within the *Leptosphaeria maculans*-*Leptosphaeria biglobosa* species complex of fungal pathogens. BMC Genomics 15, 891. 10.1186/1471-2164-15-891

Idnurm, A., Howlett, B.J., 2003. Analysis of loss of pathogenicity mutants reveals that repeat-induced point mutations can occur in the Dothideomycete *Leptosphaeria maculans*. Fungal Genet. Biol. 39, 31–37. 10.1016/S1087-1845(02)00588-1

Idnurm, A., Urquhart, A.S., Vummadi, D.R., Chang, S., Van De Wouw, A.P., López-Ruiz, F.J., 2017. Spontaneous and CRISPR/Cas9-induced mutation of the osmosensor histidine kinase of the canola pathogen *Leptosphaeria maculans*. Fungal Biol. Biotechnol. 4. 10.1186/s40694-017-0043-0

Irelan, J.T., Hagemann, A.T., Selker, E.U., 1994. High frequency repeat-induced point mutation (RIP) is not associated with efficient recombination in *Neurospora*. Genetics 138, 1093–1103. 10.1093/genetics/138.4.1093

Jiquel, A., Gay, E.J., Mas, J., George, P., Wagner, A., Fior, A., Faure, S., Balesdent, M., Rouxel, T., 2022. “Late” effectors from *Leptosphaeria maculans* as tools for identifying novel sources of resistance in *Brassica napus*. Plant Direct 6. 10.1002/pld3.435

Jiquel, A., Gervais, J., Geistodt-Kiener, A., Delourme, R., Gay, E.J., Ollivier, B., Fudal, I., Faure, S., Balesdent, M., Rouxel, T., 2021. A gene-for-gene interaction involving a ‘late’ effector contributes to quantitative resistance to the stem canker disease in *Brassica napus*. New Phytol. 231, 1510–1524. 10.1111/nph.17292

Jones, J.D.G., Dangl, J.L., 2006. The plant immune system. Nature 444, 323–329. 10.1038/nature05286

Li, H., Durbin, R., 2009. Fast and accurate short read alignment with Burrows–Wheeler transform. Bioinformatics 25, 1754–1760. 10.1093/bioinformatics/btp324

Lo Presti, L., Lanver, D., Schweizer, G., Tanaka, S., Liang, L., Tollot, M., Zuccaro, A., Reissmann, S., Kahmann, R., 2015. Fungal Effectors and Plant Susceptibility. Annu. Rev. Plant Biol. 66, 513–545. 10.1146/annurev-arplant-043014-114623

Luo, R., Liu, B., Xie, Y., Li, Z., Huang, W., Yuan, J., He, G., Chen, Y., Pan, Q., Liu, Yunjie, Tang, J., Wu, G., Zhang, H., Shi, Y., Liu, Yong, Yu, C., Wang, B., Lu, Y., Han, C., Cheung, D.W., Yiu, S.-M., Peng, S., Xiaoqian, Z., Liu, G., Liao, X., Li, Y., Yang, H., Wang, Jian, Lam, T.-W., Wang, Jun, 2012. SOAPdenovo2: an empirically improved memory-efficient short-read *de novo* assembler. Gigascience 1, 2047–217X-1–18. 10.1186/2047-217X-1-18

Mirdita, M., Schütze, K., Moriwaki, Y., Heo, L., Ovchinnikov, S., Steinegger, M., 2022. ColabFold: making protein folding accessible to all. Nat. Methods 19, 679–682. 10.1038/s41592-022-01488-1

Möller, S., Croning, M.D.R., Apweiler, R., 2001. Evaluation of methods for the prediction of membrane spanning regions. Bioinformatics 17, 646–653. 10.1093/bioinformatics/17.7.646

Neik, T.X., Amas, J., Barbetti, M., Edwards, D., Batley, J., 2020. Understanding Host–Pathogen Interactions in *Brassica napus* in the Omics Era. Plants 9, 1336. 10.3390/plants9101336

Neik, T.X., Ghanbarnia, K., Ollivier, B., Scheben, A., Severn-Ellis, A., Larkan, N.J., Haddadi, P., Fernando, D.W.G., Rouxel, T., Batley, J., Borhan, H.M., Balesdent, M., 2022. Two independent approaches converge to the cloning of a new *Leptosphaeria maculans* avirulence effector gene, *AvrLmS-Lep2*. Mol. Plant Pathol. 23, 733–748. 10.1111/mpp.13194

Nielsen, H., 2017. Predicting Secretory Proteins with SignalP, in: Kihara, D. (Ed.), Protein Function Prediction, Methods in Molecular Biology. Springer New York, New York, NY, pp. 59–73. 10.1007/978-1-4939-7015-5_6

Niks, R.E., Qi, X., Marcel, T.C., 2015. Quantitative Resistance to Biotrophic Filamentous Plant Pathogens: Concepts, Misconceptions, and Mechanisms. Annu. Rev. Phytopathol. 53, 445–470. 10.1146/annurev-phyto-080614-115928

Parlange, F., Daverdin, G., Fudal, I., Kuhn, M.-L., Balesdent, M.-H., Blaise, F., Grezes-Besset, B., Rouxel, T., 2009. *Leptosphaeria maculans* avirulence gene *AvrLm4-7* confers a dual recognition specificity by the *Rlm4* and *Rlm7* resistance genes of oilseed rape, and circumvents *Rlm4*-mediated recognition through a single amino acid change. Mol. Microbiol. 71, 851–863. 10.1111/j.1365-2958.2008.06547.x

Plissonneau, C., Daverdin, G., Ollivier, B., Blaise, F., Degrave, A., Fudal, I., Rouxel, T., Balesdent, M., 2016. A game of hide and seek between avirulence genes *AvrLm4-7* and *AvrLm3* in *Leptosphaeria maculans*. New Phytol. 209, 1613–1624. 10.1111/nph.13736

Rocafort, M., Fudal, I., Mesarich, C.H., 2020. Apoplastic effector proteins of plant-associated fungi and oomycetes. Curr. Opin. Plant Biol. 56, 9–19. 10.1016/j.pbi.2020.02.004

Rouxel, T., Balesdent, M.H., 2005. The stem canker (blackleg) fungus, *Leptosphaeria maculans* , enters the genomic era. Mol. Plant Pathol. 6, 225–241. 10.1111/j.1364-3703.2005.00282.x

Rouxel, T., Grandaubert, J., Hane, J.K., Hoede, C., van de Wouw, A.P., Couloux, A., Dominguez, V., Anthouard, V., Bally, P., Bourras, S., Cozijnsen, A.J., Ciuffetti, L.M., Degrave, A., Dilmaghani, A., Duret, L., Fudal, I., Goodwin, S.B., Gout, L., Glaser, N., Linglin, J., Kema, G.H.J., Lapalu, N., Lawrence, C.B., May, K., Meyer, M., Ollivier, B., Poulain, J., Schoch, C.L., Simon, A., Spatafora, J.W., Stachowiak, A., Turgeon, B.G., Tyler, B.M., Vincent, D., Weissenbach, J., Amselem, J., Quesneville, H., Oliver, R.P., Wincker, P., Balesdent, M.-H., Howlett, B.J., 2011. Effector diversification within compartments of the *Leptosphaeria maculans* genome affected by Repeat-Induced Point mutations. Nat. Commun. 2, 202. 10.1038/ncomms1189

Rouxel, T., Penaud, A., Pinochet, X., Brun, H., Gout, L., Delourme, R., Schmit, J., Balesdent, M.-H., 2003. A 10-year survey of populations of *Leptosphaeria maculans* in France indicates a rapid adaptation towards the *Rlm1* resistance gene of oilseed rape.

Salisbury, P., Ballinger, D., Wratten, N., Plummer, K., Howlett, B., 1995. Blackleg disease on oilseed *Brassica* in Australia: a review. Aust. J. Exp. Agric. 35, 665. 10.1071/EA9950665

Sánchez-Vallet, A., Fouché, S., Fudal, I., Hartmann, F.E., Soyer, J.L., Tellier, A., Croll, D., 2018. The Genome Biology of Effector Gene Evolution in Filamentous Plant Pathogens. Annu. Rev. Phytopathol. 56, 21–40. 10.1146/annurev-phyto-080516-035303

Shumate, A., Salzberg, S.L., 2021. Liftoff: accurate mapping of gene annotations. Bioinformatics 37, 1639–1643. 10.1093/bioinformatics/btaa1016

Soyer, J.L., Clairet, C., Gay, E.J., Lapalu, N., Rouxel, T., Stukenbrock, E.H., Fudal, I., 2021. Genome-wide mapping of histone modifications during axenic growth in two species of *Leptosphaeria maculans* showing contrasting genomic organization. Chromosome Res. 29, 219–236. 10.1007/s10577-021-09658-1

Sperschneider, J., Gardiner, D.M., Dodds, P.N., Tini, F., Covarelli, L., Singh, K.B., Manners, J.M., Taylor, J.M., 2016. EFFECTOR P: predicting fungal effector proteins from secretomes using machine learning. New Phytol. 210, 743–761. 10.1111/nph.13794

Sprague, S.J., Marcroft, S.J., Hayden, H.L., Howlett, B.J., 2006. Major Gene Resistance to Blackleg in *Brassica napus* Overcome Within Three Years of Commercial Production in Southeastern Australia. Plant Dis. 90, 190–198. 10.1094/PD-90-0190

St.Clair, D.A., 2010. Quantitative Disease Resistance and Quantitative Resistance Loci in Breeding. Annu. Rev. Phytopathol. 48, 247–268. 10.1146/annurev-phyto-080508-081904

Testa, A.C., Oliver, R.P., Hane, J.K., 2016. OcculterCut: A Comprehensive Survey of AT-Rich Regions in Fungal Genomes. Genome Biol. Evol. 8, 2044–2064. 10.1093/gbe/evw121

Van De Wouw, A.P., Elliott, C.E., Popa, K.M., Idnurm, A., 2019. Analysis of Repeat Induced Point (RIP) Mutations in *Leptosphaeria maculans* Indicates Variability in the RIP Process Between Fungal Species. Genetics 211, 89–104. 10.1534/genetics.118.301712

Van de Wouw, A.P., Scanlan, J.L., Al-Mamun, H.A., Balesdent, M., Bousset, L., Burketová, L., Del Rio Mendoza, L., Fernando, W.G.D., Franke, C., Howlett, B.J., Huang, Y., Jones, E.E., Koopmann, B., Lob, S., Mirabadi, A.Z., Nugent, B.C., Peng, G., Rossi, F.R., Schreuder, H., Tabone, A.R., Van Coller, G.J., Batley, J., Idnurm, A., 2024. A new set of international *Leptosphaeria maculans* isolates as a resource for elucidation of the basis and evolution of blackleg disease on *Brassica napus*. Plant Pathol. 73, 170–185. 10.1111/ppa.13801

Vasquez-Teuber, P., Rouxel, T., Mason, A.S., Soyer, J.L., 2024. Breeding and management of major resistance genes to stem canker/blackleg in *Brassica* crops. Theor. Appl. Genet. 137, 192. 10.1007/s00122-024-04641-w

West, J.S., Kharbanda, P.D., Barbetti, M.J., Fitt, B.D.L., 2001. Epidemiology and management of *Leptosphaeria maculans* (phoma stem canker) on oilseed rape in Australia, Canada and Europe: Epidemiology of *L. maculans* on rapeseed. Plant Pathol. 50, 10–27. 10.1046/j.1365-3059.2001.00546.x

Yang, C., Zou, Z., Fernando, W.G.D., 2021. The Effect of Temperature on the Hypersensitive Response (HR) in the *Brassica napus–Leptosphaeria maculans* Pathosystem. Plants 10, 843. 10.3390/plants10050843

Zerbino, D.R., Birney, E., 2008. Velvet: Algorithms for de novo short read assembly using de Bruijn graphs. Genome Res. 18, 821–829. 10.1101/gr.074492.107

Zhong, Z., Marcel, T.C., Hartmann, F.E., Ma, X., Plissonneau, C., Zala, M., Ducasse, A., Confais, J., Compain, J., Lapalu, N., Amselem, J., McDonald, B.A., Croll, D., Palma-Guerrero, J., 2017. A small secreted protein in *Zymoseptoria tritici* is responsible for avirulence on wheat cultivars carrying the *Stb6* resistance gene. New Phytol. 214, 619–631. 10.1111/nph.14434

