## Supplemental FIgures and Tables for "Conserved “late” effector genes from *Leptosphaeria maculans* inducing gene-for-gene quantitative resistance in *Brassica napus* semi-winter genotypes"

SUPPLEMENTARY FIGURES AND TABLES

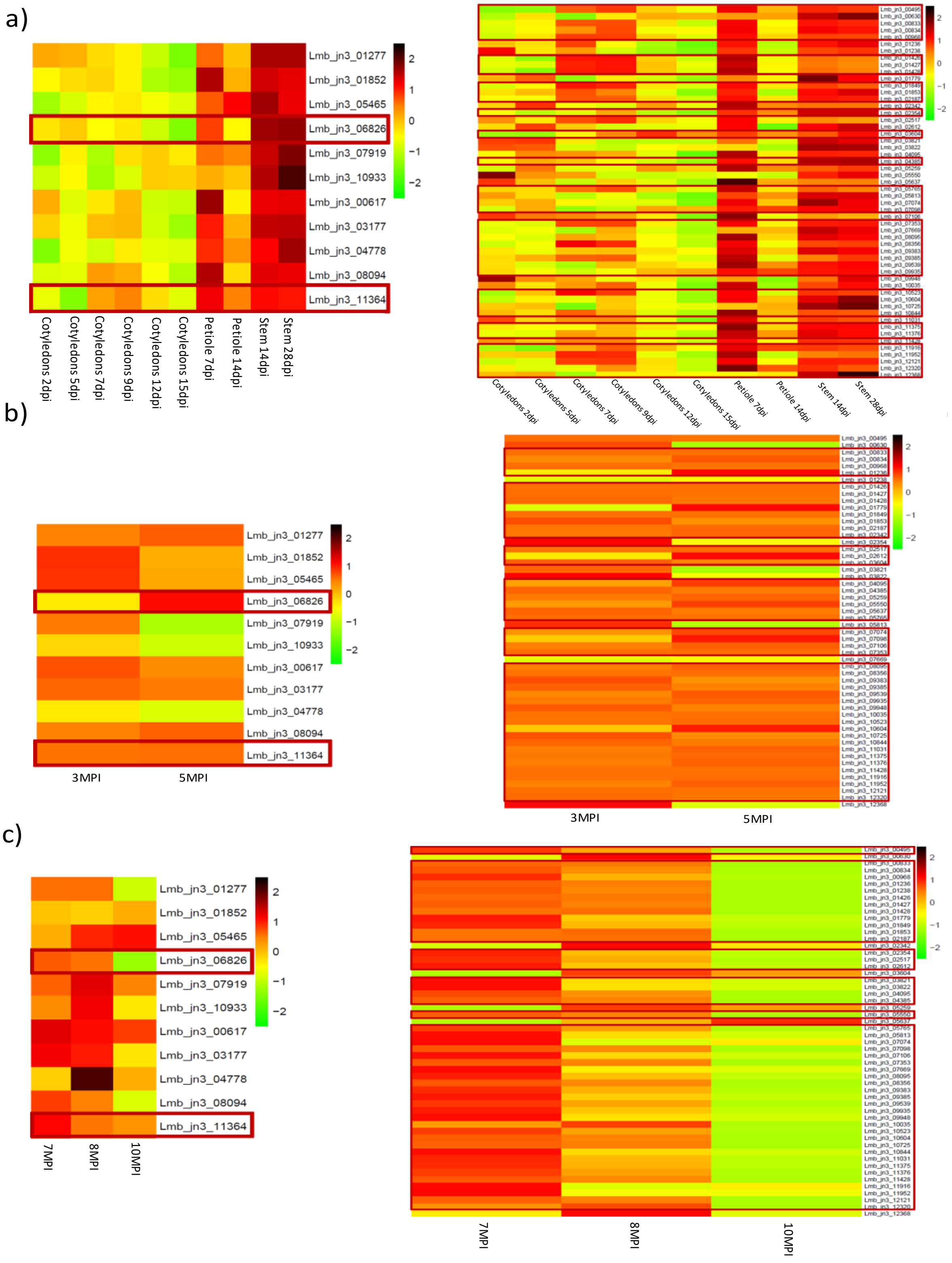

Figure S1: In planta expression data of a selection of late effector genes of *Leptosphaeria maculans* in controlled conditions (a) and in field conditions during leaf infection (b) and stem infection (c). Left graphics present the expression of the 11 late effectors studied in Jiquel *et al*. (2021, 2022) and right graphics the expression of the candidate late effector genes selected in the present study. The expressions of *LmSTEE6826* (*Lmb_jn3_06826*) and *LmSTEE98* (*Lmb_jn3_11364*) are framed in red such as the effector genes presenting the same profile of expression in the same conditions. DPI : Day Post Infection, MPI : Month Post Infection

Table S1: Characteristics of the 151 effector genes of *Leptosphaeria maculans* analyzed in this study. The effector genes were predicted by Dutreux *et al*. (2018) , the expression clusters were defined by Gay *et al*. (2021). Genomic location in AT-rich (AT), GC-equilibrated (GC) isochores or border between these two locations were defined by Dutreux *et al*. (2018). Transposable elements present 4kbp upstream and downstream of the gene were annotated by Grandaubert *et al.* (2014). Enrichment in chromatin methylation marks during axenic growth was detected by Soyer *et al*. (2021) using ChIP-seq. Peptide signals were predicted by SignalP (Nielsen, 2017), extracellular location by TargetP (Almagro Armenteros *et al.,* 2019) and transmembrane domains by TMHMM tool (Möller *et al.,* 2001). Protein homologues were identified by BLASTp using the NCBI nr database, quality of the structural prediction using AlphaFold (ColabFold v1.5.5, Mirdita *et al*., 2022) and close structural analog according to the Dali database.

| Gene ID [a] | *L. maculans* characterized effectors | Assignation to the reference expression clusters [b] | Genomic location [c] | Presence of transposable element close to the gene [d] | Transposable elements close to the gene [d] | Enrichment in chromatin methylation marks [e] | Presence of a peptide signal [f] | Extracellular location [g] | Transmembran domains [h] | Protein size | Number of cysteines | Existence of homologs outside *Leptosphaeria* genus [i] | Five first protein homologs [i] | Structural prediction [j] | Five first structural analogues [k] |
|---|---|---|---|---|---|---|---|---|---|---|---|---|---|---|---|
| *Lmb_jn3_00001* | *AvrLm3* (Plissonneau et al., 2016) | Cluster 2 | AT | YES | RLG_Rolly (Overlapping) RLG_Olly (300bp) RLC_Pholy (1000bp) | H3K9me3 | YES | YES | <=1 TM | 161 | 11 | YES | Hypothetical protein BFJ63_vAg18418 [*Fusarium oxysporum* f. sp.*narcissi*] Ecp11-1 [*Fulvia fulva*] Chain A Extracellular protein 11-1 [*Fulvia fulva*] | Low | |
| *Lmb_jn3_00187* | - | Cluster 2 | Border | YES | RLG_Brawly (400bp) Lmac_Recon_92 _20 (700bp) RLG_Olly (1100bp) RLC_Pholy (3200bp) | H3K27me3 | YES | YES | <=1 TM | 103 | 11 | YES | Unnamed protein product [*Zymoseptoria tritici* ST99CH_3D7] Ecp43-1 [*Fulvia fulva*] Uncharacterized protein CLAFUR5_14680 [*Fulvia fulva*] Hypothetical protein TI39_contig395g 00002 [*Zymoseptoria brevis*] | Very Low | |
| *Lmb_jn3_00284* | (Gervais et al., 2017) | Cluster 4 | GC | NO | | H3K27me3 | YES | YES | <=1 TM | 117 | 4 | YES | Hypothetical protein EsH8_VIII_00017 0 [*Colletotrichum jinshuiense*] Hypothetical protein CMUS01_12090 [*Colletotrichum musicola*] Uncharacterized protein CGCA056_v0056 64 [*Colletotrichum aenigma*] Uncharacterized protein | Low | |

| | | | | | | | | | | | | | | | |
|---|---|---|---|---|---|---|---|---|---|---|---|---|---|---|---|
| | | | | | | | | | | | | | CGGC5_v011370 [*Colletotrichum fructicola* Nara gc5]<br>Hypothetical protein CSOJ01_15032 [*Colletotrichum sojae*] | | |
| *Lmb_jn3_00461* | - | Cluster 2 | Border | YES | RLC_Pholy (Overlapping)<br>DTT_Finwe_2 (200bp)<br>RLx_Jolly (250bp)<br>DTT_Finwe_1 (1000bp)<br>DTT_Finwe_3 (1100bp)<br>Lmac_Recon_92_20 (1300bp)<br>RLG_Polly (1400bp)<br>Lmac_Grouper_2395_3 (1600bp)<br>RLG_Polly (2000bp) | H3K9me3 | YES | NO | <=1 TM | 148 | 6 | YES | Hypothetical protein FOMG_19260 [*Fusarium oxysporum* f. sp. *melonis* 26406]<br>Ecp7(P20) [*Fusarium agapanthi*]<br>Hypothetical protein BFJ65_g17658 [*Fusarium oxysporum* f. sp. *cepae*]<br>Hypothetical protein HZS61_008393 [*Fusarium oxysporum* f. sp. *conglutinans*]<br>Hypothetical protein FOVG_19456 [*Fusarium oxysporum* f. sp. *pisi* HDV247] | Good | Chitinase, Lysozyme<br>Cell Wall-Binding Endopeptinase-Related Protein<br>Putative L, D Transpeptidase YKUD<br>Chitin Elicitor-Binding Protein<br>Chitinase A |
| *Lmb_jn3_00495* | - | Cluster 4 | GC | YES | RLG_Olly (Overlapping)<br>DTx_Olwe (200bp)<br>RLG_Rolly (500bp) | H3K9me3 | YES | YES | <=1 TM | 124 | 7 | NO | | Very low | |
| *Lmb_jn3_00561* | - | Cluster 6 | GC | NO | | H3K27me3 | YES | YES | <=1 TM | 219 | 4 | YES | Hypothetical protein J1614_012156 [*Plenodomus biglobosus*]<br>Hypothetical protein T440DRAFT_408842 [*Plenodomus tracheiphilus IPT5*]<br>Hypothetical protein CC77DRAFT_849095 [*Alternaria alternata*]<br>Hypothetical protein | Good | DNA Repair Protein XRCC1<br>Beta-Xylanase |

| | | | | | | | | | | | | | | | |
|---|---|---|---|---|---|---|---|---|---|---|---|---|---|---|---|
| | | | | | | | | | | | | | NX059_007160 [*Plenodomus lindquistii*] Hypothetical protein AG0111_0g10857 [*Alternaria gaisen*] | | |
| *Lmb_jn3_00617* | LmSTEE78 (Jiquel et al., 2021) | Cluster 6 | GC | NO | | H3K4me2 | NO | NO | <=1 TM | 324 | 3 | YES | Hypothetical protein J1614_008571 [*Plenodomus biglobosus*] Hypothetical protein NX059_001003 [*Plenodomus lindquistii*] Glycoside hydrolase family 131 protein [*Plenodomus tracheiphilus*] Glycoside hydrolase family 131 protein [*Cucurbitaria berberidis* CBS 394.84 IPT5] Hypothetical protein N0V83_001597 [*Neocucurbitaria cava*] | Good | Beta_Glucanase Putative Uncharacterized Protein Glucuronan Lyase A Alginate Lyase PA1167 |
| *Lmb_jn3_00630* | - | Cluster 5 | GC | NO | | H3K4me2 + H3K27me3 | YES | YES | <=1 TM | 512 | 7 | YES | Hypothetical protein AA0111_g5834 [*Alternaria arborescens*] Hypothetical protein AA0121_g9039 [*Alternaria tenuissima*] Hypothetical protein AA0116_g12109 [*Alternaria tenuissima*] Hypothetical protein AA0118_g10476 [*Alternaria tenuissima*] Hypothetical protein AA0117_g7256 [*Alternaria alternata*] | Medium | |

| | | | | | | | | | | | | | | | |
|---|---|---|---|---|---|---|---|---|---|---|---|---|---|---|---|
| *Lmb_jn3_00833* | Present study | Cluster 4 | GC | YES | DTx_Gimli (1500bp) | H3K27me3 | YES | YES | <=1 TM | 99 | 9 | NO | | Good | KP6 Killer Toxin Subunit Alpha |
| *Lmb_jn3_00834* | - | Cluster 4 | GC | YES | DTx_Gimli (800bp) | H3K27me3 | NO | YES | <=1 TM | 61 | 4 | NO | | Low | |
| *Lmb_jn3_00910* | - | Cluster 6 | GC | YES | RLC_Zolly-2 (150bp) DTx_Gimli (1700bp) | H3K27me3 | YES | YES | <=1 TM | 217 | 0 | YES | Hypothetical protein J1614_008884 [*Plenodomus biglobosus*] Hypothetical protein T440DRAFT_472526 [*Plenodomus tracheiphilus IPT5*] Uncharacterized protein H3K4ME260DRAFT_365146 [*Cucurbitaria berberidis* CBS 394.84] Hypothetical protein NX059_001291 [*Plenodomus lindquistii*] Uncharacterized protein BKA58DRAFT_19842 [*Alternaria rosae*] | Very Low | |
| *Lmb_jn3_00912* | - | Cluster 6 | GC | YES | RLC_Zolly-2 (1200bp) | H3K27me3 | YES | YES | <=1 TM | 192 | 7 | YES | Hypothetical protein T440DRAFT_407557 [*Plenodomus tracheiphilus IPT5*] Hypothetical protein J1614_008887 [*Plenodomus biglobosus*] Hypothetical protein EK21DRAFT_74121 [*Setomelanomma holmii*] Uncharacterized protein H3K4ME260DRAFT_280079 [*Cucurbitaria berberidis* CBS 394.84] Hypothetical protein BS50DRAFT_605254 [*Corynespora* | Low | |

| | | | | | | | | | | | | | | | |
|---|---|---|---|---|---|---|---|---|---|---|---|---|---|---|---|
| | | | | | | | | | | | | | *cassiicola Philippines*] | | |
| *Lmb_jn3_00919* | - | Cluster 2 | Border | YES | DTx_Gimli (1500bp) DTx_Gimli (2000bp) | H3K27me3 | YES | YES | <=1 TM | 120 | 4 | YES | Hypothetical protein FDECE_10468 [*Fusarium decemcellulare*] Hypothetical protein H9Q73_014274 [*Fusarium xylarioides*] | Very Low | |
| *Lmb_jn3_00968* | - | Cluster 4 | GC | YES | RLG_Olly (2000bp) | H3K27me3 | YES | YES | <=1 TM | 102 | 10 | NO | | Low | |
| *Lmb_jn3_01236* | - | Cluster 5 | Border | YES | RLG_Brawly (100bp) RLC_Zolly-1 (200bp) RLC_Zolly-2 (500bp) | H3K9me3 + H3K27me3 | YES | YES | <=1 TM | 177 | 4 | YES | Hypothetical protein J1614_008082 [*Plenodomus biglobosus*] Hypothetical protein BKA66DRAFT_513830 [*Pyrenochaeta* sp. MPI-SDFR-AT-0127] Hypothetical protein BDV26DRAFT_289816 [*Aspergillus bertholletiae*] Hypothetical protein F4825DRAFT_476002 [*Nemania diffusa*] Hypothetical protein BS50DRAFT_596213 [*Corynespora cassiicola Philippines*] | Good | Ribonuclease U2 Ribonuclease F1 Ribonuclease T1 isozyme Protein (Ribonuclease T1) Ribonuclease MS |
| *Lmb_jn3_01238* | - | Cluster 5 | Border | NO | | H3K27me3 | YES | YES | <=1 TM | 103 | 6 | YES | Hypothetical protein BDV26DRAFT_289818 [*Aspergillus bertholletiae*] Hypothetical protein BKA66DRAFT_423003 [*Pyrenochaeta* sp. MPI-SDFR-AT-0127] Hypothetical protein NM208_g9258 [*Fusarium decemcellulare*] | Good | Carbohydrate Binding Protein Ciona Bentagamma-Crystallin Clostrillin Development Specific Protein S Beta-Crystallin B3 |

| | | | | | | | | | | | | | | |
|---|---|---|---|---|---|---|---|---|---|---|---|---|---|---|
| | | | | | | | | | | | | Hypothetical protein FDECE_1653 [*Fusarium decemcellulare*] Hypothetical protein FOVG_18305 [*Fusarium oxysporum* f. sp. *pisi* HDV247] | | |
| *Lmb_jn3_01277* | (Jiquel et al., 2022) | Cluster 5 | GC | YES | DTx_Gimli (800bp) DTx_Gimli (1200bp) | H3K27me3 | YES | YES | <=1 TM | 109 | 9 | YES | Hypothetical protein T440DRAFT_471373 [*Plenodomus tracheiphilus IPT5*] Uncharacterized protein H3K4ME260DRAFT_341652 [*Cucurbitaria berberidis* CBS 394.84] Hypothetical protein OPT61_g7390 [*Boeremia exigua*] Hypothetical protein SVAN01_07742 [*Stagonosporopsis vannaccii*] Hypothetical protein DE146DRAFT_760439 [*Phaeosphaeria* sp. MPI-PUGE-AT-0046c] | Medium | |
| *Lmb_jn3_01426* | - | Cluster 4 | GC | YES | RLG_Olly (100bp) | H3K27me3 | YES | YES | <=1 TM | 140 | 6 | YES | Hypothetical protein TW65_03826 [*Stemphylium lycopersici*] Hypothetical protein J1614_004295 [*Plenodomus biglobosus*] Hypothetical protein MPH_00188 [*Macrophomina phaseolina* MS6] Hypothetical protein PtrM4_116000 | Good | Avirulence Effector AvrLm4-7 Ribulose-1,5-Bisphosphate Carboxylase/Oxygenase L 50S Ribosomal Protein L31 Hydrogenase Maturation Protein HYPF 16S Ribosomal RNA |

| | | | | | | | | | | | | | | | |
|---|---|---|---|---|---|---|---|---|---|---|---|---|---|---|---|
| | | | | | | | | | | | | | [*Pyrenophora tritici-repentis*] Predicted protein [*Pyrenophora tritici-repentis* Pt-1C-BFP] | | |
| *Lmb_jn3_01427* | - | Cluster 4 | GC | YES | RLG_Olly (1400bp) | H3K27me3 | YES | YES | <=1 TM | 138 | 8 | YES | Hypothetical protein TW65_03827 [*Stemphylium lycopersici*] Hypothetical protein J1614_004296 [*Plenodomus biglobosus*] Hypothetical protein MPH_00189 [*Macrophomina phaseolina* MS6] Hypothetical protein MPH_11524 [*Macrophomina phaseolina* MS6] Hypothetical protein PTMSG1_10059 [*Pyrenophora teres* f. *maculata*] | Good | Avirulence effector AvrLm4-7 Probable SECDF Protein-Export Membrane Protein N Utilization Substance Protein B Ribulose-1,5-Bisphosphate Carboxylase/Oxygenase L Microtubule-Associated Protein 1A/1B Light Chain |
| *Lmb_jn3_01428* | - | Cluster 4 | GC | YES | RLG_Olly (2800bp) DTx_Gimli (3500bp) DTx_Gimli (3700bp) | H3K27me3 | YES | YES | <=1 TM | 131 | 8 | YES | Pathogenesis related protein [*Stemphylium lycopersici*] Hypothetical protein MPH_00190 [*Macrophomina phaseolina* MS6] Hypothetical protein BS50DRAFT_640827 [*Corynespora cassiicola Philippines*] Hypothetical protein MPH_11525 [*Macrophomina phaseolina* MS6] Hypothetical protein B0J12DRAFT_586512 [*Macrophomina phaseolina*] | Good | Avirulence effector AvrLm4-7 Probable SECDF Protein-Export Membrane Protein Elongation Factor 1-Beta Metal Homeostasis factor ATX1 |

| | | | | | | | | | | | | | | |
|---|---|---|---|---|---|---|---|---|---|---|---|---|---|---|
| *Lmb_jn3_01748* | - | Cluster 2 | Border | YES | RLG_Brawly (Overlapping) RLx_Jolly (100bp) DTM_Ingwe (450bp) RLG_Olly (900bp) RLG_Olly (900bp) RLG_Olly (1250bp) Lmac_Grouper_595_20 (1650bp) RLC_Pholy (2250bp) RLG_Olly (3300bp) Lmac_Grouper_595_20 (4000bp) | H3K27me3 | YES | YES | <=1 TM | 130 | 5 | NO | | Very Low |
| *Lmb_jn3_01779* | - | Cluster 5 | GC | NO | | H3K4me2 + H3K27me3 | NO | YES | <=1 TM | 58 | 1 | NO | | Very Low |
| *Lmb_jn3_01849* | - | Cluster 4 | Border | YES | Lmac_Grouper_284_9 (Overlapping) RLG_Rolly (50bp) RLx_Ayoly (400bp) RLG_Olly (600bp) RLx_Ayoly (800bp) RLG_Brawly (2100bp) DTx_Valwe (2900bp) | H3K27me3 | YES | YES | <=1 TM | 103 | 11 | NO | Hypothetical protein J1614_011012 [*Plenodomus biglobosus*] | Very Low |
| *Lmb_jn3_01851* | - | Cluster 2 | AT | YES | LmTelo1 (150bp) RLx_Jolly (200bp) RLG_Dolly (600bp) Lmac_Grouper_1227_7 (1600bp) RLC_Pholy (1650bp) | H3K27Me3 | YES | YES | <=1 TM | 92 | 9 | NO | | Medium |
| *Lmb_jn3_01852* | (Jiquel et al., 2022) | Cluster 5 | Border | YES | DTx_Gimli (300bp) DTx_Gimli (700bp) DTx_Gimli (1300bp) RLC_Zolly-1 (1400bp) RLG_Olly (1700bp) | H3K27me3 | YES | YES | <=1 TM | 89 | 8 | NO | | Low |
| *Lmb_jn3_01853* | Present study | Cluster 4 | Border | YES | DTx_Gimli (100bp) DTx_Gimli (300bp) DTx_Gimli (1700bp) RLC_Zolly-1 (2500bp) | H3K27me3 | YES | YES | <=1 TM | 110 | 10 | NO | | Low |

| | | | | | | | | | | | | | | | |
|---|---|---|---|---|---|---|---|---|---|---|---|---|---|---|---|
| | | | | | RLG_Olly (2700bp) | | | | | | | | | | |
| *Lmb_jn3_02093* | - | Cluster 2 | AT | YES | DTM_Lenwe (50bp) LmTelo1 (1100bp) Lmac_Grouper_1227_7 (1800bp) RLx_Jolly (1850bp) LmTelo2 (2200bp) RLG_Olly (2200bp) LmTelo1 (2800bp) RLG_Olly (3000bp) | H3K9me3 + H3K27me3 | YES | YES | <=1 TM | 145 | 1 | NO | | Low | |
| *Lmb_jn3_02094* | - | Cluster 2 | AT | YES | Lmac_Grouper_1227_7 (Overlapping) LmTelo2 (200bp) RLG_Olly (200bp) LmTelo1 (750bp) RLG_Olly (950bp) DTM_Lenwe (2000bp) LmTelo1 (2150bp) RLx_Jolly (2800bp) RLC_Pholy (3800bp) | H3K9me3 + H3K27me3 | YES | YES | <=1 TM | 108 | 10 | YES | Hypothetical protein FOWG_17837 [*Fusarium oxysporum* f. sp. *lycopersici* MN25] Hypothetical protein HZS61_006526 [*Fusarium oxysporum* f. sp. *conglutinans*] Hypothetical protein FOFC_13916 [*Fusarium oxysporum*] Hypothetical protein QL093DRAFT_2508277 [*Fusarium oxysporum*] Hypothetical protein V3481_006880 [*Fusarium oxysporum* f. sp. *vasinfectum*] | Medium | |
| *Lmb_jn3_02187* | Present study | Cluster 4 | GC | YES | RLG_Brawly (Overlapping) | H3K27me3 | YES | YES | <=1 TM | 107 | 5 | NO | | Low | |
| *Lmb_jn3_02342* | - | Cluster 4 | GC | YES | DTx_Valwe (1500bp) DTx_Valwe (1600bp) | H3K27me3 | YES | YES | <=1 TM | 258 | 10 | YES | Hypothetical protein J1614_010155 [*Plenodomus biglobosus*] | Good | Pectate Lyase Endo-Pectate Lyase Alpha-1,3-Glucanasa |

| | | | | | | | | | | | | | | | |
|---|---|---|---|---|---|---|---|---|---|---|---|---|---|---|---|
| | | | | | | | | | | | | | Polysaccharide lyase family 3 protein [*Plenodomus tracheiphilus* IPT5]<br>Hypothetical protein N0V83_004407 [*Neocucurbitaria cava*]<br>Pectate lyase-domain-containing protein [*Pyrenochaeta* sp. MPI-SDFR-AT-0127]<br>Pectate lyase-domain-containing protein [*Paraphoma chrysanthemicola*] | | Endopolygalacturonase Polygalacturonase II |
| *Lmb_jn3_02354* | - | Cluster 4 | GC | NO | | H3K27me3 | NO | YES | <=1 TM | 140 | 1 | YES | Hypothetical protein T440DRAFT_320149 [*Plenodomus tracheiphilus* IPT5] | Low | |
| *Lmb_jn3_02425* | - | Cluster 2 | Border | YES | Lmac_Grouper_284_9 (Overlapping)<br>Lmac_Grouper_2395_3 (100bp)<br>RLG_Olly (700bp) | H3K9me3 | YES | YES | <=1 TM | 132 | 10 | YES | Hypothetical protein BBP40_010151 [*Aspergillus hancockii*]<br>Hypothetical protein BO97DRAFT_420667 [*Aspergillus homomorphus* CBS 101889]<br>Hypothetical protein ASPGLDRAFT_30381 [*Aspergillus glaucus* CBS 516.65]<br>Uncharacterized protein N7471_000691 [*Penicillium samsonianum*]<br>Hypothetical protein PITC_068580 [*Penicillium italicum*] | Medium | |
| *Lmb_jn3_02517* | - | Cluster 4 | GC | YES | DTF_Elwe (400bp) | H3K27me3 | YES | YES | <=1 TM | 118 | 7 | NO | | Low | |

| | | | | | | | | | | | | | | | |
|---|---|---|---|---|---|---|---|---|---|---|---|---|---|---|---|
| *Lmb_jn3_02612* | - | Cluster 5 | Border | YES | DTx_Gimli (900bp) RLx_Ayoly (1100bp) | H3K27me3 | NO | YES | <=1 TM | 157 | 7 | YES | Hypothetical protein CSIM01_12189 [*Colletotrichum simmondsii*] Secreted in xylem 5 [*Colletotrichum scovillei*] Hypothetical protein BDP67DRAFT_470684 [*Colletotrichum lupini*] Hypothetical protein CSAL01_09201 [*Colletotrichum salicis*] Uncharacterized protein CCOS01_01482 [*Colletotrichum costaricense*] | Good | KP6 Killer Toxin Subunit Alpha Protein (Toxin) Protein (Toxin) |
| *Lmb_jn3_02967* | - | Cluster 2 | GC | YES | RLG_Olly (Overlapping) DTT_Finwe_1 (150bp) DTM_Sahana (1500bp) | NO | YES | YES | <=1 TM | 75 | 7 | NO | | Medium | |
| *Lmb_jn3_03177* | *LmSTEE1* (Jiquel et al., 2021) | Cluster 5 | GC | YES | DTx_Gimli (100bp) RLG_Dolly (500bp) | H3K9me3 + H3K27me3 | YES | YES | <=1 TM | 81 | 10 | YES | Hypothetical protein J1614_008687 [*Plenodomus biglobosus*] Hypothetical protein IFR05_006425 [*Cadophora* sp. M221] Hypothetical protein DL95DRAFT_467696 [*Leptodontidium* sp. 2 PMI_412] Hypothetical protein BKA61DRAFT_735650 [*Leptodontidium* sp. MPI-SDFR-AT-0119] Hypothetical protein ABOM_005030 [*Aspergillus bombycis*] | Good | None |

| | | | | | | | | | | | | | | | |
|---|---|---|---|---|---|---|---|---|---|---|---|---|---|---|---|
| *Lmb_jn3_03235* | (Gervais et al., 2017) | Cluster 4 | Border | YES | DTM_Ingwe (800bp) | H3K4me2 | YES | YES | <=1 TM | 134 | 2 | YES | Hypothetical protein J1614_004971 [*Plenodomus biglobosus*] Hypothetical protein T440DRAFT_471212 [*Plenodomus tracheiphilus* IPT5] Hypothetical protein IQ07DRAFT_405142 [*Pyrenochaeta* sp. DS3sAY3a] Hypothetical protein N0V94_006984 [*Neodidymelliopsis* sp. IMI 364377] Hypothetical protein NX059_001857 [*Plenodomus lindquistii*] | Good | None |
| *Lmb_jn3_03238* | - | Cluster 2 | AT | YES | Lmac_Grouper_595_20 (Overlapping) LmTelo1 (Overlapping) Lmac_Recon_92_20 (10bp) Lmac_Grouper_69_20 (250bp) Lmac_Grouper_595_20 (300bp) RLG_Olly (300bp) DTM_Sahana (600bp) LmTelo1 (900bp) LmTelo1 (1200bp) DTM_Ingwe (1500bp) RLG_Olly (2400bp) RLG_Polly (2600bp) RLG_Rolly (3700bp) Lmac_Grouper_595_20 (4000bp) | H3K9me3 | YES | YES | <=1 TM | 95 | 8 | NO | | Low | |
| *Lmb_jn3_03262* | *AvrLm4-7* (Parlange et al., 2009) | Cluster 2 | AT | YES | DTT_Finwe_2 (Overlapping) | H3K9me3 | YES | YES | <=1 TM | 144 | 8 | YES | Hypothetical protein J1614_000922 | Medium | |

| | | | | | | | | | | | | | | | |
|---|---|---|---|---|---|---|---|---|---|---|---|---|---|---|---|
| | | | | | RLG_Rolly (Overlapping)<br>DTT_Finwe_2 (10bp)<br>DTM_Ingwe (60bp)<br>Lmac_Grouper_2395_3 (450bp)<br>Lmac_Grouper_2395_3 (550bp)<br>RLG_Olly (700bp)<br>DTx_Olwe (1300bp)<br>DTM_Ingwe (1400bp)<br>Lmac_Grouper_2395_3 (2100bp)<br>DTM_Ingwe (2600bp)<br>LmTelo1 (2900bp)<br>RLx_Jolly (2900bp)<br>LmTelo2 (3500bp) | | | | | | | | [*Plenodomus biglobosus*]<br>Hypothetical protein BS50DRAFT_500295 [*Corynespora cassiicola Philippines*]<br>Hypothetical protein J1614_005249 [*Plenodomus biglobosus*] | | |
| *Lmb_jn3_03263* | - | Cluster 2 | Border | YES | RLG_Olly (500bp)<br>LmTelo1 (500bp)<br>RLG_Polly (1000bp)<br>RLG_Olly (1500bp)<br>Lmac_Recon_56_3 (2100bp)<br>DTM_Ingwe (2400bp)<br>Lmac_Recon_92_20 (2450bp)<br>LmTelo1 (2900bp)<br>LmTelo1 (3600bp) | H3K9me3 | YES | YES | <=1 TM | 143 | 7 | YES | Hypothetical protein J1614_000922 [*Plenodomus biglobosus*]<br>Hypothetical protein PtrM4_116010 [*Pyrenophora tritici-repentis*]<br>Hypothetical protein MPH_11524 [*Macrophomina phaseolina* MS6]<br>Hypothetical protein BS50DRAFT_500295 [*Corynespora cassiicola Philippines*]<br>Hypothetical protein MPH_00189 [*Macrophomina phaseolina* MS6] | Medium | |
| *Lmb_jn3_03397* | - | Cluster 2 | GC | NO | | H3K27me3 | YES | YES | <=1 TM | 144 | 5 | YES | Hypothetical protein NX059_002940 [*Plenodomus lindquistii*] | Good | Carbohydrate binding protein Clostrillin Spherulin 3A |

| | | | | | | | | | | | | | | | |
|---|---|---|---|---|---|---|---|---|---|---|---|---|---|---|---|
| | | | | | | | | | | | | | Hypothetical protein J1614_006546 [*Plenodomus biglobosus*] Hypothetical protein T440DRAFT_469237 [*Plenodomus tracheiphilus IPT5*] Hypothetical protein N0V83_009244 [*Neocucurbitaria cava*] Hypothetical protein BDW02DRAFT_567010 [*Decorospora gaudefroyi*] | | Beta and Gamma Crystallin Development specific protein S |
| *Lmb_jn3_03409* | - | Cluster 1 | GC | YES | RLG_Olly (100bp) | H3K27me3 | YES | YES | <=1 TM | 74 | 8 | NO | | Very Low | |
| *Lmb_jn3_03501* | - | Cluster 2 | Border | YES | DTx_Gimli (1800bp) DTM_Sahana (600bp) | H3K27me3 | YES | YES | <=1 TM | 129 | 7 | YES | Hypothetical protein J1614_006646 [*Plenodomus biglobosus*] Uncharacterized protein BDP81DRAFT_474374 [*Colletotrichum phormii*] Hypothetical protein PTT_01983 [*Pyrenophora teres* f. *teres* 0-1] LysM domain containing protein [*Pyrenophora teres* f. *maculata*] Hypothetical protein BHE90_005106 [*Fusarium euwallaceae*] | Medium | |
| *Lmb_jn3_03604* | - | Cluster 4 | GC | YES | DTx_Gimli (3900bp) | H3K27me3 | NO | YES | <=1 TM | 86 | 0 | YES | Hypothetical protein NX059_002709 [*Plenodomus lindquistii*] Hypothetical protein J1614_006760 | Medium | |

| | | | | | | | | | | | | | | | |
|---|---|---|---|---|---|---|---|---|---|---|---|---|---|---|---|
| | | | | | | | | | | | | | [*Plenodomus biglobosus*] Uncharacterized protein J4E85_009494 [*Alternaria conjuncta*] Hypothetical protein J4E91_010552 [*Alternaria rosae*] uncharacterized protein J4E93_003425 [*Alternaria ventricosa*] | | |
| *Lmb_jn3_03636* | - | Cluster 3 | GC | NO | | H3K27me3 | YES | YES | <=1 TM | 128 | 2 | NO | | Low | |
| *Lmb_jn3_03815* | *AvrLmS* paralogue | Cluster 2 | AT | YES | RLG_Polly (250bp) DTM_Ingwe (400bp) RLG_Polly (450bp) RLx_Jolly (500bp) RLG_Olly (600bp) RLx_Jolly (600bp) RLx_Jolly (800bp) DTT_Finwe_2 (900bp) RLx_Jolly (1300bp) RLC_Pholy (1800bp) RLG_Olly (3700bp) | H3K9me3 | YES | YES | <=1 TM | 141 | 8 | NO | | Low | |
| *Lmb_jn3_03821* | - | Cluster 5 | GC | NO | | H3K27me3 | YES | YES | <=1 TM | 247 | 10 | YES | Pectate lyase-domain-containing protein [*Alternaria rosae*] Hypothetical protein J4E91_009193 [*Alternaria rosae*] Hypothetical protein AA0111_g11952 [*Alternaria arborescens*] Uncharacterized protein J4E90_004559 [*Alternaria incomplexa*] Uncharacterized protein J4E82_006039 [*Alternaria postmessia*] | Good | Pectate Lyase Endo-Pectate Lyase Alginate Lyase LH3 Hexon-Interlacing Capsid Protein |

| | | | | | | | | | | | | | | |
|---|---|---|---|---|---|---|---|---|---|---|---|---|---|---|
| *Lmb_jn3_03822* | - | Cluster 5 | GC | NO | | H3K27me3 | NO | YES | <=1 TM | 107 | 2 | NO | | Low | |
| *Lmb_jn3_04095* | - | Cluster 4 | Border | YES | LmTelo2 (150bp) LmTelo1 (300bp) RLx_Ayoly (500bp) LmTelo1 (1000bp) RLG_Olly (1000bp) RLG_Dolly (1300bp) Lmac_Recon_83_20 (1800bp) RLG_Brawly (2300bp) RLG_Rolly (2500bp) | H3K9me3 | YES | YES | <=1 TM | 125 | 7 | YES | Uncharacterized protein PSV08DRAFT_232564 [*Bipolaris maydis*] Hypothetical protein FOXG_22493 [*Fusarium oxysporum* f. sp. *lycopersici* 4287] Hypothetical protein K456DRAFT_1723731 [*Colletotrichum gloeosporioides* 23] Secreted in xylem 5 [*Colletotrichum asianum*] Uncharacterized protein GCG54_00012396 [*Colletotrichum gloeosporioides*] | Good | KP6 Killer Toxin Subunit Alpha<br><br>Protein (Toxin) |
| *Lmb_jn3_04298* | - | Cluster 2 | AT | YES | DTx_Gimli (Overlapping) Lmac_Grouper_284_9 (Overlapping) DTT_Finwe_2 (150bp) RLC_Pholy (400bp) DTx_Gimli (450bp) Lmac_Grouper_2395_3 (700bp) RLG_Dolly (1200bp) RLC_Pholy (2000bp) Lmac_Grouper_1517_5 (2000bp) RLG_Olly (2000bp) RLG_Olly (2500bp) | H3K9me3 + H3K27me3 | YES | YES | <=1 TM | 93 | 10 | NO | Hypothetical protein T440DRAFT_436681 [*Plenodomus tracheiphilus IPT5*] Hypothetical protein BDZ85DRAFT_261523 [*Elsinoe ampelina*] Hypothetical protein AC578_6764 [*Pseudocercospora eumusae*] Teneurin-m isoform X3 [*Solenopsis invicta*] Teneurin-m isoform X2 [*Solenopsis invicta*] | Low | |
| *Lmb_jn3_04378* | - | Cluster 2 | GC | NO | | H3K4me2 | YES | YES | <=1 TM | 196 | 12 | NO | Hypothetical protein T440DRAFT_513574 [*Plenodomus* | Low | |

| | | | | | | | | | | | | | | | |
|---|---|---|---|---|---|---|---|---|---|---|---|---|---|---|---|
| | | | | | | | | | | | | | *tracheiphilus* IPT5] Hypothetical protein J1614_002618 [*Plenodomus biglobosus*] Hypothetical protein NX059_003971 [*Plenodomus lindquistii*] | | |
| *Lmb_jn3_04385* | This study | Cluster 5 | GC | YES | RLG_Olly (400bp) | H3K27me3 | YES | YES | <=1 TM | 90 | 6 | NO | | Medium | |
| *Lmb_jn3_04778* | *LmSTEE30* (Jiquel et al., 2021) | Cluster 5 | GC | NO | | H3K4me2 + H3K27me3 | YES | YES | <=1 TM | 108 | 10 | YES | Hypothetical protein T440DRAFT_397206 [*Plenodomus tracheiphilus IPT5*] Hypothetical protein SVAN01_01534 [*Stagonosporopsis vannaccii*] Hypothetical protein B5807_04569 [*Epicoccum nigrum*] Uncharacterized protein H3K4ME260DRAFT_297136 [*Cucurbitaria berberidis* CBS 394.84] CFEM13 protein [*Neostagonosporella sichuanensis*] | Good | None |
| *Lmb_jn3_05067* | - | Cluster 2 | AT | YES | DTx_Gimli (Overlapping) RLG_Brawly (Overlapping) Lmac_Grouper_284_9 (400bp) RLG_Polly (1000bp) RLC_Pholy (1400bp) | H3K9me3 | YES | YES | <=1 TM | 83 | 8 | NO | | Low | |
| *Lmb_jn3_05259* | - | Cluster 5 | GC | NO | | H3K27me3 | YES | YES | <=1 TM | 237 | 14 | YES | Polysaccharide lyase family 3 protein [*Plenodomus tracheiphilus* IPT5] Pectate lyase-like protein | Good | Pectate Lyase Endo-Pectate Lyase Particle-Associated Glycoside Hydrolase |

| | | | | | | | | | | | | | | | |
|---|---|---|---|---|---|---|---|---|---|---|---|---|---|---|---|
| | | | | | | | | | | | | | [*Pyrenochaeta* sp. DS3sAY3a] Hypothetical protein NX059_011979 [*Plenodomus lindquistii*] Uncharacterized protein J4E83_003839 [*Alternaria metachromatica*] Pectate lyase-like protein [*Alternaria rosae*] | | LH3 Hexon-Interlacing Capsid Protein Alpha-1,3-Glucanase |
| *Lmb_jn3_05316* | - | Cluster 2 | AT | YES | RLC_Zolly-1 (100bp) RLC_Zolly-1 (150bp) RLx_Jolly (500bp) Lmac_Grouper_595_20 (700bp) DTM_Sahana (750bp) RLG_Olly (1300bp) | H3K9me3 | YES | YES | <=1 TM | 90 | 8 | NO | | Low | |
| *Lmb_jn3_05329* | - | Cluster 1 | GC | NO | | H3K27me3 | YES | YES | <=1 TM | 82 | 4 | YES | Rapid alkalinization factor [*Plenodomus tracheiphilus* IPT5] Rapid alkalinization factor-domain-containing protein [*Elsinoe ampelina*] Rapid ALkalinization factor [*Fusarium globosum*] Hypothetical protein CT0861_00033 [*Colletotrichum tofieldiae*] ATPase [*Fusarium mundagurra*] | Low | |
| *Lmb_jn3_05445* | - | Cluster 3 | GC | NO | | H3K27me3 | YES | YES | <=1 TM | 65 | 6 | YES | Uncharacterized protein BKA58DRAFT_6160 [*Alternaria rosae*] Hypothetical protein AA0111_g5441 | Medium | |

| | | | | | | | | | | | | | | | |
|---|---|---|---|---|---|---|---|---|---|---|---|---|---|---|---|
| | | | | | | | | | | | | | [*Alternaria arborescens*] Hypothetical protein AG0111_0g869 [*Alternaria gaisen*] Hypothetical protein AA0116_g693 [*Alternaria tenuissima*] Hypothetical protein EK21DRAFT_80500 [*Setomelanomma holmii*] | | |
| *Lmb_jn3_05451* | - | Cluster 3 | GC | NO | | H3K27me3 | YES | YES | <=1 TM | 80 | 6 | NO | | Low | |
| *Lmb_jn3_05462* | - | Cluster 6 | GC | NO | | NO | YES | YES | <=1 TM | 126 | 4 | YES | Hypothetical protein J1614_000819 [*Plenodomus biglobosus*] Hypothetical protein T440DRAFT_393426 [*Plenodomus tracheiphilus IPT5*] Hypothetical protein NX059_009931 [*Plenodomus lindquistii*] Hypothetical protein TW65_03933 [*Stemphylium lycopersici*] periplasmic binding protein-like II [*Stemphylium lycopersici*] | Low | |
| *Lmb_jn3_05465* | (Jiquel et al., 2022) | Cluster 5 | GC | NO | | H3K27me3 | YES | YES | <=1 TM | 96 | 8 | YES | Hypothetical protein T440DRAFT_447411 [*Plenodomus tracheiphilus* IPT5] Hypothetical protein yc1106_00143 [*Curvularia clavata*] Hypothetical protein | Good | None |

| | | | | | | | | | | | | | | |
|---|---|---|---|---|---|---|---|---|---|---|---|---|---|---|
| | | | | | | | | | | | | | ACET3X_003158 [*Alternaria dauci*] Uncharacterized protein J4E86_000521 [*Alternaria arbusti*] Hypothetical protein TW65_07109 [*Stemphylium lycopersici*] | |
| *Lmb_jn3_05547* | *AvrLm14* (Degrave et al., 2021) | Cluster 2 | AT | YES | Lmac_Grouper_498_20 (Overlapping) RLC_Pholy (Overlapping) DTT_Finwe_2 (Overlapping) RLC_Zolly- (1000bp) DTM_Ingwe (1100bp) RLC_Pholy (1300bp) RLC_Pholy (2300bp) | H3K9me3 | YES | YES | <=1 TM | 135 | 5 | NO | Hypothetical protein J1614_006439 [*Plenodomus biglobosus*] | Low |
| *Lmb_jn3_05550* | - | Cluster 5 | Border | YES | RLG_Brawly (150bp) Lmac_Recon_56_3 (300bp) RLG_Olly (700bp) | H3K27me3 | NO | YES | <=1 TM | 67 | 0 | NO | | Medium |
| *Lmb_jn3_05637* | - | Cluster 4 | GC | NO | | H3K4me2 | YES | YES | <=1 TM | 57 | 0 | YES | Hypothetical protein ACET3X_001744 [*Alternaria dauci*] Carbohydrate esterase family 16 protein [*Bipolaris victoriae* FI3] Carbohydrate esterase family 16 protein [*Bipolaris zeicola* 26-R-13] Carbohydrate esterase [*Bipolaris maydis*] uncharacterized protein J4E86_006217 [*Alternaria arbusti*] | Low |
| *Lmb_jn3_05737* | (Gervais et al., 2017) | Cluster 4 | GC | NO | | H3K27me3 | YES | YES | <=1 TM | 95 | 8 | YES | Hypothetical protein EYC84_004918 [*Monilinia fructicola*] | Medium |

| | | | | | | | | | | | | | | |
|---|---|---|---|---|---|---|---|---|---|---|---|---|---|---|
| | | | | | | | | | | | | | Hypothetical protein EYC80_004424 [*Monilinia laxa*]<br>Hypothetical protein MFRU_059g00440 [*Monilinia fructicola*]<br>Hypothetical protein K4K59_003770 [*Colletotrichum* sp. SAR11_240]<br>Hypothetical protein Landi51_00424 [*Colletotrichum acutatum*] | |
| *Lmb_jn3_05738* | - | Cluster 2 | GC | NO | | H3K27me3 | YES | YES | <=1 TM | 129 | 8 | NO | | Low |
| *Lmb_jn3_05764* | - | Cluster 2 | GC | NO | | H3K27me3 | YES | YES | <=1 TM | 93 | 6 | NO | | Low |
| *Lmb_jn3_05765* | Present study | Cluster 4 | GC | NO | | H3K27me3 | YES | YES | <=1 TM | 84 | 6 | NO | | Low |
| *Lmb_jn3_05813* | - | Cluster 5 | Border | NO | | H3K27me3 | YES | YES | <=1 TM | 97 | 8 | YES | Uncharacterized protein CLAFUR5_08124 [*Fulvia fulva*]<br>Hypothetical protein HII31_01734 [*Pseudocercospora fuligena*]<br>Hypothetical protein E6O75_ATG03705 [*Venturia nashicola*]<br>Hypothetical protein E2P81_ATG03629 [*Venturia nashicola*]<br>Hypothetical protein TW65_00709 [*Stemphylium lycopersici*] | Medium |
| *Lmb_jn3_05821* | - | Cluster 2 | AT | YES | LmTelo1 (200bp)<br>DTM_Ingwe (300bp)<br>RLG_Polly (700bp)<br>DTx_Gimli (1700bp)<br>RLx_Ayoly (2700bp)<br>Lmac_Grouper_498_20 (3200bp) | H3K9me3 | NO | YES | <=1 TM | 91 | 4 | NO | | Low |

| | | | | | | | | | | | | | | | |
|---|---|---|---|---|---|---|---|---|---|---|---|---|---|---|---|
| | | | | | RLG_Olly (3400bp)<br>LmTelo1 (3500bp)<br>LmTelo1 (3500bp)<br>LmTelo2 (3600bp)<br>Lmac_Recon_56_3 (4000bp) | | | | | | | | | | |
| *Lmb_jn3_05822* | - | Cluster 2 | AT | YES | RLG_Olly (Overlapping)<br>RLx_Jolly (100bp)<br>DTM_Ingwe (300bp)<br>Lmac_Grouper_595_20 (700bp)<br>DTM_Ingwe (1100bp)<br>RLx_Ayoly (1500bp)<br>DTM_Sahana (1500bp)<br>DTF_Elwe (1900bp)<br>RLx_Ayoly (2100bp)<br>Lmac_Recon_83_20 (2100bp)<br>LmTelo1 (2500bp)<br>RLG_Polly (2700bp)<br>DTT_Finwe-3 (2900bp)<br>RLG_Rolly (3400bp)<br>DTT_Finwe-3 (3600bp) | H3K9me3 + H3K27me3 | YES | YES | <=1 TM | 193 | 5 | YES | Hypothetical protein EJ02DRAFT_408185 [*Clathrospora elynae*]<br>Hypothetical protein N0V95_004330 [*Ascochyta clinopodiicola*]<br>Hypothetical protein TW65_97118 [*Stemphylium lycopersici*]<br>Hypothetical protein NX059_005208 [*Plenodomus lindquistii*]<br>Hypothetical protein J1614_007190 [*Plenodomus biglobosus*] | Good | Putative uncharacterized protein |
| *Lmb_jn3_05985* | - | Cluster 2 | AT | YES | DTM_Lenwe (Overlapping)<br>Lmac_Grouper_284_9 (Overlapping)<br>RLx_Ayoly (800bp)<br>RLG_Olly (1500bp)<br>RLG_Brawly (1800bp)<br>RLG_Polly (1900bp)<br>DTF_Elwe (1900bp)<br>RLG_Olly (3000bp)<br>RLG_Olly (3400bp) | H3K9me3 + H3K27me3 | YES | YES | <=1 TM | 72 | 8 | NO | | Low | |

| | | | | | | | | | | | | | | | |
|---|---|---|---|---|---|---|---|---|---|---|---|---|---|---|---|
| | | | | | Lmac_Recon_92_20 (3900bp) Lmac_Grouper_595_20 (3900bp) | | | | | | | | | | |
| *Lmb_jn3_05986* | - | Cluster 2 | AT | YES | RLG_Brawly (Overlapping) RLG_Polly (Overlapping) DTF_Elwe (Overlapping) DTM_Lenwe (2600bp) Lmac_Grouper_284_9 (2700bp) LmTelo1 (2800bp) RLx_Ayoly (2900bp) RLG_Olly (3700bp) LmTelo2 (4000bp) | H3K9me3 + H3K27me3 | YES | YES | <=1 TM | 58 | 5 | NO | | Low | |
| *Lmb_jn3_06293* | - | Cluster 3 | Border | YES | Lmac_Grouper_284_9 (800bp) | H3K27me3 | YES | YES | <=1 TM | 126 | 8 | NO | | Medium | |
| *Lmb_jn3_06826* | (Jiquel et al., 2022) | Cluster 5 | GC | YES | DTM_Sahana (600bp) | H3K27me3 | YES | YES | <=1 TM | 87 | 9 | NO | | Low | |
| *Lmb_jn3_06962* | - | Cluster 2 | Border | YES | Lmac_Grouper_2395_3 (Overlapping)<br>Lmac_Grouper_69_20 (150bp) | H3K27me3 | YES | YES | <=1 TM | 90 | 7 | NO | | Low | |
| *Lmb_jn3_07074* | - | Cluster 5 | Border | YES | RLx_Ayoly (Overlapping) RLG_Olly (70bp) DTx_Gimli (300bp) RLC_Pholy (500bp) | H3K27me3 | YES | YES | <=1 TM | 119 | 8 | YES | Hypothetical protein F66182_3055 [*Fusarium sp.* NRRL 66182] Hypothetical protein FOQG_18271 [*Fusarium oxysporum* f. sp. *raphani* 54005] Hypothetical protein Forpe1208_v010791 [*Fusarium oxysporum* f. sp. *rapae*] Hypothetical protein FOTG_18590 [*Fusarium oxysporum* f. sp. *vasinfectum* 25433] Hypothetical protein BKA61DRAFT_583267 | Good | KP6 Killer Toxin Subunit Alpha Protein (Toxin) Uncharacterized Protein |

| | | | | | | | | | | | | | | | |
|---|---|---|---|---|---|---|---|---|---|---|---|---|---|---|---|
| | | | | | | | | | | | | | [*Leptodontidium* sp. MPI-SDFR-AT-0119] | | |
| *Lmb_jn3_07083* | - | Cluster 3 | GC | NO | | H3K4me2 + H3K27me3 | YES | YES | <=1 TM | 104 | 8 | YES | Uncharacterized protein J4E93_008803 [*Alternaria ventricosa*] Hypothetical protein IQ06DRAFT_347244 [*Phaeosphaeriaceae* sp. SRC1lsM3a] Uncharacterized protein J4E83_004896 [*Alternaria metachromatica*] Uncharacterized protein J4E92_004112 [*Alternaria infectoria*] uncharacterized protein J4E88_008151 [*Alternaria novae-zelandiae*] | Medium | |
| *Lmb_jn3_07098* | - | Cluster 5 | GC | YES | RLC_Pholy (200bp) | H3K27me3 | NO | YES | <=1 TM | 254 | 6 | YES | Lytic polysaccharide monooxygenase [*Plenodomus tracheiphilus* IPT5] Glycoside hydrolase family 61 protein [*Alternaria burnsii*] Hypothetical protein J4E91_008173 [*Alternaria rosae*] Uncharacterized protein ALTATR162_LOCUS11399 [*Alternaria atra*] Hypothetical protein N0V83_001999 [*Neocucurbitaria cava*] | Good | Polysaccharide Monooxygenase -3 Endoglucanase Putative Lytic Polysaccharide Monooxygenase GH61 Isozyme A Glycoside Hydrolase Family 61 |
| *Lmb_jn3_07106* | - | Cluster 4 | GC | YES | DTx_Valwe (4000bp) | H3K27me3 | YES | YES | <=1 TM | 253 | 0 | YES | Hypothetical protein J1614_011574 | Good | Rhamnogalacturonan Acetylesterase |

| | | | | | | | | | | | | | | | |
|---|---|---|---|---|---|---|---|---|---|---|---|---|---|---|---|
| | | | | | | | | | | | | | [*Plenodomus biglobosus*] Carbohydrate esterase family 12 protein [*Plenodomus tracheiphilus* IPT5] Hypothetical protein N0V83_001976 [*Neocucurbitaria cava*] Carbohydrate esterase family 12 protein [*Cucurbitaria berberidis* CBS 394.84] SGNH hydrolase-type esterase domain-containing protein [*Alternaria rosae*] | | Lipase/Acylhydrolase Lipase/Acylhydrolase Acetyl Xylan Esterase G-D-S-L Family Lipolytic Protein Putative Lipase from the G-D-S-L Family |
| *Lmb_jn3_07154* | - | Cluster 2 | GC | NO | | H3K4me2 | YES | YES | <=1 TM | 83 | 6 | NO | | Medium | |
| *Lmb_jn3_07285* | - | Cluster 2 | GC | YES | DTM_Ingwe (150bp) | H3K4me2 + H3K27me3 | YES | YES | <=1 TM | 256 | 4 | YES | RmlC-like cupin [*Plenodomus tracheiphilus* IPT5] Hypothetical protein J1614_010566 [*Plenodomus biglobosus*] RmlC-like cupin domain-containing protein [*Phaeosphaeriaceae* sp. PMI808] Hypothetical protein NX059_004808 [*Plenodomus lindquistii*] RmlC-like cupin [*Lizonia empirigonia*] | Good | Oxalate Oxidase 1 Peruvianin-I Canavalin 7S Vicilin Novel Protein with Potential Cupin Domain |
| *Lmb_jn3_07353* | - | Cluster 4 | Border | YES | RLC_Pholy (Overlapping) Lmac_Recon_56_3 (300bp) Lmac_Grouper_498_20 (400bp) DTx_Gimli (700bp) | H3K9me3 + H3K27me3 | YES | YES | <=1 TM | 111 | 10 | NO | | Low | |

| | | | | | | | | | | | | | | | |
|---|---|---|---|---|---|---|---|---|---|---|---|---|---|---|---|
| | | | | | DTx_Gimli (800bp)<br>RLx_Jolly (1300bp)<br>RLG_Olly (1600bp) | | | | | | | | | | |
| *Lmb_jn3_07355* | - | Cluster 1 | Border | YES | Lmac_Grouper_2395_3 (Overlapping)<br>RLG_Rolly (Overlapping)<br>LmTelo2 (200bp)<br>RLC_Zolly-1 (400bp)<br>DTF_Elwe (1000bp)<br>DTM_Ingwe (1200bp)<br>LmTelo2 (1300bp)<br>DTT_Finwe_2 (1600bp)<br>LmTelo2 (2200bp)<br>RLG_Rolly (2400bp)<br>RLG_Rolly (3000bp) | H3K9me3 + H3K27me3 | YES | YES | <=1 TM | 131 | 10 | YES | Hypothetical protein V5O48_011989 [*Marasmius crinis-equi*]<br>Hypothetical protein N7453_004843 [*Penicillium expansum*]<br>Hypothetical protein N0V82_005345 [*Gnomoniopsis sp. IMI 355080*]<br>Hypothetical protein V5O48_010522 [*Marasmius crinis-equi*]<br>Hypothetical protein DL98DRAFT_424568 [*Cadophora sp. DSE1049*] | Good | KP6 Killer Toxin Subunit Alpha Protein (Toxin)<br>Uncharacterized Protein |
| *Lmb_jn3_07393* | - | Cluster 1 | GC | NO | | H3K27me3 | NO | YES | <=1 TM | 90 | 3 | NO | | Low | |
| *Lmb_jn3_07512* | - | Cluster 2 | GC | YES | DTT_Finwe_2 (Overlapping)<br>RLx_Jolly (300bp) | H3K9me3 | YES | YES | <=1 TM | 92 | 8 | NO | | Very low | |
| *Lmb_jn3_07669* | - | Cluster 5 | GC | YES | DTx_Valwe (3400bp) | NA | NO | YES | <=1 TM | 82 | 1 | NO | | Medium | |
| *Lmb_jn3_07862* | *AvrLm6* (Fudal et al., 2007) | Cluster 2 | AT | YES | DTx_Gimli (Overlapping)<br>DTM_Ingwe (800bp)<br>RLG_Olly (1300bp)<br>RLG_Olly (1600bp)<br>RLG_Polly (2400bp) | H3K9me3 | YES | YES | <=1 TM | 145 | 6 | YES | Uncharacterized protein CTRU02_210410 [*Colletotrichum truncatum*]<br>Hypothetical protein ONS96_001429 [*Cadophora gregata* f. sp. *sojae*]<br>Uncharacterized protein GCG54_00006287 [*Colletotrichum gloeosporioides*]<br>m6 protein [*Colletotrichum kahawae*]<br>Hypothetical protein CSAL01_12831 | Good | KP6 Killer Toxin Subunit Alpha Protein (Toxin) |

| | | | | | | | | | | | | | | | |
|---|---|---|---|---|---|---|---|---|---|---|---|---|---|---|---|
| | | | | | | | | | | | | | [*Colletotrichum salicis*] | | |
| *Lmb_jn3_07863* | *AvrLm2* (Ghanbarnia et al., 2015) | Cluster 2 | AT | YES | DTM_Ingwe (100bp)<br>DTM_Ingwe (100bp)<br>LmTelo2 (150bp)<br>RLG_Olly (800bp)<br>RLG_Olly (900bp)<br>RLG_Olly (1000bp)<br>Lmac_Grouper_498_20 (2300bp)<br>RLG_Olly (2700bp) | H3K9me3 | YES | YES | <=1 TM | 233 | 8 | YES | Hypothetical protein CMEL01_08476 [*Colletotrichum melonis*]<br>Hypothetical protein CCUS01_04808 [*Colletotrichum cuscutae*]<br>hypothetical protein BDP67DRAFT_481437 [*Colletotrichum lupini*]<br>uncharacterized protein CLUP02_16779 [*Colletotrichum lupini*]<br>uncharacterized protein CORC01_14225 [*Colletotrichum orchidophilum*] | Low | |
| *Lmb_jn3_07874* | *AvrLm10_B* (Petit-Houdenot et al., 2019) | Cluster 2 | AT | YES | Lmac_Grouper_1227_7 (Overlapping)<br>RLG_Olly (50bp)<br>DTM_Ingwe (200bp)<br>RLG_Rolly (250bp)<br>DTx_Gimli (800bp)<br>Lmac_Recon_56_3 (1500bp)<br>DTT_Finwe-3 (2400bp)<br>RLG_Olly (2800bp)<br>DTT_Finwe-3 (2900bp)<br>DTM_Ingwe (3600bp)<br>DTT_Finwe-3 (3700bp) | H3K9me3 | YES | YES | <=1 TM | 191 | 2 | YES | Uncharacterized protein BDP81DRAFT_444630 [*Colletotrichum phormii*]<br>Histidine permease [*Colletotrichum scovillei*]<br>Histidine permease [*Colletotrichum acutatum*]<br>Histidine permease [*Colletotrichum salicis*]<br>Histidine permease [*Colletotrichum costaricense*] | Medium | |
| *Lmb_jn3_07875* | *AvrLm10_A* (Petit-Houdenot et al., 2019) | Cluster 2 | AT | YES | Lmac_Grouper_1227_7 (Overlapping)<br>Lmac_Grouper_69_20 (Overlapping) | H3K9me3 + H3K27me3 | YES | YES | <=1 TM | 121 | 7 | YES | Hypothetical protein CSAL01_09201 [*Colletotrichum salicis*] | Good | KP6 Killer Toxin Subunit Alpha Protein (Toxin) |

| | | | | | | | | | | | | | | | |
|---|---|---|---|---|---|---|---|---|---|---|---|---|---|---|---|
| | | | | | Lmac_Grouper_595_20 (150bp)<br>DTM_Lenwe (200bp)<br>RLG_Olly (200bp)<br>DTT_Finwe-2 (1900bp)<br>RLG_Polly (2200bp)<br>DTx_Valwe (2300bp)<br>DTT_Finwe-3 (3500bp)<br>RLG_Olly (3700bp)<br>DTT_Finwe-3 (4000bp) | | | | | | | | Hypothetical protein BDP67DRAFT_470684 [*Colletotrichum lupini*]<br>Hypothetical protein CSPX01_05366 [*Colletotrichum filicis*]<br>Uncharacterized protein CCOS01_01482 [*Colletotrichum costaricense*]<br>Hypothetical protein CSIM01_12189 [*Colletotrichum simmondsii*] | | |
| *Lmb_jn3_07895* | (Gervais et al., 2017) | Cluster 3 | GC | NO | | H3K4me2 | YES | YES | <=1 TM | 162 | 5 | YES | Hypothetical protein T440DRAFT_407891 [*Plenodomus tracheiphilus* IPT5]<br>Alt a 1 major allergen [*Pyrenochaeta* sp. MPI-SDFR-AT-0127]<br>Major allergen Alt a 1 [*Plenodomus lindquistii*]<br>Alternaria Alternata allergen *Alt A 1* [*Setomelanomma holmii*]<br>Hypothetical protein CC86DRAFT_298700 [*Ophiobolus disseminans*] | Good | Major Allergen ALT A 1<br>Effector Protein PEVD1<br>Root Induced Effector Protein TSP1<br>Trichoderma Secreted Protein TSP1 |
| *Lmb_jn3_07916* | - | Cluster 2 | AT | YES | RLG_Olly (Overlapping)<br>RLG_Olly (Overlapping)<br>RLG_Polly (50bp)<br>DTT_Finwe-2 (150bp)<br>DTT_Finwe-2 (250bp)<br>RLG_Olly (500bp)<br>DTF_Elwe (1700bp) | H3K9me3 | YES | YES | <=1 TM | 99 | 10 | NO | | Low | |

| | | | | | | | | | | | | | | |
|---|---|---|---|---|---|---|---|---|---|---|---|---|---|---|
| | | | | | RLG_Olly (2000bp) RLG_Olly (2400bp) RLx_Jolly (2400bp) RLx_Jolly (2400bp) RLC_Pholy (2400bp) | | | | | | | | | |
| *Lmb_jn3_07919* | (Jiquel et al., 2022) | Cluster 5 | Border | NO | | H3K27me3 | YES | YES | <=1 TM | 176 | 0 | YES | Hypothetical protein J1614_000971 [*Plenodomus biglobosus*] Hypothetical protein AG0111_0g10036 [*Alternaria gaisen*] Hypothetical protein AA0111_g6108 [*Alternaria arborescens*] Uncharacterized protein GT037_011127 [*Alternaria burnsii*] Uncharacterized protein J4E82_003957 [*Alternaria postmessia*] | Good | Signal Recognition Particle Subunit SRP68 Human Gene for Small Cytoplasmic 7SL RNA (7L30.1) SRP RNA ER Lumen Protein-Retaining Receptor 2 ACL4 |
| *Lmb_jn3_07939* | - | Cluster 3 | GC | NO | | H3K4me2 + H3K27me3 | YES | YES | <=1 TM | 162 | 3 | YES | Hypothetical protein J1614_000991 [*Plenodomus biglobosus*] Hypothetical protein T440DRAFT_473202 [*Plenodomus tracheiphilus* IPT5] Hypothetical protein NX059_000427 [*Plenodomus lindquistii*] Hypothetical protein BKA66DRAFT_452532 [*Pyrenochaeta* sp. MPI-SDFR-AT-0127] Uncharacterized protein | Medium | |

| | | | | | | | | | | | | | | | |
|---|---|---|---|---|---|---|---|---|---|---|---|---|---|---|---|
| | | | | | | | | | | | | | K460DRAFT_367570 [*Cucurbitaria berberidis CBS* 394.84] | | |
| *Lmb_jn3_08094* | *LmSTEE35* (Jiquel et al., 2021) | Cluster 4 | Border | YES | DTF_Elwe (200bp)<br>Lmac_Grouper_2395_3 (400bp)<br>RLG_Olly (400bp)<br>B35_Dolly (1200bp)<br>RLG_Rolly (1500bp)<br>Lmac_Grouper_2395_3 (1700bp)<br>Lmac_Recon_56_3 (2000bp)<br>RLG_Olly (2400bp) | H3K9me3 | YES | YES | <=1 TM | 124 | 7 | YES | Hypothetical protein CSAL01_09201 [*Colletotrichum salicis*]<br>Secreted in xylem 5 [*Colletotrichum asianum*]<br>Hypothetical protein K456DRAFT_1723731 [*Colletotrichum gloeosporioides* 23]<br>Uncharacterized protein CGCA056_v012554 [*Colletotrichum aenigma*]<br>Uncharacterized protein BDP55DRAFT_641301 [*Colletotrichum godetiae*] | Good | KP6 Killer Toxin Subunit Alpha Protein (Toxin) |
| *Lmb_jn3_08095* | - | Cluster 4 | Border | YES | RLG_Rolly (Overlapping)<br>Lmac_Grouper_2395_3 (50bp)<br>Lmac_Recon_56_3 (600bp)<br>DTF_Elwe (1800bp)<br>RLG_Olly (1950bp)<br>Lmac_Grouper_2395_3 (2000bp)<br>B35_Dolly (2700bp)<br>RLG_Olly (2900bp) | H3K9me3 | YES | YES | <=1 TM | 172 | 2 | NO | | Very low | |
| *Lmb_jn3_08343* | *AvrLmS* (Neik et al., 2020a) | Cluster 2 | AT | YES | RLG_Rolly (Overlapping)<br>Lmac_Grouper_2395_3 (Overlapping)<br>DTT_Finwe-2 (200bp)<br>Lmac_Grouper_1227_7 (500bp) | H3K9me3 | YES | YES | <=1 TM | 142 | 8 | NO | | Low | |

| | | | | | | | | | | | | | | | |
|---|---|---|---|---|---|---|---|---|---|---|---|---|---|---|---|
| | | | | | RLG_Polly (600bp)<br>RLG_Rolly (1200bp)<br>RLG_Polly (1200bp)<br>RLG_Polly (1600bp) | | | | | | | | | | |
| *Lmb_jn3_08356* | - | Cluster 4 | GC | YES | DTT_Finwe-3 (500bp)<br>RLG_Polly (600bp)<br>DTT_Finwe-3 (800bp)<br>RLC_Pholy (1500bp) | H3K27me3 | YES | YES | <=1 TM | 149 | 10 | YES | Uncharacterized protein N0V96_005051 [*Colletotrichum fioriniae*]<br>Hypothetical protein C8034_v003725 [*Colletotrichum sidae*]<br>Hypothetical protein Cob_v009472 [*Colletotrichum orbiculare* MAFF 240422]<br>Uncharacterized protein BDZ83DRAFT_646538 [*Colletotrichum acutatum*]<br>Hypothetical protein CGLO_17245 [*Colletotrichum gloeosporioides* Cg-14] | Very Low | |
| *Lmb_jn3_08417* | - | Cluster 2 | AT | YES | RLG_Olly (400bp)<br>Lmac_Grouper_1829_15 (700bp)<br>LmTelo1 (800bp)<br>RLx_Ayoly (900bp)<br>RLG_Polly (1400bp)<br>Lmac_Grouper_1227_4 (2200bp)<br>LmTelo1 (1800bp)<br>LmTelo2 (2000bp)<br>RLG_Olly (2300bp) | H3K9me3 + H3K27me3 | YES | YES | <=1 TM | 110 | 6 | NO | | Very Low | |
| *Lmb_jn3_08418* | - | Cluster 2 | AT | YES | RLG_Rolly (Overlapping)<br>LmTelo1 (300bp)<br>RLG_Olly (3600bp) | H3K9me3 + H3K27me3 | YES | YES | <=1 TM | 139 | 6 | YES | Predicted protein [*Pyrenophora tritici-repentis* Pt-1C-BFP]<br>Hypothetical protein | Good | Probable SECDF Protein-Export Membrane Protein Molybdenum cofactor |

| | | | | | | | | | | | | | | | |
|---|---|---|---|---|---|---|---|---|---|---|---|---|---|---|---|
| | | | | | RLG_Polly (3650bp) | | | | | | | | A1F94_006945 [*Pyrenophora tritici-repentis*]<br>Hypothetical protein PtrM4_116000 [*Pyrenophora tritici-repentis*]<br>Hypothetical protein PTT_10442 [*Pyrenophora teres* f. *teres* 0-1]<br>Hypothetical protein PTMSG1_10058 [*Pyrenophora teres* f. *maculata*] | | biosynthesis protein MOAC<br>Molybdenum cofactor biosynthesis protein C<br>Ribulose-1,5-Bisphosphate Carboxylase/Oxygenase L<br>Receptor-Type Tyrosine-Protein phosphatase-like N |
| *Lmb_jn3_08515* | - | Cluster 2 | GC | YES | DTx_Valwe (600bp) | H3K27me3 | YES | YES | <=1 TM | 103 | 8 | NO | | Low | |
| *Lmb_jn3_08794* | - | Cluster 2 | Border | YES | DTF_Elwe (10bp)<br>RLG_Polly (400bp)<br>RLx_Jolly (800bp)<br>RLG_Polly (3700bp) | H3K9me3 | YES | YES | <=1 TM | 124 | 8 | YES | Hypothetical protein IWW34DRAFT_811521 [*Fusarium oxysporum* f. sp. *albedinis*]<br>Hypothetical protein FOTG_18991 [*Fusarium oxysporum* f. sp. *vasinfectum 25433*]<br>Hypothetical protein H9L39_18419 [*Fusarium oxysporum* f. sp. *albedinis*]<br>Hypothetical protein FOQG_18878 [*Fusarium oxysporum* f. sp. *raphani* 54005]<br>Hypothetical protein FOQG_18767 [*Fusarium oxysporum* f. sp. *raphani* 54005] | Medium | |
| *Lmb_jn3_08992* | - | Cluster 1 | GC | NO | | H3K27me3 | YES | YES | <=1 TM | 250 | 4 | YES | Carbohydrate esterase family 5 protein [*Plenodomus tracheiphilus* IPT5] | Good | Cutinase 1<br>Carbohydrate Esterase Family 5<br>Cutinase-Like Protein |

| | | | | | | | | | | | | | | | |
|---|---|---|---|---|---|---|---|---|---|---|---|---|---|---|---|
| | | | | | | | | | | | | | Hypothetical protein NX059_008627 [*Plenodomus lindquistii*]<br>Hypothetical protein J1614_004881 [*Plenodomus biglobosus*]<br>Cutinase-domain-containing protein [*Pyrenochaeta* sp. DS3sAY3a]<br>Hypothetical protein N0V91_010055 [*Didymella pomorum*] | | Acetyl Xylan Esterase<br>Gene 12 protein |
| *Lmb_jn3_09025* | - | Cluster 2 | GC | NO | | H3K27me3 | YES | YES | <=1 TM | 231 | 10 | YES | Carbohydrate esterase family 5 protein [*Plenodomus tracheiphilus* IPT5]<br>Hypothetical protein NX059_008660 [*Plenodomus lindquistii*]<br>Hypothetical protein N0V94_003080 [*Neodidymelliopsis* sp. IMI 364377]<br>Acetylxylan esterase precursor [*Boeremia exigua*]<br>Carbohydrate esterase [*Nothophoma quercina*] | Good | Acetyl Xylan Esterase II<br>Cutinase-Like Enzyme<br>Conserved membrane protein of uncharacterized function<br>Cutinase<br>Cutinase 1 |
| *Lmb_jn3_09276* | - | Cluster 2 | GC | NO | | NO | YES | YES | <=1 TM | 42 | 3 | NO | | Low | |
| *Lmb_jn3_09383* | - | Cluster 4 | GC | YES | DTF_Elwe (2900bp)<br>DTx_Gimli (2900bp) | H3K27me3 | YES | YES | <=1 TM | 256 | 10 | YES | Polysaccharide lyase family 3 protein [*Exserohilum turcica* Et28A]<br>Polysaccharide lyase family 3 protein [*Stemphylium lycopersici*] | Good | Pectate Lyase<br>Endo-Pectate Lyase<br>Alpha-1,3-Glucanase<br>Preneck Appendage Protein<br>Endopolygalacturonase |

| | | | | | | | | | | | | | | | |
|---|---|---|---|---|---|---|---|---|---|---|---|---|---|---|---|
| | | | | | | | | | | | | | Pectate lyase [*Alternaria panax*] Polysaccharide lyase [*Bipolaris maydis*] Polysaccharide lyase family 3 protein [*Bipolaris sorokiniana* ND90Pr] | | |
| *Lmb_jn3_09385* | Present study | Cluster 4 | GC | YES | DTx_Gimli (200bp) | H3K27me3 | YES | YES | <=1 TM | 124 | 4 | NO | | Medium | |
| *Lmb_jn3_09539* | - | Cluster 4 | GC | YES | DTx_Valwe (2800bp) | H3K4me2 | YES | YES | <=1 TM | 264 | 2 | YES | Carbohydrate esterase family 12 protein [*Plenodomus tracheiphilus* IPT5] Hypothetical protein J1614_009210 [*Plenodomus biglobosus*] Hypothetical protein NX059_006772 [*Plenodomus lindquistii*] SGNH hydrolase-type esterase domain-containing protein [*Ampelomyces quisqualis*] Hypothetical protein H2201_008798 [*Coniosporium apollinis*] | Good | Rhamnogalacturonan Acetylesterase Lipase/Acylhydrolase Putative Lipase From The G-D-S-L Family G-D-S-L Family Lipolytic Protein LAE5 |
| *Lmb_jn3_09745* | - | Cluster 2 | Border | YES | RLx_Ayoly (Overlapping) RLG_Olly (300bp) LmTelo1 (1000bp) DTx_Valwe (1550bp) LmTelo1 (2700bp) LmTelo1 (2800bp) RLG_Polly (3200bp) RLx_Jolly (4000bp) | H3K9me3 + H3K27me3 | YES | YES | <=1 TM | 121 | 8 | YES | Hypothetical protein FOXG_22493 [*Fusarium oxysporum* f. sp. *lycopersici* 4287] Hypothetical protein FOTG_18790 [*Fusarium oxysporum* f. sp. *vasinfectum* 25433] uncharacterized protein CGCA056_v012554 | Good | KP6 Killer Toxin Subunit Alpha Protein (Toxin) |

| | | | | | | | | | | | | | | | |
|---|---|---|---|---|---|---|---|---|---|---|---|---|---|---|---|
| | | | | | | | | | | | | | [Colletotrichum aenigma] Secreted in xylem 5 [*Colletotrichum asianum*] Hypothetical protein K456DRAFT_1723731 [*Colletotrichum gloeosporioides* 23] | | |
| *Lmb_jn3_09746* | - | Cluster 2 | AT | YES | LmTelo1 (Overlapping) DTx_Valwe (Overlapping) LmTelo1 (1000bp) LmTelo1 (1100bp) RLx_Ayoly (1200bp) RLG_Olly (1300bp) RLG_Polly (1600bp) RLx_Jolly (2400bp) RLC_Pholy (3500bp) | H3K9me3 | YES | YES | <=1 TM | 167 | 3 | NO | | Low | |
| *Lmb_jn3_09838* | - | Cluster 6 | GC | NO | | H3K27me3 | NO | YES | <=1 TM | 88 | 3 | NO | | Low | |
| *Lmb_jn3_09935* | - | Cluster 4 | Border | NO | | H3K27me3 | YES | YES | <=1 TM | 256 | 11 | YES | Hypothetical protein ACET3X_002507 [*Alternaria dauci*] Polysaccharide lyase family 3 protein [*Bipolaris zeicola* 26-R-13] Pectate lyase precursor [*Alternaria rosae*] Uncharacterized protein J4E83_006960 [*Alternaria metachromatica*] Uncharacterized protein J4E85_002736 [*Alternaria conjuncta*] | Good | Pectate Lyase Endo-Pectate Lyase Short-Tailed Cyanophage Tailspike Receptor-Bindin LH3 Hexon-Interlacing Capsid Protein Alginate Lyase |
| *Lmb_jn3_09948* | - | Cluster 5 | GC | YES | DTF_Elwe (100bp) | NA | NO | YES | <=1 TM | 72 | 4 | NO | | Low | |
| *Lmb_jn3_09959* | - | Cluster 6 | GC | NO | | H3K27me3 | NO | YES | <=1 TM | 78 | 2 | NO | | Low | |

| | | | | | | | | | | | | | | | |
|---|---|---|---|---|---|---|---|---|---|---|---|---|---|---|---|
| *Lmb_jn3_10035* | - | Cluster 4 | GC | YES | DTx_Gimli (3100bp) | H3K4me2 | YES | YES | <=1 TM | 237 | 3 | YES | Necrosis-and ethylene-inducing protein-like protein 1 precursor [*Plenodomus tracheiphilus IPT5*] Hypothetical protein J1614_010756 [*Plenodomus biglobosus*] Necrosis-and ethylene-inducing protein-like protein 1 precursor [*Cucurbitaria berberidis* CBS 394.84] Hypothetical protein J4E80_006257 [*Alternaria* sp. BMP 0032] Uncharacterized protein J4E86_001461 [*Alternaria arbusti*] | Good | NEP1-Like Protein |
| *Lmb_jn3_10106* | *AvrLm5-9* (Ghanbarnia et al., 2018) | Cluster 2 | AT | YES | RLC_Pholy (Overlapping) RLC_Pholy (Overlapping) LmTelo1 (Overlapping) LmTelo1 (250bp) DTM_Ingwe (400bp) RLG_Olly (450bp) RLG_Polly (1000bp) RLG_Olly (1700bp) RLG_Olly (2700bp) Lmac_Grouper_498_20 (3900bp) RLG_Polly (4000bp) | H3K9me3 | YES | YES | <=1 TM | 142 | 7 | NO | | Low | |
| *Lmb_jn3_10523* | - | Cluster 4 | GC | NO | | NA | YES | YES | <=1 TM | 132 | 6 | YES | Uncharacterized protein CABS01_16583 [*Colletotrichum abscissum*] | Medium | |

| | | | | | | | | | | | | | | | |
|---|---|---|---|---|---|---|---|---|---|---|---|---|---|---|---|
| | | | | | | | | | | | | | Hypothetical protein BDP67DRAFT_16819 [*Colletotrichum lupini*]<br>Hypothetical protein H3K4ME256DRAFT_1917430 [*Colletotrichum gloeosporioides* 23]<br>Uncharacterized protein CLUP02_02900 [*Colletotrichum lupini*] | | |
| *Lmb_jn3_10604* | - | Cluster 5 | Border | NO | | H3K27me3 | NO | YES | <=1 TM | 90 | 9 | NO | | Low | |
| *Lmb_jn3_10706* | - | Cluster 2 | Border | YES | RLx_Ayoly (Overlapping)<br>DTM_Ingwe (50bp)<br>DTM_Ingwe (100bp) | NA | YES | YES | <=1 TM | 119 | 8 | NO | | Low | |
| *Lmb_jn3_10725* | - | Cluster 5 | GC | NO | | H3K27me3 | NO | YES | <=1 TM | 74 | 3 | NO | | Medium | |
| *Lmb_jn3_10844* | - | Cluster 4 | Border | YES | DTx_Gimli (800bp)<br>RLG_Olly (900bp)<br>RLG_Olly (1000bp)<br>RLG_Olly (1150bp) | H3K27me3 | YES | YES | <=1 TM | 233 | 0 | YES | Hypothetical protein J1614_002218 [*Plenodomus biglobosus*]<br>DUF1961-domain-containing protein [*Plenodomus tracheiphilus* IPT5]<br>Hypothetical protein NX059_003562 [*Plenodomus lindquistii*]<br>Hypothetical protein BKA63DRAFT_499199 [*Paraphoma chrysanthemicola*]<br>DUF1961-domain-containing protein [*Karstenula rhodostoma* CBS 690.94] | Good | Protein Yesu<br>Putative Glycosyl Hydrolase<br>Lectin-Like Protein<br>Protein Ergic-53<br>Putative Secreted Glucosylhydrolase |

| | | | | | | | | | | | | | | |
|---|---|---|---|---|---|---|---|---|---|---|---|---|---|---|
| *Lmb_jn3_10933* | (Jiquel et al., 2022) | Cluster 5 | GC | YES | DTx_Valwe (3300bp) | H3K4me2 | YES | YES | <=1 TM | 50 | 2 | YES | Hypothetical protein EK21DRAFT_77352 [*Setomelanomma holmii*] Hypothetical protein SNOG_12139 [*Parastagonospora nodorum* SN15] Hypothetical protein T440DRAFT_519232 [*Plenodomus tracheiphilus IPT5*] Hypothetical protein DE146DRAFT_757649 [*Phaeosphaeria* sp. MPI-PUGE-AT-0046c] Hypothetical protein M011DRAFT_480614 [*Sporormia fimetaria* CBS 119925] | Medium | |
| *Lmb_jn3_11031* | - | Cluster 4 | GC | YES | DTx_Gimli (2200bp) | H3K27me3 | YES | YES | <=1 TM | 178 | 10 | YES | Hypothetical protein J1614_006004 [*Plenodomus biglobosus*] Hypothetical protein T440DRAFT_419679 [*Plenodomus tracheiphilus IPT5*] Hypothetical protein HBH52_143540 [*Parastagonospora nodorum*] Hypothetical protein SNOG_08125 [*Parastagonospora nodorum* SN15] hHypothetical protein PTT_00797 [*Pyrenophora teres* f. *teres* 0-1] | Good | Ribonuclease MS Ribonuclease T1 Protein (Ribonuclease T1) Guanyl-Specific Ribonuclease T1 Ribonuclease F1 |
| *Lmb_jn3_11224* | - | Cluster 2 | Border | YES | DTx_Gimli (50bp) | H3K27me3 | YES | YES | <=1 TM | 96 | 10 | NO | | Low | |

| | | | | | | | | | | | | | | | |
|---|---|---|---|---|---|---|---|---|---|---|---|---|---|---|---|
| | | | | | RLx_Ayoly (1800bp) | | | | | | | | | | |
| *Lmb_jn3_11225* | - | Cluster 2 | Border | YES | RLG_Brawly (50bp)<br>RLx_Ayoly (50bp) | H3K27me3 | YES | YES | <=1 TM | 109 | 11 | NO | | Low | |
| *Lmb_jn3_11242* | - | Cluster 3 | GC | NO | | H3K4me2 | YES | YES | <=1 TM | 212 | 5 | YES | Hypothetical protein J1614_001990 [*Plenodomus biglobosus*]<br>Uncharacterized protein J4E82_005795 [*Alternaria postmessia*]<br>Hypothetical protein IG631_06712 [*Alternaria alternata*]<br>Hypothetical protein AA0111_g8720 [*Alternaria arborescens*]<br>Hypothetical protein DE146DRAFT_661025 [*Phaeosphaeria* sp. MPI-PUGE-AT-0046c] | Medium | |
| *Lmb_jn3_11364* | *LmSTEE98* (Jiquel et al., 2021) | Cluster 4 | GC | YES | DTx_Gimli (2200bp)<br>DTx_Gimli (2700bp) | H3K27me3 | YES | YES | <=1 TM | 56 | 6 | NO | | Very Low | |
| *Lmb_jn3_11375* | - | Cluster 4 | GC | NO | | H3K27me3 | YES | YES | <=1 TM | 93 | 6 | YES | Uncharacterized protein M421DRAFT_420020 [*Didymella exigua* CBS 183.55]<br>Uncharacterized protein COCSADRAFT_37678 [*Bipolaris sorokiniana* ND90Pr]<br>Hypothetical protein FDECE_10468 [*Fusarium decemcellulare*]<br>Uncharacterized protein PSV08DRAFT_261661 [*Bipolaris maydis*] | Good | None |

| | | | | | | | | | | | | | | |
|---|---|---|---|---|---|---|---|---|---|---|---|---|---|---|
| | | | | | | | | | | | | | Hypothetical protein DER45DRAFT_636414 [*Fusarium avenaceum*] | |
| *Lmb_jn3_11376* | - | Cluster 4 | GC | NO | | H3K27me3 (AI) | YES | YES | <=1 TM | 108 | 4 | YES | Uncharacterized protein PSV08DRAFT_261661 [*Bipolaris maydis*]<br>Hypothetical protein FBULB1_3794 [*Fusarium bulbicola*]<br>Uncharacterized protein COCSADRAFT_37678 [*Bipolaris sorokiniana* ND90Pr]<br>Hypothetical protein FVEG_16584 [*Fusarium verticillioides* 7600]<br>Uncharacterized protein FMAN_08286 [*Fusarium mangiferae*] | Low |
| *Lmb_jn3_11426* | - | Cluster 1 | GC | YES | RLG_Brawly (Overlapping) | H3K27me3 (AI) | YES | YES | <=1 TM | 79 | 8 | NO | | Low |
| *Lmb_jn3_11428* | - | Cluster 4 | GC | YES | RLG_Brawly (2200bp) | H3K27me3 (AI) | YES | YES | <=1 TM | 80 | 8 | NO | | Low |
| *Lmb_jn3_11454* | | Cluster 2 | AT | YES | RLG_Olly (50bp)<br>RLG_Olly (200bp)<br>DTM_Ingwe (500bp)<br>DTM_Sahana (1500bp)<br>Lmac_Grouper_498_20 (1600bp)<br>RLG_Olly (2000bp)<br>RLx_Ayoly (2200bp)<br>LmTelo2 (2800bp)<br>LmTelo2 (3800bp) | H3K9Me3 (AI) | YES | YES | <=1 TM | 248 | 8 | YES | Hypothetical protein SLS57_011064 [*Botryosphaeria dothidea*] | Very low |
| *Lmb_jn3_11455* | - | Cluster 2 | Border | YES | RLx_Ayoly (150bp)<br>Lmac_Recon_83_20 (700bp)<br>LmTelo1 (800bp) | H3K27me3 | YES | YES | <=1 TM | 81 | 8 | NO | | Low |

| | | | | | | | | | | | | | | | |
|---|---|---|---|---|---|---|---|---|---|---|---|---|---|---|---|
| | | | | | RLG_Olly (1000bp) Lmac_Recon_92_20 (1000bp) RLC_Pholy (1200bp) Lmac_Grouper_284_9 (4000bp) | | | | | | | | | | |
| *Lmb_jn3_11546* | - | Cluster 1 | GC | NO | | H3K4me2 + H3K27me3 | YES | YES | <=1 TM | 202 | 3 | YES | Hypothetical protein T440DRAFT_399805 [*Plenodomus tracheiphilus IPT5*] Hypothetical protein NX059_010146 [*Plenodomus lindquistii*] Hypothetical protein N0V83_003568 [*Neocucurbitaria cava*] Hypothetical protein IQ07DRAFT_497498 [*Pyrenochaeta* sp. DS3sAY3a] Hypothetical protein J1614_006970 [*Plenodomus biglobosus*] | Good | Superoxide Dismutase [Cu-Zn] Copper Chaperone for Super-Oxide Dismutase Extracellular Superoxide Dismutase |
| *Lmb_jn3_11916* | - | Cluster 4 | GC | NO | | H3K27me3 | YES | YES | <=1 TM | 102 | 6 | YES | Hypothetical protein NX059_000100 [*Plenodomus lindquistii*] Hypothetical protein J1614_008235 [*Plenodomus biglobosus*] Hypothetical protein T440DRAFT_458471 [*Plenodomus tracheiphilus* IPT5] Hypothetical protein TW65_07408 [*Stemphylium lycopersici*] Hypothetical protein BDU57DRAFT_546691 | Good | KP6 Killer Toxin Subunit Alpha Protein (Toxin) Uncharacterized Protein |

| | | | | | | | | | | | | | | | |
|---|---|---|---|---|---|---|---|---|---|---|---|---|---|---|---|
| | | | | | | | | | | | | | [*Ampelomyces quisqualis*] | | |
| *Lmb_jn3_11952* | - | Cluster 4 | GC | NO | | H3K27me3 | YES | YES | <=1 TM | 106 | 9 | NO | | Good | KP6 Killer Toxin Subunit Alpha Protein (Toxin) Uncharacterized Protein |
| *Lmb_jn3_11990* | - | Cluster 2 | AT | YES | DTM_Ingwe (Overlapping)<br>DTM_Ingwe 200bp)<br>Lmac_Grouper_595_20 (200bp)<br>Lmac_Grouper_2395_3 (300bp)<br>Lmac_Grouper_1829_15 (400bp)<br>DTM_Ingwe 800bp)<br>Lmac_Grouper_498_20 (900bp)<br>DTx_Gimli (1000bp)<br>Lmac_Grouper_2395_3 (1100bp)<br>Lmac_Grouper_1829_15 (1300bp)<br>DTx_Gimli (1400bp)<br>RLG_Polly (1600bp)<br>DTx_Gimli (1800bp)<br>Lmac_Grouper_595_20 (2000bp)<br>Lmac_Grouper_284_9 (2000bp)<br>RLG_Olly (2000bp)<br>Lmac_Grouper_498_20 (2800bp)<br>RLC_Pholy (3600bp) | H3K9me3 | YES | YES | <=1 TM | 127 | 8 | YES | | Low | |
| *Lmb_jn3_12121* | - | Cluster 4 | GC | YES | RLG_Brawly (Overlapping)<br>DTx_Olwe (300bp)<br>RLG_Dolly (600bp)<br>DTx_Gimli (3500bp) | H3K9me3 | YES | YES | <=1 TM | 106 | 4 | NO | | Very Low | |
| *Lmb_jn3_12124* | - | Cluster 2 | GC | YES | DTx_Gimli (100bp)<br>DTx_Gimli (1000bp) | NO | YES | YES | <=1 TM | 147 | 6 | YES | Uncharacterized protein CORC01_12764 [*Colletotrichum orchidophilum*] | Low | |

| | | | | | | | | | | | | | | |
|---|---|---|---|---|---|---|---|---|---|---|---|---|---|---|
| *Lmb_jn3_12242* | - | Cluster 2 | AT | YES | RLx_Ayoly (3200bp) | H3K4me2 + H3K9me3 + H3K27me3 | NO | YES | <=1 TM | 80 | 4 | YES | Hypothetical protein P153DRAFT_418347 [*Dothidotthia symphoricarpi* CBS 119687]<br>Hypothetical protein BU16DRAFT_473280 [*Lophium mytilinum*]<br>Hypothetical protein P280DRAFT_533107 [*Massarina eburnea* CBS 473.64]<br>Hypothetical protein DM02DRAFT_682029 [*Periconia macrospinosa*]<br>Hypothetical protein BO83DRAFT_231414 [*Aspergillus eucalypticola* CBS 122712] | Low |
| *Lmb_jn3_12249* | - | Cluster 6 | Border | YES | Lmac_Recon_56_3 (1300bp)<br>Lmac_Grouper_1227_4 (1400bp)<br>RLC_Pholy (1700bp) | H3K27me3 | YES | YES | <=1 TM | 56 | 6 | YES | Hypothetical protein J1614_001448 [*Plenodomus biglobosus*]<br>Hypothetical protein BKA63DRAFT_420914 [*Paraphoma chrysanthemicola*]<br>Hypothetical protein FB567DRAFT_598679 [*Paraphoma chrysanthemicola*]<br>Hypothetical protein IQ06DRAFT_351788 [*Stagonospora* sp. SRC1lsM3a]<br>Hypothetical protein BKA66DRAFT_519649 [*Pyrenochaeta* sp. MPI-SDFR-AT-0127] | Low |

| | | | | | | | | | | | | | | | |
|---|---|---|---|---|---|---|---|---|---|---|---|---|---|---|---|
| *Lmb_jn3_12320* | - | Cluster 4 | GC | NO | | H3K4me2 | YES | YES | <=1 TM | 210 | 2 | YES | Hypothetical protein J1614_003790 [*Plenodomus biglobosus*] Hypothetical protein NX059_011375 [*Plenodomus lindquistii*] PEBP-like protein [*Plenodomus tracheiphilus* IPT5] PEBP-like protein [*Pyrenochaeta* sp. DS3sAY3a] PEBP-like protein [*Cucurbitaria berberidis* CBS 394.84] | Good | Flowering Locus T Protein Carboxypeptidase Y Protein Flowering Locus T Terminal Flower 1 Protein Protein Heading Date 3A |
| *Lmb_jn3_12368* | - | Cluster 5 | GC | NO | | H3K4me2 + H3K27me3 | YES | YES | <=1 TM | 110 | 6 | YES | Uncharacterized protein GT037_006531 [*Alternaria burnsii*] Hypothetical protein CC77DRAFT_436293 [*Alternaria alternata*] Uncharacterized protein ALTATR162_LOCUS920 [*Alternaria atra*] Hypothetical protein AA0120_g5702 [*Alternaria tenuissima*] Uncharacterized protein J4E93_007105 [*Alternaria ventricosa*] | Good | None |
| *Lmb_jn3_12475* | - | Cluster 2 | GC | YES | DTM_Sahana (800bp) | H3K9me3 + H3K27me3 | YES | YES | <=1 TM | 158 | 3 | YES | Hypothetical protein J1614_003640 [*Plenodomus biglobosus*] Hypothetical protein T440DRAFT_456238 [*Plenodomus tracheiphilus IPT5*] | Good | SC_2L4HC2_23 Methyl-Accepting Chemotaxis Transducer Periplasmic binding protein/ LACI Transcriptional Neoleukin-2/15 |

| | | | | | | | | | | | | | | | |
|---|---|---|---|---|---|---|---|---|---|---|---|---|---|---|---|
| | | | | | | | | | | | | | Hypothetical protein EKO04_002699 [*Ascochyta lentis*]<br>Hypothetical protein EJ07DRAFT_162498 [*Lizonia empirigonia*]<br>Uncharacterized protein EKO05_0010924 [*Ascochyta rabiei*] | | Periplasmic sensor hybrid histidine kinase |
| *Lmb_jn3_12622* | - | Cluster 2 | Border | YES | RLx_Jolly (50bp)<br>RLx_Jolly (200bp)<br>RLG_Olly (500bp) | H3K27me3 | YES | YES | <=1 TM | 85 | 8 | NO | | Low | |
| *Lmb_jn3_12782* | - | Cluster 6 | GC | YES | DTM_Sahana (2100bp) | H3K27me3 | NO | YES | <=1 TM | 94 | 4 | NO | | Low | |
| *Lmb_jn3_12946* | - | Cluster 1 | GC | NO | | H3K4me2 + H3K27me3 | YES | YES | <=1 TM | 248 | 9 | YES | Hypothetical protein J1614_003163 [*Plenodomus biglobosus*]<br>Hypothetical protein T440DRAFT_394642 [*Plenodomus tracheiphilus IPT5*]<br>Hypothetical protein NX059_010839 [*Plenodomus lindquistii*]<br>Hypothetical protein BKA66DRAFT_573224 [*Pyrenochaeta* sp. MPI-SDFR-AT-0127]<br>Uncharacterized protein K460DRAFT_52514 [*Cucurbitaria berberidis* CBS 394.84] | Good | Elicitor Protein HRIP2<br>Preprothaumatin I<br>Thaumatin I<br>Protein N2P4<br>Thaumatin-2 |
| *Lmb_jn3_12986* | - | Cluster 2 | AT | YES | Lmac_Grouper_2395_3 (Overlapping)<br>RLG_Olly (200bp)<br>Lmac_Grouper_1227_4 (250bp)<br>LmTelo2 (400bp)<br>DTM_Lenwe (1100bp) | H3K9me3 | YES | YES | <=1 TM | 144 | 8 | NO | | Low | |

| | | | | | | | | | | | | | | | |
|---|---|---|---|---|---|---|---|---|---|---|---|---|---|---|---|
| | | | | | DTM_Lenwe (1300bp) DTM_Ingwe (1500bp) RLG_Polly (1800bp) RLC_Pholy (2400bp) | | | | | | | | | | |
| *Lmb_jn3_12994* | *AvrLm11* (Balesdent et al., 2013) | Cluster 2 | AT | YES | RLG_Rolly (Overlapping) Lmac_Recon_83_20 (100bp) RLG_Olly (200bp) DTM_Ingwe (500bp) RLG_Polly (800bp) RLG_Olly (1000bp) RLC_Pholy (2200bp) | H3K9me3 | YES | YES | <=1 TM | 105 | 10 | YES | Uncharacterized protein MYCFIDRAFT_212012 [*Pseudocercospora fijiensis* CIRAD86] | Good | None |
| *Lmb_jn3_13126* | *AvrLm1* (Gout et al., 2006) | Cluster 2 | AT | YES | RLG_Olly (150bp) DTT_Finwe-1 (1000bp) DTx_Olwe (1500bp) Lmac_Grouper_595_20 (1700bp) LmTelo2 (2500bp) RLx_Ayoly (3100bp) RLC_Pholy (3600bp) RLG_Olly (3900bp) | H3K9me3 | YES | YES | <=1 TM | 206 | 1 | YES | Hypothetical protein FANTH_13290 [*Fusarium anthophilum*] Hypothetical protein FDENT_13598 [*Fusarium denticulatum*] Hypothetical protein F52700_6206 [*Fusarium* sp. NRRL 52700] Hypothetical protein FMEXI_10338 [*Fusarium mexicanum*] Hypothetical protein BFJ63_vAg17718 [*Fusarium oxysporum* f. sp. *narcissi*) | Good | None |

[a] From JN3 annotation (Dutreux et al., 2018)

[b] Data from Gay et al. (2021)

[c] From JN3 annotation (Dutreux et al., 2018): Border = 10kb interval between AT-rich and GC-equilibrated isochores

[d] From JN3 TE annotation (Grandaubert et al., 2014)

[e] Data from Soyer et al. (2021)

[f] Predicted by SignalP (Nielsen, 2017)

[g] Predited by TargetP (Almagro Armenteros et al., 2019)

[h] Predicted by TMHMM tool (Möller et al., 2001)

[i] Blastp outside *L. maculans*, from NCBI nr database :
https://blast.ncbi.nlm.nih.gov/Blast.cgi?PROGRAM=blastp&PAGE_TYPE=BlastSearch&LINK_LOC=blasthome

[j] Prediction by Alphafold2 (ColabFold v1.5.5, Mirdita et al., 2022)

[k] Dali server, Z > 5, RSMD < 3

Table S2: Description of 147 effector gene conservation in the *Leptosphaeria maculans* IBCN collection described by Van de Wouw *et al.* (2024). Genomes of 205 isolates from the new IBCN collection were analyzed for sequence variation content relative to the reference strain JN3 using variant calling method.

| Gene ID [a] | Number of isolates presenting the gene | Alignment length | Number of isoforms | Number of isolates with polymorphim [b] | Isoforms description | Number of isolates per isoform | Assignation to the reference expression clusters [c] |
|---|---|---|---|---|---|---|---|
| *Lmb_jn3_00001* | 197 | 160 | 18 | 65 | I $^{8}$V | 2 | Cluster 2 |
| | | | | | A $^{17}$S | 1 | |
| | | | | | S $^{51}$N , L $^{78}$F, I $^{85}$L, H $^{105}$Y and P $^{133}$T | 7 | |
| | | | | | S $^{51}$N, I $^{58}$L, L $^{78}$F, I $^{85}$L, H $^{105}$Y and P $^{133}$T | 11 | |
| | | | | | Deletion D $^{153}$ | 1 | |
| | | | | | Deletion P $^{133}$ to Q $^{160}$ | 1 | |
| | | | | | R $^{147}$S | 1 | |
| | | | | | I $^{135}$K | 1 | |
| | | | | | D $^{35}$E | 9 | |
| | | | | | S $^{51}$N | 16 | |
| | | | | | D $^{35}$E and S $^{51}$N | 1 | |
| | | | | | S $^{51}$N, I $^{58}$H, H $^{105}$Y, G $^{131}$R and F $^{134}$Y | 5 | |
| | | | | | I $^{8}$F, S $^{51}$N, I $^{58}$H, H $^{105}$Y, G $^{131}$R, P $^{133}$T and F $^{134}$Y | 2 | |
| | | | | | A $^{17}$E, S $^{51}$N, I $^{58}$H, H $^{105}$Y, G $^{131}$R, P $^{133}$T and F $^{134}$Y | 1 | |
| | | | | | S $^{51}$N, I $^{58}$H, H $^{105}$Y, G $^{131}$R, P $^{133}$T and F $^{134}$Y | 3 | |
| | | | | | C $^{23}$G, S $^{51}$N, I $^{58}$H, H $^{105}$Y, G $^{131}$R, P $^{133}$T and F $^{134}$Y | 1 | |
| | | | | | S $^{51}$N and L $^{78}$F | 1 | |
| | | | | | D $^{35}$E and S $^{51}$N | 1 | |
| *Lmb_jn3_00187* | 203 | 102 | 1 | 14 | N $^{72}$K | 14 | Cluster 2 |
| *Lmb_jn3_00284* | 201 | 117 | 0 | 0 | | | Cluster 4 |
| *Lmb_jn3_00461* | 201 | 147 | 2 | 2 | I $^{6}$M | 1 | Cluster 2 |
| | | | | | D $^{48}$E | 1 | |
| *Lmb_jn3_00495* | 200 | 123 | 3 | 3 | A $^{5}$E | 1 | Cluster 4 |
| | | | | | G $^{43}$S | 1 | |
| | | | | | Deletion Q$^{76}$ to P$^{123}$ | 1 | |
| *Lmb_jn3_00561* | 201 | 218 | 1 | 1 | Deletion M$^{1}$ to T$^{39}$ | 1 | Cluster 6 |
| *Lmb_jn3_00617* | 201 | 323 | 0 | 0 | | | Cluster 6 |
| *Lmb_jn3_00630* | 200 | 565 | 12 | 178 | Insertion N$^{351}$ to N$^{359}$, insertion G$^{379}$ to A$^{393}$, insertion A$^{408}$ to G$^{419}$, N $^{377}$D and N $^{378}$A | 58 | Cluster 5 |
| | | | | | Insertion G$^{379}$ to A$^{393}$, insertion A$^{408}$ to G$^{419}$, N $^{377}$D and N $^{378}$A | 27 | |
| | | | | | Deletion G$^{168}$ to A$^{565}$ | 1 | |

| | | | | | | | |
|---|---|---|---|---|---|---|---|
| | | | | | Deletion $N^{338}$ to $G^{350}$, ,insertion $G^{379}$ to $A^{393}$, insertion $A^{408}$ to $G^{419}$, G $^{313}$S, N $^{377}$D and N $^{378}$A | 14 | |
| | | | | | Insertion $N^{212}$ to $G^{229}$, ,insertion $G^{379}$ to $A^{393}$, insertion $A^{408}$ to $G^{419}$, G $^{313}$S, N $^{377}$D and N $^{378}$A | 1 | |
| | | | | | Insertion $N^{212}$ to $G^{229}$, ,insertion $G^{379}$ to $A^{393}$, insertion $A^{408}$ to $G^{419}$, N $^{377}$D and N $^{378}$A | 37 | |
| | | | | | Deletion $N^{240}$ to $N^{249}$ | 1 | |
| | | | | | Deletion $N^{373}$ to $N^{378}$ and deletion $D^{394}$ to $G^{396}$ | 1 | |
| | | | | | Insertion $N^{212}$ to $G^{229}$ and deletion $N^{360}$ to $N^{368}$ | 1 | |
| | | | | | Insertion $N^{212}$ to $G^{229}$ and Y $^{14}$H | 1 | |
| | | | | | Insertion $N^{212}$ to $G^{229}$ | 35 | |
| | | | | | Deletion $N^{360}$ to $N^{368}$ | 1 | |
| *Lmb_jn3_00833* | 201 | 98 | 1 | 1 | K $^{33}$N | 1 | Cluster 4 |
| *Lmb_jn3_00834* | 201 | 60 | 0 | 0 | | | Cluster 4 |
| *Lmb_jn3_00910* | 202 | 216 | 1 | 1 | A $^{15}$S | 1 | Cluster 6 |
| *Lmb_jn3_00912* | 201 | 191 | 0 | 0 | | | Cluster 6 |
| *Lmb_jn3_00919* | 201 | 119 | 0 | 0 | | | Cluster 2 |
| *Lmb_jn3_00968* | 138 | 101 | 2 | 2 | Deletion $C^{101}$ and K $^{98}$N | 1 | Cluster 4 |
| | | | | | Deletion $M^{1}$ to $G^{83}$ | 1 | |
| *Lmb_jn3_01236* | 201 | 176 | 2 | 4 | F $^{9}$V | 1 | Cluster 5 |
| | | | | | T $^{150}$A | 3 | |
| *Lmb_jn3_01238* | 201 | 102 | 2 | 4 | L $^{13}$V | 3 | Cluster 5 |
| | | | | | V $^{70}$I | 1 | |
| *Lmb_jn3_01277* | 201 | 108 | 1 | 1 | C $^{74}$Y | 1 | Cluster 5 |
| *Lmb_jn3_01426* | 201 | 139 | 1 | 2 | Q $^{59}$R | 2 | Cluster 4 |
| *Lmb_jn3_01427* | 201 | 137 | 1 | 1 | N $^{133}$D | 1 | Cluster 4 |
| *Lmb_jn3_01428* | 201 | 130 | 1 | 1 | H $^{12}$Y | 1 | Cluster 4 |
| *Lmb_jn3_01748* | 199 | 129 | 6 | 61 | A $^{8}$V | 1 | Cluster 2 |
| | | | | | G $^{68}$D and E $^{73}$D | 2 | |
| | | | | | G $^{68}$D | 1 | |
| | | | | | P $^{19}$A, A $^{27}$V, Q $^{62}$K, G $^{68}$D, N $^{70}$D and S $^{77}$Y | 1 | |
| | | | | | P $^{19}$A, A $^{27}$V, Q $^{62}$K, G $^{68}$D and S $^{77}$Y | 52 | |
| | | | | | Q $^{62}$K, G $^{68}$D and S $^{77}$Y | 4 | |
| *Lmb_jn3_01779* | 201 | 57 | 0 | 0 | | | Cluster 5 |
| *Lmb_jn3_01849* | 201 | 102 | 1 | 1 | G $^{25}$D | 1 | Cluster 4 |
| *Lmb_jn3_01851* | 201 | 91 | 3 | 3 | S $^{15}$F | 1 | 2 |

| | | | | | | | |
|---|---|---|---|---|---|---|---|
| | | | | | H $^{22}$D | 1 | |
| | | | | | D $^{64}$H | 1 | |
| *Lmb_jn3_01852* | 201 | 88 | 0 | 0 | | | Cluster 5 |
| *Lmb_jn3_01853* | 202 | 109 | 0 | 0 | | | Cluster 4 |
| *Lmb_jn3_02093* | 200 | 144 | 1 | 49 | R $^{119}$S | 49 | Cluster 2 |
| *Lmb_jn3_02094* | 201 | 107 | 4 | 6 | L $^{3}$S | 3 | Cluster 2 |
| | | | | | V $^{8}$I | 1 | |
| | | | | | L $^{11}$I | 1 | |
| | | | | | S $^{84}$L | 1 | |
| *Lmb_jn3_02187* | 201 | 106 | 5 | 7 | T $^{53}$I | 1 | Cluster 4 |
| | | | | | Deletion Q$^{9}$ and P $^{31}$L | 2 | |
| | | | | | G $^{72}$R | 1 | |
| | | | | | Deletion D$^{68}$ to K$^{106}$ | 1 | |
| | | | | | Highly divergent | 2 | |
| *Lmb_jn3_02342* | 201 | 257 | 0 | 0 | | | Cluster 4 |
| *Lmb_jn3_02354* | 201 | 139 | 6 | 123 | D $^{32}$N, I $^{92}$L and T $^{97}$S | 3 | Cluster 4 |
| | | | | | D $^{32}$N and I $^{92}$L | 33 | |
| | | | | | I $^{92}$L | 48 | |
| | | | | | D $^{32}$N, Y $^{35}$C, R $^{36}$Q, I $^{92}$L and T $^{97}$S | 28 | |
| | | | | | Deletion E$^{123}$ | 4 | |
| | | | | | D $^{32}$N, Y $^{35}$C and R $^{36}$Q | 7 | |
| *Lmb_jn3_02425* | 200 | 131 | 2 | 25 | V $^{70}$A | 23 | Cluster 2 |
| | | | | | D $^{76}$E | 2 | |
| *Lmb_jn3_02517* | 201 | 117 | 0 | 0 | | | Cluster 4 |
| *Lmb_jn3_02612* | 201 | 156 | 0 | 0 | | | Cluster 5 |
| *Lmb_jn3_02967* | 200 | 74 | 3 | 102 | N $^{20}$D | 1 | Cluster 2 |
| | | | | | T $^{38}$Y, K $^{39}$N, Y $^{41}$N, V $^{44}$I, F $^{50}$Y, D $^{51}$N, V $^{65}$I and P $^{67}$S | 87 | |
| | | | | | T $^{38}$Y, K $^{39}$N, Y $^{41}$N, V $^{44}$I, F $^{50}$Y and D $^{51}$N | 14 | |
| *Lmb_jn3_03177* | 201 | 80 | 0 | 0 | | | Cluster 5 |
| *Lmb_jn3_03235* | 201 | 133 | 1 | 2 | P $^{71}$A | 2 | Cluster 4 |
| *Lmb_jn3_03238* | 70 | 94 | 8 | 68 | Deletion Y$^{85}$ to M$^{94}$, A $^{13}$T and A $^{14}$T | 1 | Cluster 2 |
| | | | | | Deletion A$^{13}$, deletion Y$^{85}$ and L $^{74}$S | 1 | |
| | | | | | Deletion A$^{13}$ and L $^{74}$S | 14 | |
| | | | | | L $^{74}$S | 37 | |
| | | | | | S $^{61}$G and L $^{74}$S | 12 | |

| | | | | | | | |
|---|---|---|---|---|---|---|---|
| | | | | | S $^{61}$G, L $^{74}$S and G $^{78}$R | 1 | |
| | | | | | P $^{53}$L | 1 | |
| | | | | | Deletion H$^{81}$, C $^{27}$Y, T $^{35}$I, D $^{49}$N, P $^{53}$L, S $^{61}$G, D $^{62}$N, G $^{63}$S, E $^{70}$K, G $^{78}$R, Q $^{82}$Y, C $^{83}$Y, H $^{84}$Y, G $^{90}$S and C $^{93}$Y | 1 | |
| *Lmb_jn3_03262* | 186 | 143 | 26 | 144 | K $^{41}$N | 10 | Cluster 2 |
| | | | | | R $^{32}$S | 2 | |
| | | | | | D $^{86}$N and G $^{120}$R | 66 | |
| | | | | | D $^{86}$N, G $^{120}$R and Q $^{121}$K | 3 | |
| | | | | | L $^{45}$S and G $^{120}$R | 9 | |
| | | | | | Q $^{35}$R, D $^{86}$N and G $^{120}$R | 1 | |
| | | | | | A $^{77}$V and G $^{120}$R | 1 | |
| | | | | | V $^{74}$I and G $^{120}$R | 1 | |
| | | | | | I $^{80}$T and G $^{120}$R | 29 | |
| | | | | | I $^{80}$T, W $^{85}$L and G $^{120}$R | 2 | |
| | | | | | Deletion Q$^{35}$, deletion R$^{64}$, deletion Q$^{75}$, deletion R$^{100}$, deletion W$^{131}$, E $^{6}$K, G $^{29}$R, R $^{32}$C, D $^{47}$N, Q $^{63}$R, M $^{70}$I, C $^{79}$Y, I $^{80}$T, D $^{83}$N, S $^{113}$L, G $^{120}$R, H $^{127}$Y, D $^{128}$N, M $^{135}$I, R $^{140}$C and D $^{143}$N | 1 | |
| | | | | | Deletion S$^{26}$ to D$^{143}$ | 1 | |
| | | | | | Deletion Q$^{35}$, deletion R$^{64}$ and A$^{65}$, deletion W$^{85}$ , deletion I$^{94}$ to D$^{143}$, G $^{29}$R, E $^{43}$K, D $^{47}$N, D $^{57}$N, Q $^{63}$R, W $^{66}$A, D $^{83}$N and D $^{86}$N | 1 | |
| | | | | | Deletion Q$^{35}$, deletion R$^{64}$ and A$^{65}$, deletion W$^{85}$, deletion Y1$^{08}$ to D$^{143}$, G $^{29}$R, E $^{43}$K, D $^{47}$N, D $^{57}$N, Q $^{63}$R, W $^{66}$A, D $^{83}$N, D $^{86}$N and R $^{100}$Q | 1 | |
| | | | | | Deletion Q$^{35}$, deletion Q$^{60}$, deletion R$^{64}$ and A$^{65}$, deletion C$^{76}$, deletion W$^{85}$, deletion R$^{100}$, deletion Q$^{121}$, deletion W$^{131}$, E $^{6}$K, T $^{20}$I, S $^{26}$L, G $^{29}$R, R $^{32}$C, E $^{43}$K, D $^{47}$N, D $^{57}$N, Q $^{63}$R, W $^{66}$A, Q $^{75}$C, D $^{83}$N, D $^{86}$N, S $^{113}$L, G $^{116}$R, G $^{120}$S, G $^{124}$S, H $^{127}$Y, D $^{128}$N, E $^{130}$K, M $^{135}$I, R $^{140}$C and G $^{141}$S | 2 | |
| | | | | | Deletion Q$^{35}$, deletion Q$^{60}$, deletion R$^{64}$ and A$^{65}$, deletion C$^{76}$, deletion W$^{85}$, deletion R$^{100}$, deletion Q$^{121}$, deletion W$^{131}$, E $^{6}$K, T $^{20}$I, S $^{26}$L, G $^{29}$R, E $^{43}$K, D $^{47}$N, D $^{57}$N, Q $^{63}$R, , W $^{66}$A, M $^{70}$I, Q $^{75}$C, D $^{83}$N, D $^{86}$N, H $^{97}$Y, S $^{113}$L, G $^{116}$R, G $^{120}$S, T $^{123}$N, G $^{124}$S, H $^{127}$Y, D $^{128}$N, E $^{130}$K, V $^{133}$I, M $^{135}$I, R $^{140}$C and G $^{141}$S | 1 | |
| | | | | | Deletion Q$^{35}$, deletion Q$^{60}$, deletion A$^{65}$, deletion C$^{76}$, deletion W$^{85}$, deletion R$^{100}$, deletion Q$^{121}$, deletion W$^{131}$, E $^{6}$K, T $^{20}$I, S $^{26}$L, G $^{29}$R, E $^{43}$K, D $^{47}$N, D $^{57}$N, W $^{66}$A, Q $^{75}$C, D $^{83}$N, H $^{97}$Y, V $^{103}$I, S $^{113}$L, G $^{116}$R, G $^{120}$S, G $^{124}$S, H $^{127}$Y, D $^{128}$N, E $^{130}$K, V $^{133}$I, M $^{135}$I, R $^{140}$C and G $^{141}$S | 1 | |
| | | | | | Deletion Q$^{35}$, deletion R$^{64}$ and A$^{65}$, deletion C$^{76}$, deletion W$^{85}$, deletion R$^{100}$, deletion Q$^{121}$, deletion W$^{131}$, E $^{6}$K, T $^{20}$I, S $^{26}$L, G $^{29}$R, E $^{43}$K, D $^{47}$N, T $^{52}$I, D $^{57}$N, Q $^{63}$R, W $^{66}$A, Q $^{75}$C, D $^{83}$N, H $^{97}$Y, V $^{103}$I, S $^{113}$L, G $^{116}$R, G $^{120}$S, G $^{124}$S, H $^{127}$Y, D $^{128}$N, E $^{130}$K, V $^{133}$I, M $^{135}$I, R $^{140}$C and G $^{141}$S | 1 | |
| | | | | | Deletion Q$^{35}$, deletion Q$^{60}$, deletion A$^{65}$, deletion C$^{76}$, deletion W$^{85}$, deletion R$^{100}$, deletion Q$^{121}$, deletion W$^{131}$, E $^{6}$K, T $^{20}$I, S $^{26}$L, G $^{29}$R, E $^{43}$K, D $^{47}$N, D $^{57}$N, W $^{66}$A, Q $^{75}$C, D $^{83}$N, S $^{113}$L, G $^{116}$R, G $^{120}$S, G $^{124}$S, H $^{127}$Y, D $^{128}$N, E $^{130}$K, M $^{135}$I, R $^{140}$C and G $^{141}$S | 1 | |

| | | | | | | | |
|---|---|---|---|---|---|---|---|
| | | | | | Deletion Q$^{35}$, deletion R$^{64}$ and A$^{65}$, deletion W$^{85}$, deletion R$^{100}$, deletion W$^{131}$, deletion C$^{142}$ and C$^{143}$, E $^{6}$K, R $^{32}$S, E $^{43}$K, D $^{47}$N, D $^{57}$N, Q $^{63}$R, W $^{66}$A, G $^{67}$S, M $^{70}$I, C $^{79}$Y, , D $^{83}$N, V $^{87}$I, G $^{120}$S, Q $^{121}$L G $^{124}$S, D $^{128}$N, E $^{130}$K, M $^{135}$I and R $^{140}$C | 2 | |
| | | | | | Deletion Q$^{35}$, deletion Q$^{60}$, deletion R$^{64}$ and A$^{65}$, deletion C$^{76}$, deletion W$^{85}$, deletion R$^{100}$, deletion Q$^{121}$, deletion W$^{131}$, E $^{6}$K, T $^{20}$I, S $^{26}$L, G $^{29}$R, E $^{43}$K, D $^{47}$N, D $^{57}$N, Q $^{63}$R, , W $^{66}$A, M $^{70}$I, Q $^{75}$C, D $^{83}$N, , S $^{113}$L, G $^{116}$R, G $^{120}$S, G $^{124}$S, H $^{127}$Y, D $^{128}$N, E $^{130}$K, M $^{135}$I, R $^{140}$C and G $^{141}$S | 3 | |
| | | | | | Deletion Q$^{35}$, deletion Q$^{60}$, deletion R$^{64}$ and A$^{65}$, deletion C$^{76}$, deletion W$^{85}$, deletion R$^{100}$, deletion Q$^{121}$, deletion W$^{131}$, E $^{6}$K, T $^{20}$I, S $^{26}$L, G $^{29}$R, E $^{43}$K, D $^{47}$N, D $^{57}$N, Q $^{63}$R, W $^{66}$A, Q $^{75}$Y, D $^{83}$N, S $^{113}$L, G $^{116}$R, G $^{120}$S, G $^{124}$S, H $^{127}$Y, D $^{128}$N, E $^{130}$K, M $^{135}$I, R $^{140}$C and G $^{141}$S | 1 | |
| | | | | | Deletion Q$^{35}$, deletion Q$^{60}$, deletion R$^{64}$ and A$^{65}$, deletion C$^{76}$, deletion W$^{85}$, deletion R$^{100}$, deletion Q$^{121}$, deletion W$^{131}$, E $^{6}$K, T $^{20}$I, S $^{26}$L, G $^{29}$R, E $^{43}$K, D $^{47}$N, D $^{57}$N, Q $^{63}$R, , W $^{66}$A, Q $^{75}$C, D $^{83}$N, S $^{113}$L, G $^{116}$R, G $^{120}$S, G $^{124}$S, H $^{127}$Y, D $^{128}$N, E $^{130}$K, M $^{135}$I, R $^{140}$C and G $^{141}$S | 1 | |
| | | | | | Deletion Q$^{35}$, deletion Q$^{60}$, deletion R$^{64}$ and A$^{65}$, deletion W$^{85}$, deletion W$^{131}$, E $^{6}$K, T $^{20}$I, S $^{26}$L, G $^{29}$R, R $^{32}$S, E $^{43}$K, D $^{47}$N, C $^{48}$Y, D $^{57}$N, Q $^{63}$R, W $^{66}$A, M $^{70}$I,C $^{79}$Y, D $^{83}$N, R $^{100}$Q, S $^{113}$L, G $^{116}$R, G $^{120}$S, Q $^{121}$L, G $^{124}$S, H $^{127}$Y, D $^{128}$N, E $^{130}$K, M $^{135}$I,R $^{140}$C and D $^{143}$N | 1 | |
| | | | | | Deletion Q$^{35}$, deletion Q$^{60}$, deletion R$^{64}$ and A$^{65}$, deletion W$^{85}$, deletion R$^{100}$, deletion W$^{131}$, E $^{6}$K, T $^{20}$I, S $^{26}$L, G $^{29}$R, R $^{32}$S, T $^{40}$I, , E $^{43}$K, D $^{47}$N, C $^{48}$Y, D $^{57}$N, Q $^{63}$R, W $^{66}$A, M $^{70}$I,C $^{79}$Y, D $^{83}$N, S $^{113}$L, G $^{116}$R, G $^{120}$S, Q $^{121}$L, G $^{124}$S, H $^{127}$Y, D $^{128}$N, M $^{135}$I,R $^{140}$C and D $^{143}$N | 1 | |
| | | | | | Deletion Q$^{35}$, deletion Q$^{60}$, deletion R$^{64}$ and A$^{65}$, deletion C$^{76}$, deletion W$^{85}$, deletion W$^{131}$,E $^{6}$K, T $^{20}$I, S $^{26}$L, G $^{29}$R, , R $^{32}$S, T $^{40}$I, , E $^{43}$K, D $^{47}$N, C $^{48}$Y, D $^{57}$N, Q $^{63}$R, W $^{66}$A, M $^{70}$I, Q $^{75}$Y, , D $^{83}$N, R $^{100}$Q, S $^{113}$L, G $^{116}$R, G $^{120}$S, Q $^{121}$L, H $^{127}$Y, D $^{128}$N, E $^{130}$K, M $^{135}$I and R $^{140}$C | 1 | |
| *Lmb_jn3_03263* | 201 | 142 | 2 | 21 | S $^{124}$R | 19 | Cluster 2 |
| | | | | | S $^{124}$R and Q $^{126}$E | 2 | |
| *Lmb_jn3_03397* | 200 | 143 | 0 | 0 | | | Cluster 2 |
| *Lmb_jn3_03409* | 201 | 73 | 1 | 1 | Deletion E$^{24}$ to C$^{73}$ | 1 | Cluster 1 |
| *Lmb_jn3_03501* | 201 | 128 | 1 | 1 | Deletion M$^{1}$ | 1 | Cluster 2 |
| *Lmb_jn3_03604* | 202 | 85 | 0 | 0 | | | Cluster 4 |
| *Lmb_jn3_03636* | 201 | 127 | 0 | 0 | | | Cluster 3 |
| *Lmb_jn3_03815* | 193 | 140 | 10 | 184 | M $^{10}$I, N $^{96}$K and I $^{117}$T | 5 | Cluster 2 |
| | | | | | A $^{72}$V, N $^{96}$K and I $^{117}$T | 1 | |
| | | | | | N $^{96}$K and I $^{117}$T | 28 | |
| | | | | | N $^{96}$K | 9 | |
| | | | | | N $^{96}$K, Q $^{97}$E and S $^{116}$G | 5 | |
| | | | | | Deletion Q$^{30}$, N $^{96}$K and S $^{116}$G | 1 | |
| | | | | | N $^{96}$K and S $^{116}$G | 103 | |
| | | | | | L $^{39}$P, N $^{96}$K and S $^{116}$G | 16 | |

| | | | | | | | |
|---|---|---|---|---|---|---|---|
| | | | | | N $^{25}$H, L $^{39}$P, N $^{96}$K and I $^{117}$T | 15 | |
| | | | | | N $^{25}$K, N $^{96}$K and I $^{117}$T | 1 | |
| *Lmb_jn3_03821* | 201 | 246 | 0 | 0 | | | Cluster 5 |
| *Lmb_jn3_03822* | 202 | 106 | 0 | 0 | | | Cluster 5 |
| *Lmb_jn3_04095* | 201 | 124 | 0 | 0 | | | Cluster 4 |
| *Lmb_jn3_04298* | 189 | 92 | 1 | 4 | I $^{82}$N | 4 | Cluster 2 |
| *Lmb_jn3_04378* | 201 | 195 | 0 | 0 | | | Cluster 2 |
| *Lmb_jn3_04385* | 201 | 89 | 0 | 0 | | | Cluster 5 |
| *Lmb_jn3_04778* | 201 | 107 | 0 | 0 | | | Cluster 5 |
| *Lmb_jn3_05067* | 198 | 82 | 2 | 10 | M $^{74}$I | 9 | Cluster 2 |
| | | | | | F $^{7}$L | 1 | |
| *Lmb_jn3_05259* | 202 | 236 | 2 | 2 | N $^{85}$D | 1 | Cluster 5 |
| | | | | | Deletion G$^{224}$ to G$^{236}$ | 1 | |
| *Lmb_jn3_05316* | 189 | 89 | 3 | 5 | R $^{26}$G, V $^{69}$F and A $^{84}$V | 2 | Cluster 2 |
| | | | | | R $^{80}$K | 2 | |
| | | | | | T $^{9}$M, P $^{40}$T, F $^{43}$V, S $^{44}$N, I $^{45}$T, L $^{50}$S, T $^{57}$I, I $^{68}$L, V $^{69}$L, A $^{70}$V, W $^{79}$S and S $^{85}$G | 1 | |
| *Lmb_jn3_05329* | 201 | 81 | 1 | 9 | R $^{47}$P | 9 | Cluster 1 |
| *Lmb_jn3_05445* | 201 | 64 | 0 | 0 | | | Cluster 3 |
| *Lmb_jn3_05451* | 201 | 79 | 0 | 0 | | | Cluster 3 |
| *Lmb_jn3_05462* | 201 | 125 | 0 | 0 | | | Cluster 6 |
| *Lmb_jn3_05465* | 201 | 95 | 1 | 1 | G $^{50}$E | 1 | Cluster 5 |
| *Lmb_jn3_05547* | 144 | 134 | 1 | 1 | T $^{5}$M | 1 | Cluster 2 |
| *Lmb_jn3_05550* | 17 | 66 | 3 | 6 | Deletion M$^{1}$ to V$^{14}$ | 3 | Cluster 5 |
| | | | | | Deletion M$^{1}$ to A$^{18}$ | 2 | |
| | | | | | Deletion M$^{1}$ to S$^{6}$ | 1 | |
| *Lmb_jn3_05637* | 201 | 57 | 0 | 0 | | | Cluster 4 |
| *Lmb_jn3_05737* | 201 | 94 | 1 | 1 | S $^{65}$F | 1 | Cluster 4 |
| *Lmb_jn3_05738* | 201 | 128 | 0 | 0 | | | Cluster 2 |
| *Lmb_jn3_05764* | 201 | 92 | 0 | 0 | | | Cluster 2 |
| *Lmb_jn3_05765* | 201 | 83 | 0 | 0 | | | Cluster 4 |
| *Lmb_jn3_05813* | 202 | 96 | 1 | 1 | K $^{72}$R | 1 | Cluster 5 |
| *Lmb_jn3_05821* | 190 | 90 | 1 | 3 | Highly divergent | 3 | Cluster 2 |
| *Lmb_jn3_05822* | 201 | 192 | 5 | 44 | N $^{160}$K | 1 | Cluster 2 |
| | | | | | S $^{170}$G | 2 | |
| | | | | | A $^{163}$P | 1 | |

| | | | | | | | |
|---|---|---|---|---|---|---|---|
| | | | | | K $^{162}$Q | 3 | |
| | | | | | K $^{178}$N | 37 | |
| *Lmb_jn3_05985* | 201 | 71 | 2 | 10 | T $^{38}$I | 9 | Cluster 2 |
| | | | | | Deletion R$^{23}$, deletion R$^{30}$, deletion R$^{67}$ to C$^{71}$, M $^{1}$I, V $^{13}$I, V $^{14}$I, Q $^{22}$R, S $^{33}$L, D $^{39}$N, R, $^{47}$Y, G $^{51}$R, A $^{52}$V and C $^{55}$Y | 1 | |
| *Lmb_jn3_05986* | 202 | 57 | 1 | 1 | Deletion R$^{57}$, S $^{4}$L, V $^{8}$I, A $^{9}$V, M $^{16}$I, P $^{35}$L, S $^{39}$L, S $^{43}$L and G $^{53}$R | 1 | Cluster 2 |
| *Lmb_jn3_06293* | 199 | 125 | 0 | 0 | | | Cluster 3 |
| *Lmb_jn3_06826* | 201 | 86 | 0 | 0 | | | Cluster 5 |
| *Lmb_jn3_06962* | 201 | 89 | 0 | 0 | | | Cluster 2 |
| *Lmb_jn3_07074* | 200 | 118 | 0 | 0 | | | Cluster 5 |
| *Lmb_jn3_07083* | 201 | 103 | 0 | 0 | | | Cluster 3 |
| *Lmb_jn3_07098* | 201 | 253 | 1 | 1 | A $^{7}$T | 1 | Cluster 5 |
| *Lmb_jn3_07106* | 201 | 252 | 2 | 12 | S $^{217}$A | 1 | Cluster 4 |
| | | | | | I $^{9}$M | 11 | |
| *Lmb_jn3_07154* | 201 | 82 | 0 | 0 | | | Cluster 2 |
| *Lmb_jn3_07285* | 201 | 255 | 1 | 1 | A $^{47}$E | 1 | Cluster 2 |
| *Lmb_jn3_07353* | 200 | 110 | 3 | 16 | Q $^{71}$P | 3 | Cluster 4 |
| | | | | | V $^{12}$I | 12 | |
| | | | | | D $^{83}$N and E $^{108}$K | 1 | |
| *Lmb_jn3_07355* | 200 | 130 | 2 | 2 | Deletion M$^{1}$ to A$^{11}$ | 1 | Cluster 1 |
| | | | | | A $^{16}$T | 1 | |
| *Lmb_jn3_07393* | 201 | 89 | 0 | 0 | | | Cluster 1 |
| *Lmb_jn3_07512* | 202 | 91 | 2 | 183 | H $^{51}$Y | 177 | Cluster 2 |
| | | | | | D $^{45}$Y and H $^{51}$Y | 6 | |
| *Lmb_jn3_07862* | 176 | 144 | 12 | 88 | V $^{2}$M and K $^{127}$E | 51 | Cluster 2 |
| | | | | | V $^{2}$M | 27 | |
| | | | | | V $^{2}$M and Q $^{144}$H | 1 | |
| | | | | | Deletion M$^{1}$ to L$^{12}$, deletion Q$^{21}$, deletion W$^{59}$, deletion W$^{134}$, deletion W$^{139}$ to Q$^{144}$, G $^{35}$R, D $^{37}$N, R $^{40}$Q, G $^{49}$R, E $^{63}$K, P $^{66}$L, E $^{68}$K, H $^{73}$Y, G $^{77}$S, H $^{89}$Y, D $^{90}$N, D $^{91}$N, G $^{92}$S, G $^{95}$S, G $^{96}$R, S $^{118}$L, P $^{124}$L and G $^{129}$R | 1 | |
| | | | | | Deletion Q$^{21}$, deletion R$^{40}$, deletion W$^{59}$, deletion W$^{134}$, deletion W$^{139}$, deletion W$^{144}$, M $^{1}$I, H $^{23}$Y, G $^{35}$R D $^{37}$N, V $^{45}$I, G $^{49}$R, T $^{60}$I, E $^{63}$K, P $^{66}$L, H $^{73}$Y, A $^{74}$T, G $^{77}$S, T $^{80}$I, H $^{89}$Y, D $^{90}$N, G $^{92}$S, G $^{95}$S, G $^{96}$R, E $^{105}$K, H $^{106}$Y, S $^{118}$L, P $^{124}$L, and G $^{129}$R | 1 | |
| | | | | | Deletion M$^{1}$ to I$^{14}$, deletion Q$^{21}$, deletion R$^{40}$, deletion W$^{59}$, deletion W$^{139}$, deletion W$^{144}$, H $^{23}$Y, C $^{26}$Y, G $^{31}$S, D $^{34}$N, G $^{35}$R, D $^{37}$N, G $^{49}$R, V $^{53}$I, E $^{63}$K, H $^{73}$Y, G $^{77}$S, H $^{89}$Y, D $^{90}$N, D $^{91}$N, G $^{92}$S, G $^{95}$S, G $^{96}$R, H $^{106}$Y, S $^{118}$L, P $^{124}$L, G $^{129}$R and V $^{137}$I | 1 | |

| | | | | | | | |
|---|---|---|---|---|---|---|---|
| | | | | | Deletion Q$^{21}$, deletion R$^{40}$, deletion W$^{59}$, deletion Q$^{87}$, deletion W$^{134}$, deletion W$^{139}$, deletion W$^{144}$, H$^{23}$Y,E $^{29}$K, G $^{31}$S, D $^{37}$N, D $^{38}$N, V $^{45}$I, G $^{49}$R, E $^{63}$K, P $^{66}$L, H $^{73}$Y, G $^{77}$S, T $^{80}$I, D $^{90}$N, D $^{91}$N, G $^{92}$S, G $^{95}$S, C $^{109}$Y, S $^{111}$L, S $^{118}$L, P $^{124}$L, G $^{126}$R and V $^{137}$I | 1 | |
| | | | | | Deletion Q$^{21}$, deletion R$^{40}$, deletion W$^{59}$, deletion W$^{139}$, deletion W$^{144}$, V $^{2}$I, H$^{23}$Y, E $^{29}$K, , G $^{35}$R, D $^{37}$N, D $^{38}$N, V $^{45}$I, G $^{49}$R, E $^{63}$K, P $^{66}$L, E $^{68}$K, H $^{73}$Y, G $^{77}$S, D $^{90}$N, G $^{92}$S, G $^{95}$S, G $^{96}$R, M $^{99}$I, P $^{124}$L, and G $^{129}$R | 1 | |
| | | | | | Deletion Q$^{21}$, deletion R$^{40}$, deletion W$^{59}$, deletion W$^{139}$, deletion W$^{144}$, M $^{1}$I, V $^{2}$I, H$^{23}$Y, G $^{31}$S, G $^{35}$R, D $^{37}$N, D $^{38}$N, G $^{49}$R, E $^{63}$K, H $^{73}$Y, A $^{74}$T, T $^{80}$I, H $^{89}$Y, D $^{90}$N, G $^{92}$S, G $^{96}$R, S $^{111}$L, G $^{126}$R, G $^{129}$R and V $^{137}$I | 1 | |
| | | | | | Deletion Q$^{21}$, deletion R$^{40}$, , deletion Q$^{87}$, deletion W$^{134}$, deletion W$^{139}$, deletion W$^{144}$, V $^{2}$I, H$^{23}$Y, G $^{31}$S, G $^{35}$R, D $^{37}$N, G $^{49}$R, E $^{63}$K, H $^{73}$Y, A $^{74}$T, G $^{77}$S, T $^{80}$I, H $^{89}$Y, M $^{99}$I, S $^{118}$L and P $^{124}$L | 1 | |
| | | | | | Deletion Q$^{21}$, deletion W$^{59}$, deletion W$^{139}$, deletion W$^{144}$, M $^{1}$I, V $^{2}$I, H$^{23}$Y, G $^{35}$R, D $^{37}$N, R $^{40}$Q, G $^{49}$R, E $^{63}$K, E $^{68}$K, H $^{73}$Y, , A $^{74}$T, G $^{77}$S, H $^{89}$Y, G $^{92}$S, G $^{95}$S, E $^{105}$K and S $^{111}$L | 1 | |
| | | | | | Deletion Q$^{21}$, deletion W$^{59}$, deletion W$^{139}$, deletion W$^{144}$, V $^{2}$M, H$^{23}$Y, G $^{31}$S, G $^{35}$R, D $^{37}$N, G $^{49}$R, E $^{63}$K, H $^{73}$Y, G $^{77}$S, H $^{89}$Y, D $^{90}$N, D $^{91}$N, G $^{92}$S, G $^{126}$R and G $^{129}$R | 1 | |
| *Lmb_jn3_07863* | 195 | 232 | 6 | 94 | N $^{133}$H | 57 | Cluster 2 |
| | | | | | Deletion V$^{227}$ to H$^{232}$ and N $^{133}$H | 1 | |
| | | | | | Q $^{146}$E | 1 | |
| | | | | | N $^{133}$G and Q $^{146}$E | 31 | |
| | | | | | N $^{133}$H and Q $^{146}$E | 3 | |
| | | | | | Deletion Q$^{33}$, deletion Q$^{42}$, deletion Q$^{53}$ to W$^{55}$, deletion Q$^{57}$, deletion W$^{94}$, deletion Q$^{105}$, deletion Q$^{111}$, deletion R$^{137}$, deletion Q$^{140}$, deletion Q$^{146}$, deletion Q$^{160}$, deletion Q$^{162}$, deletion W$^{173}$, deletion Q$^{190}$, deletion Q$^{194}$, deletion Q$^{197}$, M $^{13}$I, S $^{17}$L, S $^{26}$L, D $^{30}$N, M $^{35}$I, S $^{46}$L, H $^{49}$Y, H $^{52}$Y, R $^{81}$C, G $^{85}$R, C $^{87}$Y, S $^{88}$L, C $^{96}$Y, D $^{98}$N, P $^{102}$L, T $^{104}$M, E $^{107}$K, R $^{113}$Q, C $^{116}$Y, R $^{117}$H, E $^{119}$K, D $^{127}$N, N $^{133}$H, P $^{148}$L, D $^{151}$N, P $^{154}$L, E $^{166}$K, G $^{181}$S, S $^{191}$L, G $^{200}$R, G $^{210}$R, M $^{216}$I, D $^{221}$N and A $^{223}$V | 1 | |
| *Lmb_jn3_07874* | 200 | 190 | 3 | 44 | M $^{92}$I | 40 | Cluster 2 |
| | | | | | Deletion F$^{169}$ and L$^{190}$ | 1 | |
| | | | | | Deletion L$^{187}$ to L$^{190}$, M $^{92}$I, L $^{183}$F, I $^{184}$N and I $^{185}$Y | 3 | |
| *Lmb_jn3_07875* | 200 | 120 | 0 | 0 | | | Cluster 2 |
| *Lmb_jn3_07895* | 175 | 161 | 2 | 6 | Deletion M$^{1}$ | 3 | Cluster 3 |
| | | | | | A $^{143}$G | 3 | |
| *Lmb_jn3_07916* | 193 | 98 | 1 | 1 | S $^{4}$C | 1 | Cluster 2 |
| *Lmb_jn3_07919* | 201 | 175 | 1 | 21 | V $^{38}$I | 21 | Cluster 5 |
| *Lmb_jn3_07939* | 201 | 161 | 0 | 0 | | | Cluster 3 |
| *Lmb_jn3_08094* | 201 | 123 | 0 | 0 | | | Cluster 4 |
| *Lmb_jn3_08095* | 201 | 171 | 1 | 1 | S $^{64}$P and N $^{143}$S | 1 | Cluster 4 |
| *Lmb_jn3_08343* | 173 | 141 | 26 | 151 | Deletion Q$^{24}$, S $^{49}$L and G $^{93}$N | 1 | Cluster 2 |
| | | | | | Deletion Q$^{24}$, H $^{29}$Y, S $^{49}$L and G $^{93}$N | 1 | |

| | | | | | | | |
|---|---|---|---|---|---|---|---|
| | | | | | S $^{49}$L and G $^{93}$N | 2 | |
| | | | | | Deletion Q$^{24}$ | 1 | |
| | | | | | G $^{93}$D and R $^{95}$Q | 35 | |
| | | | | | G $^{93}$D | 1 | |
| | | | | | Deletion Q$^{24}$ and G $^{93}$N | 6 | |
| | | | | | G $^{93}$N | 62 | |
| | | | | | Deletion Q$^{24}$, A $^{17}$T and G $^{93}$N | 2 | |
| | | | | | A $^{17}$T and G $^{93}$N | 15 | |
| | | | | | A $^{17}$I and G $^{93}$N | 1 | |
| | | | | | Deletion Q$^{24}$, A $^{17}$I and G $^{93}$N | 1 | |
| | | | | | F $^{12}$S and G $^{93}$N | 1 | |
| | | | | | M $^{1}$I and G $^{93}$N | 1 | |
| | | | | | S $^{90}$L and G $^{93}$N | 6 | |
| | | | | | S $^{90}$L, I $^{92}$M and G $^{93}$N | 1 | |
| | | | | | R $^{41}$Q and G $^{93}$N | 2 | |
| | | | | | Deletion Q$^{24}$, deletion R$^{39}$ and S$^{40}$, deletion R$^{66}$, deletion R$^{87}$, deletion W$^{105}$, deletion Q$^{119}$, R $^{41}$S, S $^{49}$L, M $^{67}$I, E $^{71}$K, S $^{82}$L, S $^{90}$L, G $^{93}$N, E $^{114}$K and M $^{115}$I | 2 | |
| | | | | | Deletion R$^{41}$, deletion Q$^{69}$, S $^{49}$L, R $^{66}$W and G $^{93}$N | 1 | |
| | | | | | Highly divergent | 4 | |
| | | | | | Deletion Q$^{24}$, deletion R$^{39}$ and S$^{40}$, deletion R$^{87}$, deletion Q$^{119}$, H $^{29}$Y, R$^{41}$S, S $^{49}$L, R $^{66}$, E $^{71}$K, S $^{82}$L, G $^{93}$N, E $^{114}$K and M $^{115}$I | 1 | |
| | | | | | Deletion R$^{39}$ and S$^{40}$, A $^{17}$V, R $^{41}$S, S $^{49}$L, R $^{66}$W, E $^{71}$K, S $^{82}$L, G $^{93}$N, G $^{94}$R, E $^{114}$K, M $^{115}$I and M $^{117}$I | 1 | |
| | | | | | Deletion R$^{39}$ and S$^{40}$, deletion R$^{87}$, deletion Q$^{119}$A $^{17}$V, C $^{28}$Y, H $^{29}$Y, R $^{41}$S, D $^{47}$N, C $^{48}$Y, S $^{49}$L, R $^{66}$W, M $^{67}$I, C $^{77}$Y, G $^{93}$N, G $^{94}$R and M $^{115}$I | 1 | |
| | | | | | Deletion S$^{40}$, deletion Q$^{119}$, R $^{41}$S,A $^{61}$T, E $^{71}$K, R $^{87}$Q, G $^{93}$N, R $^{106}$Q, M $^{115}$I and M $^{117}$I | 1 | |
| | | | | | Deletion S$^{40}$, deletion R$^{87}$, deletion Q$^{119}$, R $^{41}$S, G $^{44}$S, D $^{47}$N, , S $^{49}$L, R $^{66}$W, E $^{71}$K, G $^{81}$R, S $^{90}$L, G $^{93}$N, M $^{117}$I, G $^{124}$R and D $^{126}$N | 1 | |
| *Lmb_jn3_08356* | 201 | 148 | 0 | 0 | | | Cluster 4 |
| *Lmb_jn3_08417* | 196 | 109 | 0 | 0 | | | Cluster 2 |
| *Lmb_jn3_08418* | 201 | 138 | 1 | 1 | Deletion C$^{53}$ to C$^{138}$ | 1 | Cluster 2 |
| *Lmb_jn3_08515* | 201 | 102 | 1 | 6 | S $^{20}$L | 6 | Cluster 2 |
| *Lmb_jn3_08992* | 201 | 249 | 1 | 2 | Q $^{166}$H | 2 | Cluster 1 |
| *Lmb_jn3_09025* | 201 | 230 | 3 | 12 | P $^{205}$T | 2 | Cluster 2 |
| | | | | | S $^{156}$C | 9 | |
| | | | | | T $^{213}$M | 1 | |
| *Lmb_jn3_09276* | 201 | 41 | 1 | 1 | Deletion R$^{29}$ to G$^{41}$, S $^{22}$X, P $^{23}$T, Y $^{24}$A, T $^{25}$I, A $^{27}$S and V $^{28}$L | 1 | Cluster 2 |
| *Lmb_jn3_09383* | 201 | 255 | 2 | 96 | D $^{138}$G | 95 | Cluster 4 |

| | | | | | | | |
|---|---|---|---|---|---|---|---|
| | | | | | T $^{158}$I | 1 | |
| *Lmb_jn3_09385* | 202 | 123 | 5 | 13 | Deletion M$^{1}$ to A$^{21}$ | 1 | Cluster 4 |
| | | | | | F $^{49}$L | 1 | |
| | | | | | N $^{62}$K and E $^{86}$V | 2 | |
| | | | | | N $^{62}$K | 9 | |
| *Lmb_jn3_09539* | 201 | 263 | 1 | 1 | G $^{73}$S | 1 | Cluster 4 |
| *Lmb_jn3_09745* | 153 | 120 | 9 | 12 | N $^{106}$K | 2 | Cluster 2 |
| | | | | | Deletion A$^{98}$ to S$^{120}$ | 1 | |
| | | | | | R $^{27}$M | 2 | |
| | | | | | S $^{10}$Y | 1 | |
| | | | | | A $^{97}$E | 1 | |
| | | | | | L $^{9}$F | 1 | |
| | | | | | Deletion W$^{59}$ | 1 | |
| | | | | | A $^{5}$P | 2 | |
| | | | | | Deletion Q$^{21}$, deletion Q$^{26}$, deletion Q$^{109}$, A $^{5}$P, S $^{54}$L and S $^{74}$L | 1 | |
| *Lmb_jn3_09746* | 143 | 166 | 10 | 39 | S $^{160}$A and A $^{162}$S | 19 | Cluster 2 |
| | | | | | S $^{160}$A and A $^{162}$T | 2 | |
| | | | | | S $^{160}$A | 4 | |
| | | | | | S $^{160}$A and L $^{166}$I | 1 | |
| | | | | | E $^{116}$Q and S $^{160}$A | 4 | |
| | | | | | G $^{110}$S and S $^{160}$A | 1 | |
| | | | | | S $^{160}$T | 5 | |
| | | | | | G $^{110}$A | 1 | |
| | | | | | R $^{70}$S | 1 | |
| | | | | | D $^{77}$N, G $^{78}$R, R $^{83}$Q, E $^{92}$K and E $^{103}$K | 1 | |
| *Lmb_jn3_09838* | 201 | 87 | 1 | 5 | I $^{62}$L | 5 | Cluster 6 |
| *Lmb_jn3_09935* | 201 | 255 | 1 | 46 | V $^{45}$I | 46 | Cluster 4 |
| *Lmb_jn3_09948* | 200 | 71 | 1 | 1 | Deletion Q$^{68}$, E $^{14}$K, R $^{30}$Q, R $^{34}$Q, P $^{37}$L, H $^{46}$Y, E $^{55}$K and S $^{62}$L | 1 | Cluster 5 |
| *Lmb_jn3_10035* | 201 | 236 | 0 | 0 | | | Cluster 4 |
| *Lmb_jn3_10106* | 200 | 141 | 3 | 61 | R $^{38}$L and K $^{55}$R | 2 | Cluster 2 |
| | | | | | K $^{55}$T | 9 | |
| | | | | | K $^{55}$R | 50 | |
| *Lmb_jn3_10523* | 201 | 131 | 1 | 1 | T $^{69}$S | 1 | Cluster 4 |
| *Lmb_jn3_10604* | 201 | 89 | 0 | 0 | | | Cluster 5 |
| *Lmb_jn3_10706* | 201 | 118 | 0 | 0 | | | Cluster 2 |
| *Lmb_jn3_10725* | 201 | 73 | 2 | 11 | Deletion L$^{1}$ to A$^{24}$ | 6 | Cluster 5 |

| | | | | | | | |
|---|---|---|---|---|---|---|---|
| | | | | | A $^{29}$V, V $^{58}$M and V $^{66}$A | 5 | |
| *Lmb_jn3_10844* | 200 | 232 | 2 | 2 | G $^{100}$A | 1 | Cluster 4 |
| | | | | | D $^{105}$N | 1 | |
| *Lmb_jn3_10933* | 201 | 49 | 0 | 0 | | | Cluster 5 |
| *Lmb_jn3_11031* | 201 | 177 | 2 | 2 | R $^{139}$H | 1 | Cluster 4 |
| | | | | | Deletion K$^{167}$ to Q $^{177}$ | 1 | |
| *Lmb_jn3_11224* | 201 | 95 | 0 | 0 | | | Cluster 2 |
| *Lmb_jn3_11225* | 201 | 108 | 1 | 1 | P $^{93}$Q | 1 | Cluster 2 |
| *Lmb_jn3_11242* | 201 | 211 | 4 | 14 | Deletion N$^{207}$ to K$^{211}$ | 1 | Cluster 3 |
| | | | | | Q $^{184}$K | 2 | |
| | | | | | P $^{53}$S and Q $^{184}$N | 3 | |
| | | | | | Q $^{184}$N | 8 | |
| *Lmb_jn3_11364* | 201 | 55 | 4 | 4 | V $^{12}$I | 1 | Cluster 4 |
| | | | | | V $^{22}$I | 1 | |
| | | | | | T $^{17}$I and I $^{27}$I | 1 | |
| | | | | | Deletion C$^{31}$ to C$^{55}$ and Q $^{30}$H | 1 | |
| *Lmb_jn3_11375* | 201 | 92 | 1 | 1 | S $^{85}$L | 1 | Cluster 4 |
| *Lmb_jn3_11376* | 201 | 107 | 0 | 0 | | | Cluster 4 |
| *Lmb_jn3_11426* | 201 | 78 | 0 | 0 | | | Cluster 1 |
| *Lmb_jn3_11428* | 201 | 79 | 0 | 0 | | | Cluster 4 |
| *Lmb_jn3_11454* | 167 | 247 | 2 | 2 | I $^{80}$V | 1 | Cluster 2 |
| | | | | | P $^{28}$T | 1 | |
| *Lmb_jn3_11455* | 201 | 80 | 3 | 96 | A $^{6}$G, T $^{16}$A and T $^{21}$A | 1 | Cluster 2 |
| | | | | | T $^{16}$A, T $^{21}$A and S $^{76}$R | 1 | |
| | | | | | T $^{16}$A and T $^{21}$A | 94 | |
| *Lmb_jn3_11546* | 202 | 201 | 1 | 1 | Deletion S$^{125}$ to H$^{201}$ | 1 | Cluster 1 |
| *Lmb_jn3_11916* | 201 | 101 | 0 | 0 | | | Cluster 4 |
| *Lmb_jn3_11952* | 201 | 112 | 17 | 72 | Deletion T$^{93}$ to E$^{112}$ | 1 | Cluster 4 |
| | | | | | Deletion C$^{89}$ to E$^{112}$ | 1 | |
| | | | | | Deletion C$^{111}$ and E$^{112}$ | 1 | |
| | | | | | Deletion T$^{75}$ to E$^{112}$, F $^{70}$X, R $^{71}$A, L $^{72}$P and E $^{74}$R | 1 | |
| | | | | | Deletion G$^{94}$ to E$^{112}$ and Q $^{76}$K | 47 | |
| | | | | | L $^{79}$F | 1 | |
| | | | | | Deletion G$^{59}$ to E$^{112}$ | 1 | |
| | | | | | Deletion W$^{61}$ to E$^{112}$, G $^{59}$X and I $^{60}$L | 1 | |

| | | | | | | | |
|---|---|---|---|---|---|---|---|
| | | | | | Deletion Q$^{76}$ to E$^{112}$ | 1 | |
| | | | | | Q$^{76}$K | 8 | |
| | | | | | Deletion E$^{95}$ to E$^{112}$ and Q$^{76}$K | 2 | |
| | | | | | Deletion M$^{1}$ to H$^{18}$ and deletion T$^{75}$ to E$^{112}$ | 1 | |
| | | | | | Insertion G$^{101}$ to T$^{107}$ | 1 | |
| | | | | | Deletion V$^{92}$ to E$^{112}$ and Q$^{76}$K | 1 | |
| | | | | | Deletion G$^{94}$ to E$^{112}$, Q$^{76}$K and F$^{86}$Y | 1 | |
| | | | | | Deletion G$^{94}$ to E$^{112}$ | 2 | |
| | | | | | Deletion S$^{78}$ to E$^{112}$ and Q$^{76}$K | 1 | |
| *Lmb_jn3_11990* | 201 | 126 | 6 | 140 | Deletion K$^{122}$, deletion A$^{126}$, F$^{7}$I, G$^{14}$V, S$^{16}$G, Y$^{17}$F, M$^{19}$I, D$^{20}$E, P$^{23}$L, L$^{24}$V, Y$^{25}$S, N$^{27}$D, N$^{29}$Y, D$^{31}$L, R$^{32}$K, F$^{35}$Y, P$^{37}$L, D$^{40}$G, T$^{49}$A, K$^{50}$T, T$^{53}$K, L$^{54}$Q, V$^{56}$A, M$^{60}$I, E$^{63}$G, R$^{68}$G, G$^{69}$E, L$^{73}$V, Q$^{74}$R, A$^{76}$T, Q$^{77}$H, L$^{79}$F, G$^{80}$N, S$^{84}$T, P$^{87}$S, G$^{89}$R, K$^{97}$T, F$^{102}$T, V$^{103}$T, Q$^{105}$K, P$^{108}$L, K$^{112}$Q, D$^{114}$I, K$^{115}$N, A$^{118}$I, D$^{119}$N, G$^{120}$A and N$^{121}$D | 101 | Cluster 2 |
| | | | | | Deletion K$^{122}$, deletion A$^{126}$, F$^{7}$I, G$^{14}$V, S$^{16}$G, Y$^{17}$F, M$^{19}$I, D$^{20}$E, P$^{23}$L, L$^{24}$V, Y$^{25}$S, N$^{27}$D, N$^{29}$Y, D$^{31}$L, R$^{32}$K, F$^{35}$Y, P$^{37}$L, D$^{40}$G, T$^{49}$A, K$^{50}$T, T$^{53}$K, L$^{54}$Q, V$^{56}$A, M$^{60}$I, E$^{63}$G, R$^{68}$G, G$^{69}$E, I$^{73}$V, Q$^{74}$R, A$^{76}$T, Q$^{77}$H, L$^{79}$F, G$^{80}$N, S$^{84}$T, P$^{87}$S, G$^{89}$R, K$^{97}$T, F$^{102}$T, V$^{103}$T, Q$^{105}$K, P$^{108}$L, K$^{112}$Q, D$^{114}$I, K$^{115}$N, A$^{118}$I, D$^{119}$N and G$^{120}$A | 15 | |
| | | | | | Deletion A$^{126}$, F$^{7}$I, G$^{14}$V, S$^{16}$G, Y$^{17}$F, M$^{19}$I, D$^{20}$E, P$^{23}$L, L$^{24}$V, Y$^{25}$S, N$^{27}$D, N$^{29}$Y, D$^{31}$L, R$^{32}$K, F$^{35}$Y, P$^{37}$L, D$^{40}$G, T$^{49}$A, K$^{50}$T, T$^{53}$K, L$^{54}$Q, V$^{56}$A, M$^{60}$I, E$^{63}$G, R$^{68}$G, G$^{69}$E, L$^{73}$V, Q$^{74}$R, A$^{76}$T, Q$^{77}$H, L$^{79}$F, G$^{80}$N, S$^{84}$T, P$^{87}$S, G$^{89}$R, K$^{97}$T, F$^{102}$T, V$^{103}$T, Q$^{105}$K, P$^{108}$L, K$^{112}$Q, D$^{114}$I, K$^{115}$N, A$^{118}$I, D$^{119}$N, G$^{120}$A and K$^{122}$Q | 21 | |
| | | | | | Deletion M$^{1}$ to Y$^{17}$ | 1 | |
| | | | | | Deletion D$^{31}$ to A$^{126}$, F$^{7}$I, G$^{14}$V, S$^{16}$G, Y$^{17}$F, M$^{19}$I, D$^{20}$E, P$^{23}$L, L$^{24}$V, Y$^{25}$S, N$^{27}$D and N$^{29}$Y | 1 | |
| | | | | | Deletion K$^{122}$, deletion A$^{126}$, F$^{7}$I, G$^{14}$V, S$^{16}$G, Y$^{17}$F, M$^{19}$I, D$^{20}$E, P$^{23}$L, L$^{24}$V, Y$^{25}$S, N$^{27}$D, N$^{29}$Y, D$^{31}$L, R$^{32}$K, F$^{35}$Y, P$^{37}$L, D$^{40}$G, T$^{49}$A, K$^{50}$T, T$^{53}$K, L$^{54}$Q, V$^{56}$A, M$^{60}$I, E$^{63}$G, R$^{68}$G, G$^{69}$E, L$^{73}$V, Q$^{74}$R, A$^{76}$T, Q$^{77}$H, L$^{79}$F, G$^{80}$N, S$^{84}$T, P$^{87}$S, G$^{89}$R, K$^{97}$T, F$^{102}$T, V$^{103}$T, Q$^{105}$K, P$^{108}$L, K$^{112}$Q, D$^{114}$I, K$^{115}$N, A$^{118}$I, D$^{119}$N, G$^{120}$, N$^{121}$D and P$^{123}$T | 1 | |
| *Lmb_jn3_12121* | 201 | 105 | 11 | 63 | Deletion W$^{52}$ | 2 | Cluster 4 |
| | | | | | P$^{73}$L | 4 | |
| | | | | | N$^{99}$K | 3 | |
| | | | | | N$^{99}$I | 1 | |
| | | | | | M$^{23}$I | 1 | |
| | | | | | C$^{40}$Y | 1 | |
| | | | | | Deletion Q$^{75}$ | 40 | |
| | | | | | L$^{102}$F | 3 | |
| | | | | | P$^{42}$L | 1 | |
| | | | | | Deletion R$^{63}$ and W$^{62}$R | 5 | |
| | | | | | Deletion R$^{63}$, A$^{56}$T and W$^{62}$R | 2 | |
| *Lmb_jn3_12124* | 201 | 146 | 2 | 4 | V$^{88}$L | 1 | Cluster 2 |

| | | | | | | | |
|---|---|---|---|---|---|---|---|
| | | | | | D $^{7}$N | 3 | |
| *Lmb_jn3_12249* | 135 | 55 | 0 | 0 | | | Cluster 6 |
| *Lmb_jn3_12320* | 180 | 209 | 11 | 12 | Deletion V$^{163}$ to Q$^{209}$ | 1 | Cluster 4 |
| | | | | | Deletion G$^{160}$ to Q$^{209}$ | 1 | |
| | | | | | Deletion E$^{169}$ to Q$^{209}$ | 1 | |
| | | | | | Deletion M$^{1}$, deletion P$^{2}$, deletion Y$^{148}$ to Q$^{209}$ | 1 | |
| | | | | | Deletion G$^{192}$ to Q$^{209}$ | 1 | |
| | | | | | Deletion R$^{197}$ to Q$^{209}$ | 2 | |
| | | | | | Deletion G$^{200}$ to Q$^{209}$ | 1 | |
| | | | | | Deletion E$^{206}$ to Q$^{209}$ | 1 | |
| | | | | | Deletion M$^{1}$ to S$^{3}$ | 1 | |
| | | | | | Deletion M$^{1}$ toT$^{8}$ | 1 | |
| | | | | | Deletion M$^{1}$ to T$^{36}$ | 1 | |
| *Lmb_jn3_12368* | 201 | 109 | 3 | 60 | I $^{90}$V | 2 | Cluster 5 |
| | | | | | A $^{14}$T, K $^{24}$N and I $^{90}$V | 57 | |
| | | | | | A $^{14}$T and K $^{24}$N | 1 | |
| *Lmb_jn3_12475* | 201 | 157 | 2 | 3 | F $^{44}$S | 1 | Cluster 2 |
| | | | | | G $^{106}$R | 2 | |
| *Lmb_jn3_12622* | 198 | 96 | 3 | 39 | Highly divergent | 14 | Cluster 2 |
| | | | | | Insertion N$^{20}$ to A$^{30}$, insertion E$^{58}$, deletion G$^{41}$, deletion L$^{91}$ to D$^{96}$, K $^{2}$R, T $^{19}$I, D $^{33}$N, K $^{34}$E, A $^{36}$V, Q $^{37}$A, V $^{40}$S, I $^{42}$V, R $^{44}$G, L $^{46}$A, G $^{47}$N, T$^{49}$E, T $^{50}$A, V $^{56}$L, S $^{57}$Y, T $^{59}$K, P$^{60}$T, D $^{61}$S, K $^{62}$Q, L $^{64}$S, R $^{65}$V, L $^{66}$Y, K $^{68}$G, E $^{71}$K, L $^{72}$Q, V $^{74}$I, E $^{75}$V, V $^{76}$K, Y $^{77}$V, G $^{78}$N, L $^{79}$N, K $^{80}$H, H $^{81}$S, K $^{83}$T and I $^{87}$D | 24 | |
| | | | | | Insertion N$^{20}$ to A$^{30}$, insertion E$^{58}$, deletion G$^{41}$, deletion L$^{91}$ to D$^{96}$, K $^{2}$R, A $^{13}$D, T $^{19}$I, D $^{33}$N, K $^{34}$E, A $^{36}$V, Q $^{37}$A, V $^{40}$S, I $^{42}$V, R $^{44}$G, L $^{46}$A, G $^{47}$N, T$^{49}$E, T $^{50}$A, V $^{56}$L, S $^{57}$Y, T $^{59}$K, P$^{60}$T, D $^{61}$S, K $^{62}$Q, L $^{64}$S, R $^{65}$V, L $^{66}$Y, K $^{68}$G, E $^{71}$K, L $^{72}$Q, V $^{74}$I, E $^{75}$V, V $^{76}$K, Y $^{77}$V, G $^{78}$N, L $^{79}$N, K $^{80}$H, H $^{81}$S, K $^{83}$T and I $^{87}$D | 1 | |
| *Lmb_jn3_12782* | 201 | 93 | 2 | 2 | Deletion K$^{76}$ to F$^{93}$ | 1 | Cluster 6 |
| | | | | | L $^{48}$P | 1 | |
| *Lmb_jn3_12946* | 201 | 247 | 1 | 1 | T $^{29}$I | 1 | Cluster 1 |
| *Lmb_jn3_12986* | 199 | 143 | 7 | 29 | D $^{70}$E | 3 | Cluster 2 |
| | | | | | S $^{71}$R | 7 | |
| | | | | | S $^{71}$I | 1 | |
| | | | | | D $^{94}$N | 1 | |
| | | | | | S $^{5}$L | 15 | |
| | | | | | S $^{5}$L, D $^{44}$N, R $^{56}$W and S $^{129}$L | 1 | |
| | | | | | Deletion M$^{1}$ to L$^{11}$, deletion R$^{56}$, deletion C$^{97}$, deletion R$^{105}$, deletion Q$^{118}$, R $^{23}$W, P $^{34}$L, D $^{37}$N, G $^{38}$R, S $^{58}$L, C $^{75}$Y, A $^{78}$T, D $^{81}$N, E $^{82}$K, H $^{98}$Y, W $^{99}$Y, G $^{125}$R, S $^{129}$L and D $^{135}$N | 1 | |

| | | | | | | | |
|---|---|---|---|---|---|---|---|
| *Lmb_jn3_12994* | 189 | 104 | 4 | 38 | E $^{31}$T | 30 | Cluster 2 |
| | | | | | E $^{31}$T and K $^{34}$E | 6 | |
| | | | | | N $^{6}$S | 1 | |
| | | | | | R $^{74}$Q, E $^{89}$K and E $^{99}$K | 1 | |
| *Lmb_jn3_13126* | 34 | 205 | 2 | 3 | Deletion M$^{1}$ to Q$^{3}$, deletion Q$^{47}$, deletion Q$^{143}$, deletion Q$^{192}$, S $^{10}$L, A $^{14}$V, S $^{18}$L, P $^{24}$L, S $^{55}$L, A $^{59}$V, E $^{63}$K, H $^{108}$Y, H $^{118}$Y, E $^{119}$K, P $^{122}$L, D $^{128}$N, P $^{132}$L, E $^{151}$K, H $^{155}$Y, G $^{173}$S, A $^{174}$V, S $^{175}$L, G $^{179}$S, D $^{183}$N, T $^{186}$I, A $^{202}$V and A $^{204}$V | 1 | Cluster 2 |
| | | | | | I $^{125}$K and H $^{155}$Y | 2 | |

[a] From JN3 annotation (Dutreux et al., 2018)

[b] Number of isolates carrying a polymorphic effector compared to JN3 proteome using variant calling method

[c] Data from Gay et al. (2021)

Table S3: *Leptosphaeria maculans* isolates used in the present study. Transformants used for the first screening are indicated with an asterisk. The validations were done by inoculating with two transformants per *LmSTEE* gene.

| Isolate name | Background [a] | Avirulence genes based on phenotyping data [b] | Reference |
|---|---|---|---|
| JN2 | - | *AvrLm5, AvrLm6, AvrLm7, AvrLm8, AvrLm10, AvrLm11, AvrLmS, AvrLm14, AvrLepR1* | Balesdent et al., 2001 |
| JN3 | - | *AvrLm1, AvrLm4, AvrLm5, AvrLm6, AvrLm7, AvrLm8, AvrLm10, AvrLm11, AvrLmS, AvrLm14, AvrLepR1* | Rouxel et al., 2011 |
| INV13.269 | - | *AvrLm5, AvrLm6, AvrLm8, AvrLm10, AvrLm11, AvrLmS, AvrLm14, AvrLepR1* | Plissonneau et al., 2016 |
| X83.51 | - | *AvrLm5, AvrLm10, AvrLm14, AvrLepR1* | Present study |
| pA4-7::LmSTEE1_1* | INV13.269 | AvrLm5, AvrLm6, AvrLm8, AvrLm10, AvrLm11, AvrLmS, AvrLm14, AvrLepR1, AvrLmSTEE1 | Jiquel et al., 2021 |
| pA4-7::LmSTEE1_3 | INV13.269 | AvrLm5, AvrLm6, AvrLm8, AvrLm10, AvrLm11, AvrLmS, AvrLm14, AvrLepR1, AvrLmSTEE1 | Jiquel et al., 2021 |
| pA4-7::LmSTEE35_3 | INV13.269 | AvrLm5, AvrLm6, AvrLm8, AvrLm10, AvrLm11, AvrLmS, AvrLm14, AvrLepR1, AvrLmSTEE35 | Jiquel et al., 2021 |
| pA4-7::LmSTEE35_4* | INV13.269 | AvrLm5, AvrLm6, AvrLm8, AvrLm10, AvrLm11, AvrLmS, AvrLm14, AvrLepR1, AvrLmSTEE35 | Jiquel et al., 2021 |
| pA4-7::LmSTEE78_1* | INV13.269 | AvrLm5, AvrLm6, AvrLm8, AvrLm10, AvrLm11, AvrLmS, AvrLm14, AvrLepR1, AvrLmSTEE78 | Jiquel et al., 2021 |
| pA4-7::LmSTEE78_4 | INV13.269 | AvrLm5, AvrLm6, AvrLm8, AvrLm10, AvrLm11, AvrLmS, AvrLm14, AvrLepR1, AvrLmSTEE78 | Jiquel et al., 2021 |
| pA4-7::LmSTEE98_4* | INV13.269 | AvrLm5, AvrLm6, AvrLm8, AvrLm10, AvrLm11, AvrLmS, AvrLm14, AvrLepR1, AvrLmSTEE98 | Jiquel et al., 2021 |
| pA4-7::LmSTEE98_9 | INV13.269 | AvrLm5, AvrLm6, AvrLm8, AvrLm10, AvrLm11, AvrLmS, AvrLm14, AvrLepR1, AvrLmSTEE98 | Jiquel et al., 2021 |
| pA4-7::LmSTEE1277_1 | INV13.269 | AvrLm5, AvrLm6, AvrLm8, AvrLm10, AvrLm11, AvrLmS, AvrLm14, AvrLepR1, AvrLmSTEE1277 | Jiquel et al., 2022 |
| pA4-7::LmSTEE1277_4* | INV13.269 | AvrLm5, AvrLm6, AvrLm8, AvrLm10, AvrLm11, AvrLmS, AvrLm14, AvrLepR1, AvrLmSTEE1277 | Jiquel et al., 2022 |
| pA4-7::LmSTEE1852_4 | INV13.269 | AvrLm5, AvrLm6, AvrLm8, AvrLm10, AvrLm11, AvrLmS, AvrLm14, AvrLepR1, AvrLmSTEE1852 | Jiquel et al., 2022 |
| pA4-7::LmSTEE1852_5* | INV13.269 | AvrLm5, AvrLm6, AvrLm8, AvrLm10, AvrLm11, AvrLmS, AvrLm14, AvrLepR1, AvrLmSTEE1852 | Jiquel et al., 2022 |
| pA4-7::LmSTEE5465_2 | INV13.269 | AvrLm5, AvrLm6, AvrLm8, AvrLm10, AvrLm11, AvrLmS, AvrLm14, AvrLepR1, AvrLmSTEE5465 | Jiquel et al., 2022 |
| pA4-7::LmSTEE5465_3* | INV13.269 | AvrLm5, AvrLm6, AvrLm8, AvrLm10, AvrLm11, AvrLmS, AvrLm14, AvrLepR1, AvrLmSTEE5465 | Jiquel et al., 2022 |
| pA4-7::LmSTEE6826_2 | INV13.269 | AvrLm5, AvrLm6, AvrLm8, AvrLm10, AvrLm11, AvrLmS, AvrLm14, AvrLepR1, AvrLmSTEE6826 | Jiquel et al., 2022 |

| | | | |
|---|---|---|---|
| pA4-7::LmSTEE6826_3* | INV13.269 | AvrLm5, AvrLm6, AvrLm8, AvrLm10, AvrLm11, AvrLmS, AvrLm14, AvrLepR1, AvrLmSTEE6826 | Jiquel et al., 2022 |
| pA4-7::LmSTEE7919_3* | INV13.269 | AvrLm5, AvrLm6, AvrLm8, AvrLm10, AvrLm11, AvrLmS, AvrLm14, AvrLepR1, AvrLmSTEE7919 | Jiquel et al., 2022 |
| pA4-7::LmSTEE7919_4 | INV13.269 | AvrLm5, AvrLm6, AvrLm8, AvrLm10, AvrLm11, AvrLmS, AvrLm14, AvrLepR1, AvrLmSTEE7919 | Jiquel et al., 2022 |
| pA4-7::LmSTEE10933_2 | INV13.269 | AvrLm5, AvrLm6, AvrLm8, AvrLm10, AvrLm11, AvrLmS, AvrLm14, AvrLepR1, AvrLmSTEE10933 | Jiquel et al., 2022 |
| pA4-7::LmSTEE10933_3* | INV13.269 | AvrLm5, AvrLm6, AvrLm8, AvrLm10, AvrLm11, AvrLmS, AvrLm14, AvrLepR1, AvrLmSTEE10933 | Jiquel et al., 2022 |
| pA4-7::LmSTEE833_9* | X83.51 | AvrLm5, AvrLm10, AvrLm14, AvrLepR1, AvrLmSTEE833 | Present study |
| pA4-7::LmSTEE833_10 | X83.51 | AvrLm5, AvrLm10, AvrLm14, AvrLepR1, AvrLmSTEE833 | Present study |
| pA4-7::LmSTEE1853_8 | X83.51 | AvrLm5, AvrLm10, AvrLm14, AvrLepR1, AvrLmSTEE1853 | Present study |
| pA4-7::LmSTEE1853_24* | X83.51 | AvrLm5, AvrLm10, AvrLm14, AvrLepR1, AvrLmSTEE1853 | Present study |
| pA4-7::LmSTEE2187_4 | X83.51 | AvrLm5, AvrLm10, AvrLm14, AvrLepR1, AvrLmSTEE2187 | Present study |
| pA4-7::LmSTEE2187_18* | X83.51 | AvrLm5, AvrLm10, AvrLm14, AvrLepR1, AvrLmSTEE2187 | Present study |
| pA4-7::LmSTEE4385_10 | X83.51 | AvrLm5, AvrLm10, AvrLm14, AvrLepR1, AvrLmSTEE4385 | Present study |
| pA4-7::LmSTEE4385_13* | X83.51 | AvrLm5, AvrLm10, AvrLm14, AvrLepR1, AvrLmSTEE4385 | Present study |
| pA4-7::LmSTEE5765_2* | X83.51 | AvrLm5, AvrLm10, AvrLm14, AvrLepR1, AvrLmSTEE5765 | Present study |
| pA4-7::LmSTEE5765_11 | X83.51 | AvrLm5, AvrLm10, AvrLm14, AvrLepR1, AvrLmSTEE5765 | Present study |
| pA4-7::LmSTEE9385_24* | X83.51 | AvrLm5, AvrLm10, AvrLm14, AvrLepR1, AvrLmSTEE9385 | Present study |
| pA4-7::LmSTEE9385_33 | X83.51 | AvrLm5, AvrLm10, AvrLm14, AvrLepR1, AvrLmSTEE9385 | Present study |
| pA4-7::LmSTEE98_8* | X83.51 | AvrLm5, AvrLm10, AvrLm14, AvrLepR1, AvrLmSTEE98 | Present study |
| pA4-7::LmSTEE98_12 | X83.51 | AvrLm5, AvrLm10, AvrLm14, AvrLepR1, AvrLmSTEE98 | Present study |
| pA4-7::LmSTEE6826_2* | X83.51 | AvrLm5, AvrLm10, AvrLm14, AvrLepR1, AvrLmSTEE6826 | Present study |
| pA4-7::LmSTEE6826_4 | X83.51 | AvrLm5, AvrLm10, AvrLm14, AvrLepR1, AvrLmSTEE6826 | Present study |

[a] Background isolates used for the obtention of Over-Expressed in Cotyledons transformants

[b] Avirulence genes present in the isolates. The presence of the avirulence gene was estimated by phenotyping method on genotypes carrying the associated *Rlm* gene.

* Isolates used for the first screening of the 207 *B. napus* genotypes. The validations were done by inoculating with both transformants per *LmSTEE* gene.

Table S4: Interactions between 207 genotypes of *B. napus* and 16 *LmSTEE* genes from *Leptosphaeria maculans*. Transformants used for the first screening are indicated with an asterisk. The validations were done by inoculating with two transformants per *LmSTEE* gene. Type of resistance response: hypersensitive response (HR), intermediate resistance (RI) corresponding to a delayed resistance phenotype and, heterogeneous resistance (RH) corresponding to an alternance of susceptible and resistant phenotypes for a genotype x *LmSTEE* interaction.

| | |
|---|---|
| Susceptibility : S | At least 75% of inoculated plants are susceptible (score >3) or the average score is >= 3.5 |
| Hypersensitive response : HR | At least 75% of inoculated plants are resistant (score <=3) or the average score is < 2,5 |
| Intermediate resistance : RI | At least 75% of inoculated plants have an intermediate rating (=3) |
| Heterogenous resistance : RH | Less than 75% of inoculated plants are susceptible/resistant and the average score is between 2.5 and 3.5. |
| (light grey) | Interactions already tested in Jiquel *et al*. (2021, 2022) |
| (dark grey) | Less than two homogeneous plants or less than three heterogeneous plants |

| | JN2 | INV13.269 | pA4-7::LmSTEE1_1* | pA4-7::LmSTEE1_3 | pA4-7::LmSTEE1277_1 | pA4-7::LmSTEE1277_4* | pA4-7::LmSTEE1852_4 | pA4-7::LmSTEE1852_5* | pA4-7::LmSTEE35_3 | pA4-7::LmSTEE35_4* | pA4-7::LmSTEE5465_2 | pA4-7::LmSTEE5465_3* | pA4-7::LmSTEE78_1* | pA4-7::LmSTEE78_4 | pA4-7::LmSTEE7919_3* | pA4-7::LmSTEE7919_4 | pA4-7::LmSTEE10933_2 | pA4-7::LmSTEE10933_3* | pA4-7::LmSTEE6826_2 |
|---|---|---|---|---|---|---|---|---|---|---|---|---|---|---|---|---|---|---|---|
| YUDAL | S | | | | | | | | | | | | | | | | | | |
| ASTRONOM | HR | | | | | | | | | | | | | | | | | | |
| BERLIOZZ | S | | | | | | | | | | | | | | | | | | |
| ES_MAMBO | S | | | | | | | | | | | | | | | | | | |
| ESC16061 | S | | | | | | | | | | | | | | | | | | |
| ESC17073 | S | | | | | | | | | | | | | | | | | | |
| INN_GMLm_RG001 | S | | | | | | | | | | | | | | | | | | |
| INN_GMLm_RG002 | S | | | | | | | | | | | | | | | | | | |
| INN_GMLm_RG003 | S | | | | | | | | | | | | | | | | | | |
| INN_GMLm_RG004 | S | | | | | | | | | | | | | | | | | | |
| INN_GMLm_RG005 | | | | | | | | | | | | | | | | | | | |
| INN_GMLm_RG006 | S | | | | | | | | | | | | | | | | | | |
| INN_GMLm_RG007 | S | | | | | | | | | | | | | | | | | | |
| INN_GMLm_RG008 | S | | | | | | | | | | | | | | | | | | |
| INN_GMLm_RG009 | | | | | | | | | | | | | | | | | | | |
| INN_GMLm_RG010 | S | | | | | | | | | | | | | | | | | | |
| INN_GMLm_RG011 | S | | | | | | | | | | | | | | | | | | |
| INN_GMLm_RG012 | S | | | | | | | | | | | | | | | | | | |
| INN_GMLm_RG013 | S | | | | | | | | | | | | | | | | | | |
| INN_GMLm_RG014 | S | | | | | | | | | | | | | | | | | | |

| | |
|---|---|
| INN_GMLm_RG015 | S |
| INN_GMLm_RG016 | S |
| INN_GMLm_RG017 | S |
| INN_GMLm_RG018 | S |
| INN_GMLm_RG019 | S |
| INN_GMLm_RG020 | S |
| INN_GMLm_RG021 | S |
| INN_GMLm_RG022 | S |
| INN_GMLm_RG023 | S |
| INN_GMLm_RG024 | S |
| INN_GMLm_RG025 | S |
| INN_GMLm_RG027 | S |
| INN_GMLm_RG028 | S |
| INN_GMLm_RG029 | S |
| INN_GMLm_RG030 | S |
| INN_GMLm_RG031 | S |
| INN_GMLm_RG032 | S |
| INN_GMLm_RG033 | S |
| INN_GMLm_RG034 | S |
| INN_GMLm_RG035 | S |
| INN_GMLm_RG036 | S |
| INN_GMLm_RG037 | S |
| INN_GMLm_RG039 | S |
| INN_GMLm_RG040 | S |
| INN_GMLm_RG041 | S |
| INN_GMLm_RG042 | S |
| INN_GMLm_RG043 | S |
| INN_GMLm_RG044 | S |
| INN_GMLm_RG045 | S |
| INN_GMLm_RG046 | S |
| INN_GMLm_RG047 | S |
| INN_GMLm_RG048 | S |
| INN_GMLm_RG049 | S |
| INN_GMLm_RG050 | S |
| INN_GMLm_RG051 | S |
| INN_GMLm_RG052 | S |
| INN_GMLm_RG053 | S |
| INN_GMLm_RG054 | S |
| INN_GMLm_RG055 | S |

| | |
|---|---|
| INN_GMLm_RG056 | S |
| INN_GMLm_RG057 | S |
| INN_GMLm_RG058 | S |
| INN_GMLm_RG059 | S |
| INN_GMLm_RG060 | S |
| INN_GMLm_RG061 | S |
| INN_GMLm_RG062 | S |
| INN_GMLm_RG063 | S |
| INN_GMLm_RG064 | S |
| INN_GMLm_RG065 | S |
| INN_GMLm_RG066 | S |
| INN_GMLm_RG067 | S |
| INN_GMLm_RG068 | S |
| INN_GMLm_RG069 | S |
| INN_GMLm_RG070 | S |
| INN_GMLm_RG071 | S |
| INN_GMLm_RG072 | S |
| INN_GMLm_RG073 | S |
| INN_GMLm_RG074 | S |
| INN_GMLm_RG075 | S |
| INN_GMLm_RG076 | S |
| INN_GMLm_RG077 | S |
| INN_GMLm_RG078 | S |
| INN_GMLm_RG079 | S |
| INN_GMLm_RG080 | HR |
| INN_GMLm_RG081 | S |
| INN_GMLm_RG082 | S |
| INN_GMLm_RG083 | S |
| INN_GMLm_RG084 | S |
| INN_GMLm_RG085 | S |
| INN_GMLm_RG086 | S |
| INN_GMLm_RG087 | S |
| INN_GMLm_RG088 | S |
| INN_GMLm_RG089 | S |
| INN_GMLm_RG090 | S |
| INN_GMLm_RG091 | S |
| INN_GMLm_RG092 | S |
| INN_GMLm_RG093 | S/HR |
| INN_GMLm_RG094 | S |

| | | | | | | | | | | | | | | | | | | | |
|---|---|---|---|---|---|---|---|---|---|---|---|---|---|---|---|---|---|---|---|
| INN_GMLm_RG095 | S | | | | | | | | | | | | | | | | | | |
| INN_GMLm_RG096 | S | | | | | | | | | | | | | | | | | | |
| INN_GMLm_RG097 | S | | | | | | | | | | | | | | | | | | |
| INN_GMLm_RG098 | S | | | | | | | | | | | | | | | | | | |
| INN_GMLm_RG099 | S | | | | | | | | | | | | | | | | | | |
| INN_GMLm_RG100 | HR | | | | | | | | | | | | | | | | | | |
| INN_GMLm_RG101 | S | | | | | | | | | | | | | | | | | | |
| INN_GMLm_RG102 | S | S | HR | S | HR | S | S | S | S | S | S | S | S | S | S | S | S | S | S |
| INN_GMLm_RG103 | S | S | S | | | S | | S | | S | | S | S | | S | | | S | |
| INN_GMLm_RG104 | S | S | S | | HR | S | S | S | S | S | S | S | S | S | S | S | S | S | S |
| INN_GMLm_RG105 | S | S | HR | S | HR | S | S | S | S | S | S | S | S | S | S | | S | S | |
| INN_GMLm_RG106 | S | S | HR | RH | RH | S | S | S | S | S | S | S | S | S | S | | S | S | |
| INN_GMLm_RG107 | S | S | S | | HR | S | S | S | S | S | S | S | S | S | S | | | S | |
| INN_GMLm_RG108 | S | S | | | HR | S | S | S | S | S | S | S | S | S | S | | S | S | |
| INN_GMLm_RG109 | S | S | S | | HR | S | S | S | S | S | S | S | S | S | S | | | S | |
| INN_GMLm_RG110 | S | S | S | | | S | | S | | S | | S | S | | S | | | S | |
| INN_GMLm_RG111 | S | S | S | | | S | | S | | S | | S | S | | S | | | S | |
| INN_GMLm_RG112 | S | S | S | | | S | | S | | S | | S | S | | S | | | S | |
| INN_GMLm_RG113 | S | S | S | S | HR | S | S | S | S | S | S | S | S | S | S | S | S | S | |
| INN_GMLm_RG114 | S | S | HR | S | HR | S | S | S | S | S | S | S | S | S | S | S | S | S | |
| INN_GMLm_RG115 | S | S | S | | HR | S | S | S | S | S | S | S | S | S | S | S | | S | S |
| INN_GMLm_RG116 | S | S | RH | S | HR | S | S | S | S | S | S | S | S | S | S | S | S | S | |
| INN_GMLm_RG117 | S | S | HR | S | HR | S | S | S | S | S | S | S | S | S | S | S | S | S | |
| INN_GMLm_RG118 | S | S | S | | HR | S | S | S | S | S | S | S | S | S | S | S | S | S | |
| INN_GMLm_RG119 | S | S | S | | HR | S | S | S | S | S | S | S | S | S | S | | S | S | |
| INN_GMLm_RG120 | S | S | S | S | HR | S | S | S | S | S | S | S | S | S | S | S | | S | |
| INN_GMLm_RG121 | S | S | S | | HR | S | | S | S | S | S | S | S | | S | S | S | S | |
| INN_GMLm_RG122 | HR | S | RH | S | HR | S | | S | S | S | S | S | S | | S | S | S | S | |
| INN_GMLm_RG123 | S | S | S | | HR | S | S | S | S | S | S | S | S | | S | S | S | S | |
| INN_GMLm_RG124 | S | S | RH | S | HR | S | S | S | S | S | S | S | S | S | S | S | | | |
| INN_GMLm_RG125 | S | S | S | | | S | | S | S | S | S | S | S | | S | | S | S | |
| INN_GMLm_RG126 | S | S | RH | S | S | S | S | S | S | S | S | S | S | | S | S | S | S | |
| INN_GMLm_RG127 | S | S | S | | RH | S | S | S | S | S | S | S | S | | S | S | | S | |
| INN_GMLm_RG128 | S | S | S | | HR | S | S | S | S | S | S | S | S | | S | S | S | S | |
| INN_GMLm_RG129 | S | S | S | | HR | S | S | S | RH | S | S | S | S | | S | S | S | S | |
| INN_GMLm_RG130 | S | S | | | | | | | | | | | | | | | | | |
| INN_GMLm_RG131 | S | S | S | | HR | S | | S | S | S | S | S | S | | S | S | S | S | |
| INN_GMLm_RG132 | S | S | S | | HR | S | | S | S | S | S | S | S | | S | S | S | S | |
| INN_GMLm_RG133 | S | S | S | | | | | S | S | S | S | | S | | S | | S | S | |

| | | | | | | | | | | | | | | | | | | | |
|---|---|---|---|---|---|---|---|---|---|---|---|---|---|---|---|---|---|---|---|
| INN_GMLm_RG134 | | | | | | | | | | | | | | | | | | | |
| INN_GMLm_RG135 | S | S | S | | HR | S | | S | S | S | S | S | S | | S | S | S | S | |
| INN_GMLm_RG136 | S | S | S | | HR | HR | | S | S | S | S | RH | S | | S | | HR | RH | |
| INN_GMLm_RG137 | S | S | S | | HR | S | | S | S | S | | S | S | | S | | | S | |
| INN_GMLm_RG138 | S | S | S | | HR | S | S | S | | S | S | S | S | | S | | | S | |
| INN_GMLm_RG139 | S | S | S | | HR | S | | S | RH | HR | S | S | S | | S | | | S | |
| INN_GMLm_RG140 | S | S | S | | HR | RH | | S | | S | S | S | S | | S | | | S | |
| INN_GMLm_RG141 | S | S | S | | RH | S | | S | | S | S | S | S | | S | | | S | |
| INN_GMLm_RG142 | S | S | S | | RH | S | | S | S | S | S | S | S | | S | | | S | |
| INN_GMLm_RG143 | S | S | S | | HR | S | S | S | S | S | S | S | S | | S | | S | S | |
| INN_GMLm_RG144 | S | S | S | S | HR | S | | S | S | S | S | S | S | | S | | S | S | |
| INN_GMLm_RG145 | S | S | S | | HR | S | S | S | | S | S | S | S | | S | | | S | |
| INN_GMLm_RG146 | S | S | S | | RH | S | | S | S | S | S | S | S | | S | | | S | |
| INN_GMLm_RG147 | S | S | S | | HR | S | S | S | S | S | S | S | S | | S | | S | S | |
| INN_GMLm_RG148 | S | S | S | | RH | S | | S | S | S | S | S | S | | S | | S | S | |
| INN_GMLm_RG149 | S | S | S | | HR | S | | S | S | S | S | S | S | | S | | S | S | |
| INN_GMLm_RG150 | S | S | S | | HR | S | S | S | S | S | S | S | S | | S | | | S | |
| INN_GMLm_RG151 | S | S | S | | HR | S | | S | RH | RH | RH | RH | S | | S | | HR | RH | |
| INN_GMLm_RG152 | S | S | S | | RH | S | | S | S | | S | S | S | | S | | | S | |
| INN_GMLm_RG153 | S | S | HR | S | | S | S | S | S | S | S | S | S | | S | S | S | S | |
| INN_GMLm_RG154 | S | S | HR | | | S | S | S | RH | S | | | | | S | | | | |
| INN_GMLm_RG155 | S | S | HR | S | HR | S | S | S | | S | S | S | S | | S | S | S | S | |
| INN_GMLm_RG156 | S | S | | | | S | S | S | S | S | S | S | S | | | | S | S | |
| INN_GMLm_RG157 | S | S | S | S | | S | S | S | S | S | S | S | S | | S | S | S | S | |
| INN_GMLm_RG158 | S | S | S | | | S | S | S | S | S | | | S | | S | | | | |
| INN_GMLm_RG159 | S | S | S | S | | S | | S | S | S | | S | S | | S | | S | S | |
| INN_GMLm_RG160 | S | S | S | S | | S | S | S | S | S | S | S | S | | S | | S | S | |
| INN_GMLm_RG161 | S | S | HR | S | | S | S | S | S | S | S | S | S | | S | | S | S | |
| INN_GMLm_RG162 | | | | | | | | | | | | | | | | | | | |
| INN_GMLm_RG163 | S | S | HR | S | HR | S | | | | | | S | S | | S | | S | S | |
| INN_GMLm_RG164 | S | S | HR | S | | S | S | S | S | S | S | S | S | | S | | S | S | |
| INN_GMLm_RG165 | | | | | | | | | | | | | | | | | | | |
| INN_GMLm_RG166 | S | S | | | | S | | | | | | S | S | | S | | | | |
| INN_GMLm_RG167 | | | | | | | | | | | | | | | | | | | |
| INN_GMLm_RG168 | S | S | S | | | S | S | | | S | | S | S | | S | | S | | S |
| INN_GMLm_RG169 | S | S | S | | HR | S | S | S | S | S | S | S | S | | S | S | | S | |
| INN_GMLm_RG170 | S | S | S | | | S | S | | | S | | S | S | | S | | S | | S |
| INN_GMLm_RG171 | S | S | S | | | S | S | | | S | | S | S | | S | | S | | S |
| INN_GMLm_RG172 | S | S | S | | | S | S | | | S | | S | S | | S | | S | | S |

| | | | | | | | | | | | | | | | | | | | |
|---|---|---|---|---|---|---|---|---|---|---|---|---|---|---|---|---|---|---|---|
| INN_GMLm_RG173 | S | S | S | | | S | | S | | S | | S | S | | S | | | S | |
| INN_GMLm_RG174 | S | S | S | | | S | | S | | S | | S | S | | S | | | S | |
| INN_GMLm_RG175 | S | S | S | | | S | | S | | S | | S | S | | S | | | S | |
| INN_GMLm_RG176 | S | S | S | | | S | | S | | S | | S | S | | S | | | S | |
| INN_GMLm_RG177 | S | S | S | | | S | S | | | S | | S | S | | S | | S | | S |
| INN_GMLm_RG178 | S | S | S | | | S | | S | | S | | S | S | | S | | | S | |
| INN_GMLm_RG179 | S | S | S | | | S | | S | | S | | S | S | | S | | | S | |
| INN_GMLm_RG180 | S | S | S | | | S | | S | | S | | S | S | | S | | | S | |
| INN_GMLm_RG181 | S | S | S | | | S | | S | | S | | S | S | | S | | | S | |
| INN_GMLm_RG182 | S | S | S | | | S | | S | | S | | S | S | | S | | | S | |
| INN_GMLm_RG183 | S | S | S | | | S | | S | | S | | S | S | | S | | | S | |
| INN_GMLm_RG184 | S | S | S | | | S | | S | | S | | S | S | | S | | | S | |
| INN_GMLm_RG185 | S | S | S | | HR | S | S | S | S | S | S | S | S | S | S | S | S | S | |
| INN_GMLm_RG186 | S | S | | | | | | | | | | | | | | | | | S |
| INN_GMLm_RG187 | S | S | S | | | S | | S | | S | | S | S | | S | | | S | |
| INN_GMLm_RG188 | S | S | HR | S | | S | S | S | | S | S | S | S | S | S | | | S | |
| INN_GMLm_RG189 | S | S | HR | S | HR | S | S | S | S | S | S | S | S | S | S | S | | S | |
| INN_GMLm_RG190 | S | S | HR | S | | S | S | S | S | S | S | S | S | S | S | S | S | S | |
| INN_GMLm_RG191 | S | S | RH | S | S | S | S | S | S | S | S | S | S | S | S | S | S | S | |
| INN_GMLm_RG192 | S | S | S | | RH | S | S | S | S | S | S | S | S | | S | S | S | S | |
| INN_GMLm_RG193 | | | | | | | | | | | | | | | | | | | |
| INN_GMLm_RG194 | S | S | S | | HR | S | S | S | S | S | S | S | S | | S | S | S | S | |
| INN_GMLm_RG195 | S | S | HR | S | RH | S | S | S | S | S | S | S | S | S | S | S | S | S | |
| INN_GMLm_RG196 | S | S | HR | S | RH | S | S | S | S | S | S | S | S | | S | S | S | S | |
| INN_GMLm_RG197 | | | | | | | | | | | | | | | | | | | |
| INN_GMLm_RG198 | S | S | | | | | | | | | | | | | | | | | |
| INN_GMLm_RG199 | HR | S | S | | | S | S | S | S | S | S | S | S | | S | S | S | S | S |
| INN_GMLm_RG200 | S | S | HR | S | RH | S | S | S | S | S | | S | S | | S | S | S | S | |
| INN_GMLm_RG202 | S | S | RH | S | HR | S | S | S | S | S | S | S | S | | S | S | S | S | |

Table S5: 207 *Brassica napus* genotypes used for the pathogenicity assays.

| Species | Genotype | Panel | Year released | Accession type [a] | Diversity group (principal component analysis)[b] | Origins | Residual heterozygosity [c] |
|---|---|---|---|---|---|---|---|
| BRASSICA_NAPUS | YUDAL | EPHICAS | 1974 | ++ | Semi-winter | KOREA | 0,008942768 |
| BRASSICA_NAPUS | ES_MAMBO | EPHICAS | 2014 | NA | Winter | FRANCE | NA |
| BRASSICA_NAPUS | ESC16061 | EPHICAS | NA | NA | Winter | FRANCE | NA |
| BRASSICA_NAPUS | ESC17073 | EPHICAS | NA | NA | Winter | FRANCE | NA |
| BRASSICA_NAPUS | ASTRONOM | EPHICAS | 2013 | 00 | Winter | FRANCE | 0,270569183 |
| BRASSICA_NAPUS | BERLIOZZ | EPHICAS | 2013 | 00 | Winter | FRANCE | 0,278616659 |
| BRASSICA_NAPUS | INN_GMLm_RG001 | EPHICAS | 1969 | ++ | Winter | NETHERLANDS | 0,039246969 |
| BRASSICA_NAPUS | INN_GMLm_RG002 | EPHICAS | 2007 | 00 | Winter | FRANCE | 0,000475964 |
| BRASSICA_NAPUS | INN_GMLm_RG003 | EPHICAS | 1982 ? | ++ | Winter | BELGIUM | 0,124229808 |
| BRASSICA_NAPUS | INN_GMLm_RG004 | EPHICAS | 1992 | 00 | Winter | GERMANY/ GREAT BRITAIN | FIXED |
| BRASSICA_NAPUS | INN_GMLm_RG005 | EPHICAS | NA | 0/+ | Winter | GREAT BRITAIN | NA |
| BRASSICA_NAPUS | INN_GMLm_RG006 | EPHICAS | 2006 | 00 | Winter | GREAT BRITAIN | 0,197856221 |
| BRASSICA_NAPUS | INN_GMLm_RG007 | EPHICAS | <1979 | ++ | Semi-winter | JAPAN | 0,005936146 |
| BRASSICA_NAPUS | INN_GMLm_RG008 | EPHICAS | <1981 | 00 | Semi-winter | KOREA | 0,006294383 |
| BRASSICA_NAPUS | INN_GMLm_RG009 | EPHICAS | <1981 | 00 | Semi-winter | JAPAN | 0,005608974 |
| BRASSICA_NAPUS | INN_GMLm_RG010 | EPHICAS | NA | 00 | Winter | GERMANY | 0,002705714 |
| BRASSICA_NAPUS | INN_GMLm_RG011 | EPHICAS | 1986? | 00 | Winter | POLAND | 0,017841197 |
| BRASSICA_NAPUS | INN_GMLm_RG012 | EPHICAS | 2005 | 00 | Winter | GERMANY | 0,033502699 |
| BRASSICA_NAPUS | INN_GMLm_RG013 | EPHICAS | NA | ++ | Winter | BELGIUM | NA |
| BRASSICA_NAPUS | INN_GMLm_RG014 | EPHICAS | NA | 00 | Semi-winter | GREAT BRITAIN | 0,008659397 |
| BRASSICA_NAPUS | INN_GMLm_RG015 | EPHICAS | NA | NA | Winter | NA | NA |
| BRASSICA_NAPUS | INN_GMLm_RG016 | EPHICAS | <1980 | ++ | Winter | POLAND | NA |
| BRASSICA_NAPUS | INN_GMLm_RG017 | EPHICAS | 2004 | 00 | Winter | FRANCE | 0,000634618 |
| BRASSICA_NAPUS | INN_GMLm_RG018 | EPHICAS | <2003 | 00 | Winter | USA | 0,053403475 |
| BRASSICA_NAPUS | INN_GMLm_RG019 | EPHICAS | <1981 | 00 | Spring | CZECHOSLOVAKIA | 0,067096993 |
| BRASSICA_CARINATA | INN_GMLm_RG020 | EPHICAS | NA | NA | Spring | CHINA ? | NA |
| BRASSICA_NAPUS | INN_GMLm_RG021 | EPHICAS | <1979 | ++ | Semi-winter | JAPAN | 0,008219178 |
| BRASSICA_NAPUS | INN_GMLm_RG022 | EPHICAS | 2000 | 00 | Winter | GREAT BRITAIN | FIXED |
| BRASSICA_NAPUS | INN_GMLm_RG023 | EPHICAS | 1985? | 00 | Winter | GREAT BRITAIN | 0,004464998 |
| BRASSICA_NAPUS | INN_GMLm_RG024 | EPHICAS | NA | 00 | Winter | NA | 0,004318618 |
| BRASSICA_NAPUS | INN_GMLm_RG025 | EPHICAS | 1973 | NA/+ | Semi-winter | PRK | NA |
| BRASSICA_NAPUS | INN_GMLm_RG026 | EPHICAS | NA | NA | Rutabaga | DANEMARK | NA |

| | | | | | | | |
|---|---|---|---|---|---|---|---|
| BRASSICA_NAPUS | INN_GMLm_RG027 | EPHICAS | <1970 | ++ | Winter | GERMANY | NA |
| BRASSICA_NAPUS | INN_GMLm_RG028 | EPHICAS | <1981 | 00 | Winter | NA | 0,042840854 |
| BRASSICA_NAPUS | INN_GMLm_RG029 | EPHICAS | 1979 | 0/+ | Winter | GERMANY | NA |
| BRASSICA_NAPUS | INN_GMLm_RG030 | EPHICAS | 1974 | 0/+ | Winter | GERMANY | NA |
| BRASSICA_NAPUS | INN_GMLm_RG031 | EPHICAS | NA | 00 | Winter | GREAT BRITAIN | 0,004291163 |
| BRASSICA_NAPUS | INN_GMLm_RG032 | EPHICAS | NA | 00 | Winter | NA | NA |
| BRASSICA_NAPUS | INN_GMLm_RG033 | EPHICAS | NA | 0/+ | Rutabaga | CANADA | 0,023734529 |
| BRASSICA_NAPUS | INN_GMLm_RG034 | EPHICAS | NA | 0/+ | Winter | POLAND | 0,00923714 |
| BRASSICA_NAPUS | INN_GMLm_RG035 | EPHICAS | 1985? | 00 | Spring | SWEDEN | NA |
| BRASSICA_NAPUS | INN_GMLm_RG036 | EPHICAS | 1973 | ++ | Semi-winter | JAPAN | NA |
| BRASSICA_NAPUS | INN_GMLm_RG037 | EPHICAS | NA | NA | Spring | FINLAND | NA |
| BRASSICA_NAPUS | INN_GMLm_RG038 | EPHICAS | NA | 00 | Winter | GERMANY | 0,054910075 |
| BRASSICA_NAPUS | INN_GMLm_RG039 | EPHICAS | NA | 00 | Winter | GREAT BRITAIN | NA |
| BRASSICA_NAPUS | INN_GMLm_RG040 | EPHICAS | 1987 | 00 | Spring | ROMANIA | 0,319097222 |
| BRASSICA_NAPUS | INN_GMLm_RG041 | EPHICAS | NA | NA | Winter | NA | NA |
| BRASSICA_NAPUS | INN_GMLm_RG042 | EPHICAS | 1986? | 00 | Winter | GERMANY | 0,019436178 |
| BRASSICA_NAPUS | INN_GMLm_RG043 | EPHICAS | 1986? | 00 | Winter | POLAND | 0,006295932 |
| BRASSICA_NAPUS | INN_GMLm_RG044 | EPHICAS | NA | 00 | Winter | SWEDEN | 0,083434175 |
| BRASSICA_NAPUS | INN_GMLm_RG045 | EPHICAS | NA | 00 | Winter | NA | 0,002065459 |
| BRASSICA_NAPUS | INN_GMLm_RG046 | EPHICAS | NA | 00 | Semi-winter | KOREA | 0,008843651 |
| BRASSICA_NAPUS | INN_GMLm_RG047 | EPHICAS | NA | 00 | Semi-winter | KOREA | 0,037779619 |
| BRASSICA_NAPUS | INN_GMLm_RG048 | EPHICAS | NA | 00 | Winter | RUSSIA | 0,003181674 |
| BRASSICA_NAPUS | INN_GMLm_RG049 | EPHICAS | NA | NA | Semi-winter | KOREA | NA |
| BRASSICA_NAPUS | INN_GMLm_RG050 | EPHICAS | NA | 00 | Winter | NA | 0,002074027 |
| BRASSICA_NAPUS | INN_GMLm_RG051 | EPHICAS | NA | 00 | Winter | NA | 0,003177125 |
| BRASSICA_NAPUS | INN_GMLm_RG052 | EPHICAS | <1985 | 00 | Winter | GERMANY | 0,012249443 |
| BRASSICA_NAPUS | INN_GMLm_RG053 | EPHICAS | 2003 | NA | Winter | FRANCE | FIXED |
| BRASSICA_NAPUS | INN_GMLm_RG054 | EPHICAS | 2002 | NA | Winter | GERMANY | FIXED |
| BRASSICA_NAPUS | INN_GMLm_RG055 | EPHICAS | 1953 | NA | Spring | GERMANY /CANADA | NA |
| BRASSICA_NAPUS | INN_GMLm_RG056 | EPHICAS | 2005 | 00 | Winter | GERMANY | 0,002219756 |
| BRASSICA_NAPUS | INN_GMLm_RG057 | EPHICAS | 1980 | 0/+ | Winter | FRANCE | NA |
| BRASSICA_NAPUS | INN_GMLm_RG058 | EPHICAS | NA | 00 | Winter | GERMANY | 0,024764339 |
| BRASSICA_NAPUS | INN_GMLm_RG059 | EPHICAS | 1985? | 0/+ | Winter | GERMANY | NA |
| BRASSICA_NAPUS | INN_GMLm_RG060 | EPHICAS | NA | 00 | Winter | GERMANY | 0,013403542 |

| | | | | | | | |
|---|---|---|---|---|---|---|---|
| BRASSICA_NAPUS | INN_GMLm_RG061 | EPHICAS | 1997 | 00 | Winter | GERMANY | 0,024316592 |
| BRASSICA_NAPUS | INN_GMLm_RG062 | EPHICAS | NA | ++ | Winter | NA | NA |
| BRASSICA_NAPUS | INN_GMLm_RG063 | EPHICAS | NA | NA | Winter | GERMANY/NETHERLANDS | FIXED |
| BRASSICA_NAPUS | INN_GMLm_RG064 | EPHICAS | NA | 00 | Winter | GREAT BRITAIN | 0,006355259 |
| BRASSICA_NAPUS | INN_GMLm_RG065 | EPHICAS | NA | 00 | Semi-winter | JAPAN | 0,013877551 |
| BRASSICA_NAPUS | INN_GMLm_RG066 | EPHICAS | 1994 | 00 | Winter | POLAND | 0,014689622 |
| BRASSICA_NAPUS | INN_GMLm_RG067 | EPHICAS | <1980 | ++ | Winter | SWEDEN | 0,005275108 |
| BRASSICA_NAPUS | INN_GMLm_RG068 | EPHICAS | NA | NA | NA | JAPAN | NA |
| BRASSICA_NAPUS | INN_GMLm_RG069 | EPHICAS | <1982 | ++ | Winter | CZECHOSLOVAKIA | NA |
| BRASSICA_NAPUS | INN_GMLm_RG070 | EPHICAS | NA | NA | Winter | NA | NA |
| BRASSICA_NAPUS | INN_GMLm_RG071 | EPHICAS | NA | NA/+ | Spring | SWEDEN | NA |
| BRASSICA_NAPUS | INN_GMLm_RG072 | EPHICAS | <1980 | NA | Semi-winter | JAPAN | 0,02968971 |
| BRASSICA_NAPUS | INN_GMLm_RG073 | EPHICAS | NA | 00 | Winter | UKRAINE | 0,082198444 |
| BRASSICA_NAPUS | INN_GMLm_RG074 | EPHICAS | NA | NA | Semi-winter | GREAT BRITAIN | NA |
| BRASSICA_NAPUS | INN_GMLm_RG075 | EPHICAS | NA | 00 | Winter | RUSSIA | 0,290519345 |
| BRASSICA_NAPUS | INN_GMLm_RG076 | EPHICAS | NA | 00 | Winter | PAKISTAN | 0,155063291 |
| BRASSICA_NAPUS | INN_GMLm_RG077 | EPHICAS | 1996 | 00 | Winter | GERMANY | 0,010228544 |
| BRASSICA_NAPUS | INN_GMLm_RG078 | EPHICAS | 1963 | ++ | Spring | GERMANY | 0,135734072 |
| BRASSICA_NAPUS | INN_GMLm_RG079 | EPHICAS | NA | 00 | Winter | POLAND | 0,002380952 |
| BRASSICA_NAPUS | INN_GMLm_RG080 | EPHICAS | NA | 00 | Semi-winter | GREAT BRITAIN | 0,012895854 |
| BRASSICA_NAPUS | INN_GMLm_RG081 | EPHICAS | NA | ++ | Winter | GERMANY / FRANCE | NA |
| BRASSICA_NAPUS | INN_GMLm_RG082 | EPHICAS | 1996 | 00 | Winter | GERMANY | 0,003968884 |
| BRASSICA_NAPUS | INN_GMLm_RG083 | EPHICAS | NA | ++ | Winter | NA | 0,009138381 |
| BRASSICA_NAPUS | INN_GMLm_RG084 | EPHICAS | NA | NA | Winter | NA | NA |
| BRASSICA_NAPUS | INN_GMLm_RG085 | EPHICAS | NA | 00 | Winter | URSS | 0,095091528 |
| BRASSICA_NAPUS | INN_GMLm_RG086 | EPHICAS | NA | 00 | Semi-winter | NA | 0,008588076 |
| BRASSICA_NAPUS | INN_GMLm_RG087 | EPHICAS | NA | 00 | Winter | NA | 0,002074358 |
| BRASSICA_NAPUS | INN_GMLm_RG088 | EPHICAS | NA | 0/NA | Spring | GERMANY | NA |
| BRASSICA_NAPUS | INN_GMLm_RG089 | EPHICAS | <1980 | 00 | Winter | CZECHOSLOVAKIA/POLAND | 0,212406135 |
| BRASSICA_NAPUS | INN_GMLm_RG090 | EPHICAS | NA | 00 | Winter | GERMANY | 0,001111288 |
| BRASSICA_NAPUS | INN_GMLm_RG091 | EPHICAS | NA | NA | Winter | POLAND | NF |
| BRASSICA_NAPUS | INN_GMLm_RG092 | EPHICAS | 1954 | 00 | Winter | GERMANY | 0,002232855 |
| BRASSICA_NAPUS | INN_GMLm_RG093 | EPHICAS | 1999 | 0+ | NA | GERMANY | FIXED |
| BRASSICA_NAPUS | INN_GMLm_RG094 | EPHICAS | NA | NA/+ | Semi-winter | JAPAN | NA |

| | | | | | | | |
|---|---|---|---|---|---|---|---|
| BRASSICA_NAPUS | INN_GMLm_RG095 | EPHICAS | NA | 00 | Winter | DANEMARK | 0,001586294 |
| BRASSICA_NAPUS | INN_GMLm_RG096 | EPHICAS | NA | 00 | NA | SWEDEN | FIXED |
| BRASSICA_NAPUS | INN_GMLm_RG097 | EPHICAS | <1982 | ++ | Winter | CZECHOSLOVAKIA | 0,013766608 |
| BRASSICA_NAPUS | INN_GMLm_RG098 | EPHICAS | NA | ++ | Winter | POLAND | 0,002547365 |
| BRASSICA_NAPUS | INN_GMLm_RG099 | EPHICAS | NA | NA | Rutabaga | GERMANY/GREAT BRITAIN/USA | FIXED |
| BRASSICA_NAPUS | INN_GMLm_RG100 | EPHICAS | <1982 | NA | NA | CZECHOSLOVAKIA/POLAND | NA |
| BRASSICA_NAPUS | INN_GMLm_RG101 | EPHICAS | NA | 00 | Winter | NA | 0,001270245 |
| BRASSICA_NAPUS | INN_GMLm_RG102 | Semi-winter | NA | NA | Semi winter | JAPAN | NF |
| BRASSICA_NAPUS | INN_GMLm_RG104 | Semi-winter | 1970 | ++ | Semi winter | JAPAN | 0,00892133 |
| BRASSICA_NAPUS | INN_GMLm_RG105 | Semi-winter | NA | 00 | Semi winter | KOREA | 0,238053866 |
| BRASSICA_NAPUS | INN_GMLm_RG106 | Semi-winter | NA | 00 | Semi winter | KOREA | 0,060745362 |
| BRASSICA_NAPUS | INN_GMLm_RG107 | Semi-winter | NA | 00 | Semi winter | JAPAN | 0,033371882 |
| BRASSICA_NAPUS | INN_GMLm_RG108 | Semi-winter | 1962 | 00 | Semi winter | KOREA | 0,071622079 |
| BRASSICA_NAPUS | INN_GMLm_RG109 | Semi-winter | NA | 00 | Semi winter | CHINA | 0,192385787 |
| BRASSICA_NAPUS | INN_GMLm_RG110 | Semi-winter | NA | 00 | Semi winter | JAPAN | 0,336008374 |
| BRASSICA_NAPUS | INN_GMLm_RG112 | Semi-winter | 1971 | 00 | Semi winter | JAPAN | 0,291571353 |
| BRASSICA_NAPUS | INN_GMLm_RG113 | Semi-winter | NA | 00 | Semi winter | CHINA | 0,144133897 |
| BRASSICA_NAPUS | INN_GMLm_RG114 | Semi-winter | NA | 00 | Semi winter | CHINA | 0,140029207 |
| BRASSICA_NAPUS | INN_GMLm_RG115 | Semi-winter | NA | 00 | Semi winter | JAPAN | 0,280152275 |
| BRASSICA_NAPUS | INN_GMLm_RG116 | Semi-winter | NA | 00 | Semi winter | CHINA | 0,092598739 |
| BRASSICA_NAPUS | INN_GMLm_RG117 | Semi-winter | NA | NA | Semi winter | CHINA | NF |
| NAPO_BRASSICA | INN_GMLm_RG118 | Semi-winter | <1978 | 00 | Semi winter | NEW ZEALAND | 0,010143979 |
| BRASSICA_NAPUS | INN_GMLm_RG119 | Semi-winter | NA | 00 | Semi winter | KOREA | 0,006756757 |
| BRASSICA_NAPUS | INN_GMLm_RG120 | Semi-winter | 2004 | 00 | Semi winter | NEW ZEALAND | 0,166918049 |
| BRASSICA_NAPUS | INN_GMLm_RG121 | Semi-winter | NA | NA | Semi winter | NEW ZEALAND | NF |
| BRASSICA_NAPUS | INN_GMLm_RG122 | Semi-winter | NA | 00 | Semi winter | NEW ZEALAND | 0,018181818 |
| BRASSICA_NAPUS | INN_GMLm_RG123 | Semi-winter | 1973 | 00 | Semi winter | NORTH KOREA | 0,007402639 |
| BRASSICA_NAPUS | INN_GMLm_RG124 | Semi-winter | 1973 | 00 | Semi winter | JAPAN | 0,007246377 |
| BRASSICA_NAPUS | INN_GMLm_RG125 | Semi-winter | 1973 | 00 | Semi winter | JAPAN | 0,008021819 |
| BRASSICA_NAPUS | INN_GMLm_RG126 | Semi-winter | 1973 | 00 | Semi winter | JAPAN | 0,007073955 |
| BRASSICA_NAPUS | INN_GMLm_RG127 | Semi-winter | NA | 00 | Semi winter | CHINA | 0,017306331 |
| BRASSICA_NAPUS | INN_GMLm_RG128 | Semi-winter | 1973 | 00 | Semi winter | JAPAN | 0,003695373 |
| BRASSICA_NAPUS | INN_GMLm_RG129 | Semi-winter | 1971 | 00 | Semi winter | JAPAN | 0,008215206 |
| BRASSICA_NAPUS | INN_GMLm_RG131 | Semi-winter | NA | NA | Semi winter | CHINA | NF |

| | | | | | | | |
|---|---|---|---|---|---|---|---|
| BRASSICA_NAPUS | INN_GMLm_RG132 | Semi-winter | NA | 00 | Semi winter | NA | 0,007080785 |
| BRASSICA_NAPUS | INN_GMLm_RG133 | Semi-winter | NA | 00 | Semi winter | JAPAN | 0,076095812 |
| BRASSICA_NAPUS | INN_GMLm_RG135 | Semi-winter | 1981 | 00 | Semi winter | JAPAN | 0,005615274 |
| BRASSICA_NAPUS | INN_GMLm_RG136 | Semi-winter | NA | NA | Semi winter | CHINA | NF |
| BRASSICA_NAPUS | INN_GMLm_RG137 | Semi-winter | 1973 | ++ | Semi winter | JAPAN | 0,006568144 |
| BRASSICA_NAPUS | INN_GMLm_RG138 | Semi-winter | NA | NA | Semi winter | KOREA | 0,070075139 |
| BRASSICA_NAPUS | INN_GMLm_RG139 | Semi-winter | NA | 00 | Semi winter | KOREA | 0,010387924 |
| BRASSICA_NAPUS | INN_GMLm_RG140 | Semi-winter | <1979 | ++ | Semi winter | CHINA | 0,159871968 |
| BRASSICA_NAPUS | INN_GMLm_RG141 | Semi-winter | NA | 00 | Semi winter | CHINA | 0,187821612 |
| BRASSICA_NAPUS | INN_GMLm_RG142 | Semi-winter | NA | NA | Semi winter | CHINA | 0,066763896 |
| BRASSICA_NAPUS | INN_GMLm_RG143 | Semi-winter | NA | NA | Semi winter | CHINA | NF |
| BRASSICA_NAPUS | INN_GMLm_RG144 | Semi-winter | <1981 | 00 | Semi winter | KOREA | 0,244518506 |
| BRASSICA_NAPUS | INN_GMLm_RG145 | Semi-winter | <1981 | 00 | Semi winter | KOREA | 0,128514749 |
| BRASSICA_NAPUS | INN_GMLm_RG146 | Semi-winter | <1981 | 00 | Semi winter | FRANCE | 0,314483955 |
| BRASSICA_NAPUS | INN_GMLm_RG147 | Semi-winter | NA | 00 | Semi winter | KOREA | 0,273848827 |
| BRASSICA_NAPUS | INN_GMLm_RG148 | Semi-winter | NA | 00 | Semi winter | KOREA | 0,156988873 |
| BRASSICA_NAPUS | INN_GMLm_RG149 | Semi-winter | <1982 | ++ | Semi winter | JAPAN | 0,17001056 |
| BRASSICA_NAPUS | INN_GMLm_RG150 | Semi-winter | NA | 00 | Semi winter | KOREA | 0,303265666 |
| BRASSICA_NAPUS | INN_GMLm_RG151 | Semi-winter | NA | 00 | Semi winter | KOREA | 0,014098201 |
| BRASSICA_NAPUS | INN_GMLm_RG152 | Semi-winter | NA | 00 | Semi winter | KOREA | 0,246533128 |
| BRASSICA_NAPUS | INN_GMLm_RG153 | Semi-winter | NA | NA | Semi winter | KOREA | 0,185101215 |
| BRASSICA_NAPUS | INN_GMLm_RG154 | Semi-winter | NA | 00 | Semi winter | KOREA | 0,171760287 |
| BRASSICA_NAPUS | INN_GMLm_RG155 | Semi-winter | NA | 00 | Semi winter | JAPAN | 0,009242744 |
| BRASSICA_NAPUS | INN_GMLm_RG156 | Semi-winter | NA | 00 | Semi winter | CHINA | 0,008052827 |
| BRASSICA_NAPUS | INN_GMLm_RG157 | Semi-winter | <1979 | 00 | Semi winter | CHINA | 0,124687031 |
| BRASSICA_NAPUS | INN_GMLm_RG158 | Semi-winter | <1981 | 00 | Semi winter | NORTH KOREA | 0,014954486 |
| BRASSICA_NAPUS | INN_GMLm_RG159 | Semi-winter | 1973 | ++ | Semi winter | JAPAN | 0,009841884 |
| BRASSICA_NAPUS | INN_GMLm_RG160 | Semi-winter | <1981 | 00 | Semi winter | KOREA | 0,065158677 |
| BRASSICA_NAPUS | INN_GMLm_RG161 | Semi-winter | NA | 00 | Semi winter | CHINA | 0,006734007 |
| BRASSICA_NAPUS | INN_GMLm_RG163 | Semi-winter | NA | 00 | Semi winter | KOREA | 0,02897308 |
| BRASSICA_NAPUS | INN_GMLm_RG164 | Semi-winter | NA | 00 | Semi winter | TAIWAN | 0,133603788 |
| BRASSICA_NAPUS | INN_GMLm_RG166 | Semi-winter | NA | NA | Semi winter | VIETNAM | 0,008250698 |
| BRASSICA_NAPUS | INN_GMLm_RG168 | Semi-winter | NA | 00 | Semi winter | GERMANY | 0,008002561 |
| BRASSICA_NAPUS | INN_GMLm_RG169 | Semi-winter | NA | NA | Semi winter | AUSTRALIA | 0,225047321 |

| | | | | | | | |
|---|---|---|---|---|---|---|---|
| BRASSICA_NAPUS | INN_GMLm_RG170 | Semi-winter | NA | NA | Semi winter | AUSTRALIA | 0,075146218 |
| BRASSICA_NAPUS | INN_GMLm_RG171 | Semi-winter | NA | NA | Semi winter | AUSTRALIA | 0,078512936 |
| BRASSICA_NAPUS | INN_GMLm_RG172 | Semi-winter | NA | NA | Semi winter | AUSTRALIA | 0,031324338 |
| BRASSICA_NAPUS | INN_GMLm_RG173 | Semi-winter | NA | NA | Semi winter | SWEDEN | 0,011007203 |
| BRASSICA_NAPUS | INN_GMLm_RG174 | Semi-winter | NA | NA | Semi winter | AUSTRALIA | 0,066393105 |
| BRASSICA_NAPUS | INN_GMLm_RG175 | Semi-winter | NA | NA | Semi winter | JAPAN | 0,021494115 |
| BRASSICA_NAPUS | INN_GMLm_RG176 | Semi-winter | NA | 00 | Semi winter | CHINA | 0,126382836 |
| BRASSICA_NAPUS | INN_GMLm_RG177 | Semi-winter | NA | NA | Semi winter | AUSTRALIA | 0,028238993 |
| BRASSICA_NAPUS | INN_GMLm_RG178 | Semi-winter | NA | NA | Semi winter | AUSTRALIA | 0,012042599 |
| BRASSICA_NAPUS | INN_GMLm_RG179 | Semi-winter | NA | NA | Semi winter | AUSTRALIA | 0,045253095 |
| BRASSICA_NAPUS | INN_GMLm_RG180 | Semi-winter | NA | NA | Semi winter | CHINA | 0,005983263 |
| BRASSICA_NAPUS | INN_GMLm_RG181 | Semi-winter | NA | NA | Semi winter | CHINA | NA |
| BRASSICA_NAPUS | INN_GMLm_RG182 | Semi-winter | NA | NA | Semi winter | CHINA | NA |
| BRASSICA_NAPUS | INN_GMLm_RG183 | Semi-winter | NA | NA | Semi winter | CHINA | 0,008011592 |
| BRASSICA_NAPUS | INN_GMLm_RG184 | Semi-winter | NA | NA | Semi winter | NORTH KOREA | 0,007191197 |
| BRASSICA_NAPUS | INN_GMLm_RG185 | Semi-winter | NA | NA | Semi winter | NORTH KOREA | 0,007266264 |
| BRASSICA_NAPUS | INN_GMLm_RG186 | Semi-winter | NA | NA | Semi winter | CHINA | 0,01253757 |
| BRASSICA_NAPUS | INN_GMLm_RG187 | Semi-winter | NA | NA | Semi winter | CHINA | 0,009258772 |
| BRASSICA_NAPUS | INN_GMLm_RG188 | Semi-winter | 1964 | 00 | Semi winter | JAPAN | 0,009285943 |
| BRASSICA_NAPUS | INN_GMLm_RG189 | Semi-winter | 1973 | ++ | Semi winter | JAPAN | 0,075408809 |
| BRASSICA_NAPUS | INN_GMLm_RG190 | Semi-winter | <1982 | 00 | Semi winter | CHINA | 0,135498863 |
| BRASSICA_NAPUS | INN_GMLm_RG191 | Semi-winter | NA | 00 | Semi winter | NEW ZEALAND | 0,008542876 |
| BRASSICA_NAPUS | INN_GMLm_RG192 | Semi-winter | NA | 00 | Semi winter | NORTH KOREA | 0,007532051 |
| BRASSICA_NAPUS | INN_GMLm_RG193 | Semi-winter | NA | 00 | Semi winter | JAPAN | 0,217462662 |
| BRASSICA_NAPUS | INN_GMLm_RG194 | Semi-winter | 1973 | 00 | Semi winter | JAPAN | 0,008718114 |
| BRASSICA_NAPUS | INN_GMLm_RG195 | Semi-winter | <1982 | 0/+ | Semi winter | ITALIA | 0,013286377 |
| BRASSICA_NAPUS | INN_GMLm_RG196 | Semi-winter | NA | 00 | Semi winter | CHINA | 0,013301282 |
| BRASSICA_NAPUS | INN_GMLm_RG197 | Semi-winter | NA | ++ | Semi winter | VIETNAM | 0,006271298 |
| BRASSICA_NAPUS | INN_GMLm_RG198 | Semi-winter | 1960 | ++ | Semi winter | JAPAN | 0,007619493 |
| NAPO_BRASSICA | INN_GMLm_RG199 | Semi-winter | <1987 | 00 | Semi winter | USA | 0,012440661 |
| NAPO_BRASSICA | INN_GMLm_RG200 | Semi-winter | <1970 | 00 | Semi winter | DANEMARK/GREAT BRITAIN | 0,021020042 |
| BRASSICA_NAPUS | INN_GMLm_RG202 | Semi-winter | NA | NA | Semi winter | AUSTRALIA | 0,027105643 |

[a] The accession type (++, +/0, 0/+ or 00) is indicated with: " ++ ", varieties rich in erucic acid (C22:1) and glucosinolates (GSL) " +/0 ", varieties rich in C22:1 and poor in GSL " 0/+ ", varieties poor in C22:1 and rich in GSL " 00 ", varieties poor in C22:1 and in GSL

[b] See Figure 6

[c] NF = Non fixed

NA = Non available

Table S6: Primers used in the present study.

| Name | Sequence [5' – 3'] | Experiment |
| --- | --- | --- |
| Lmb_jn3_00833_EcoRI_F | GAGAGAGAATTCATGAGGTTCATCTTTCCTGC | Cloning |
| Lmb_jn3_00833_XhoI_R | GAGAGACTCGAGGCTTGCGTCAATGCAAGTTCC | Cloning |
| Lmb_jn3_01853_EcoRI_F | GAGAGAGAATTCATGAAGCTTACATTCCTTTGTTG | Cloning |
| Lmb_jn3_01853_XhoI_R | GAGAGACTCGAGGTCACGCTCCAACTTTATCCG | Cloning |
| Lmb_jn3_02187_EcoRI_F | GAGAGAGAATTCATGAACCTCCTTCTTATTGC | Cloning |
| Lmb_jn3_02187_XhoI_R | GAGAGACTCGAGGGAAGAAGTGTCTCAGCTTC | Cloning |
| Lmb_jn3_05765_EcoRI_F | GAGAGAGAATTCATGAGACTATCTATTGCACTTTC | Cloning |
| Lmb_jn3_05765_SalI_R | GAGAGAGTCGACTCAAGAACAGTTGGCGAG | Cloning |
| Lmb_jn3_00833_SalI_F | GAGAGAGTCGACGGCTTTCGGTTCAGGGTC | Cloning |
| Lmb_jn3_09385_EcoRI_F | GAGAGAGAATTCATGAAGCTAATTGTTCTCGCG | Cloning |
| Lmb_jn3_09385_XhoI_R | GAGAGACTCGAGGGGAGGAAATGTTGGCATC | Cloning |
| 04385_Gibs_F | CGAATTCGTTAACAAGCTTGTCGACATGAAGGCTATCTTTGTTTCTAC | Cloning |
| 04385_Gibs_R | CTTCTGTCGAATCGGTACCCTCGAGCTACTCGCCGGATGCAATTAGG | Cloning |
| 00833_qPCR_L2 | TGGATCTTGTACTATGGGATGGC | qRT-PCR |
| 00833_qPCR_R2 | TGTTACTCCTCTCTTCTGTTCGG | qRT-PCR |
| 01853_qPCR_L2 | TCGGCTTAGTACCTGGCTTTAC | qRT-PCR |
| 01853_qPCR_R2 | ACCAGACATTTCCGCTTTCG | qRT-PCR |
| 02187_qPCR_L2 | TGCTGCTCAGTTTTCTATGGC | qRT-PCR |
| 02187_qPCR_R2 | AAGGGTACTTCGTCGCTACTG | qRT-PCR |
| 04385_qPCR_L1 | CTCTGACAGGAGCACTAGCC | qRT-PCR |
| 04385_qPCR_R1 | GTGTTTTCTGGCAATGGACCTC | qRT-PCR |
| 05765_qPCR_L1 | GCTGAGTATGTTCGGAGCAAC | qRT-PCR |
| 05765_qPCR_R1 | TTCCGCCTTGTTTGTCACTC | qRT-PCR |
| 09385_qPCR_L1 | TTCACAGGGGGCAATATGGTC | qRT-PCR |
| 09385_qPCR_R1 | ACAGCCTTTCCTACTCAGCAC | qRT-PCR |
| LmSTEE98_qUp | ATGAAGACCTTTATCACCATTGC | qRT-PCR |
| LmSTEE98_qLow | CCGCCACAGTGAACATAAT | qRT-PCR |
| Lm6826_qUp | ACGCGTTGTAGGCTATCGTC | qRT-PCR |
| Lm6826_qLow | CATCTCACTTGCTGCGATTC | qRT-PCR |
| Actin_qUp | AGTGCGATGTCGATGTCAG | qRT-PCR |

| | | |
|---|---|---|
| Actin_qLow | AAGAGCGGTGATTTCCTTCT | qRT-PCR |
| qEF1-AR | TACCGGCGGCAATGATGAGGATA | qRT-PCR |
| qEF1-AF | ACAAATTGAAGGCCGAGCGTGAAC | qRT-PCR |
